# A germline-targeting chimpanzee SIV envelope glycoprotein elicits a new class of V2-apex directed cross-neutralizing antibodies

**DOI:** 10.1101/2022.10.18.512699

**Authors:** Frederic Bibollet-Ruche, Ronnie M. Russell, Wenge Ding, Weimin Liu, Yingying Li, Kshitij Wagh, Daniel Wrapp, Rumi Habib, Ashwin N. Skelly, Ryan S. Roark, Scott Sherrill-Mix, Shuyi Wang, Juliette Rando, Emily Lindemuth, Kendra Cruickshank, Younghoon Park, Rachel Baum, Andrew Jesse Connell, Hui Li, Elena E. Giorgi, Ge S. Song, Shilei Ding, Andrés Finzi, Amanda Newman, Giovanna E. Hernandez, Emily Machiele, Derek W. Cain, Katayoun Mansouri, Mark G. Lewis, David C. Montefiori, Kevin J. Wiehe, S. Munir Alam, I-Ting Teng, Peter D. Kwong, Raiees Andrabi, Laurent Verkoczy, Dennis R. Burton, Bette T. Korber, Kevin O. Saunders, Barton F. Haynes, Robert J. Edwards, George M. Shaw, Beatrice H. Hahn

**Affiliations:** Departments of Medicine and Microbiology, University of Pennsylvania; Philadelphia, PA 19104, USA; Theoretical Biology and Biophysics, Los Alamos National Laboratory; Los Alamos, NM 87545, USA; Duke Human Vaccine Institute, Duke University School of Medicine; Durham, NC 27710, USA; Department of Medicine, Duke University School of Medicine; Durham, NC 27710, USA; Vaccine and Immunotherapy Center, The Wistar Institute; Philadelphia, PA, 19104, USA; Department of Immunology and Microbiology, The Scripps Research Institute; La Jolla, CA 92037, USA; Centre de Recherche du CHUM; Montreal, QC H2X 0A9, Canada; Département de Microbiologie, Infectiologie et Immunologie, Université de Montréal; Montreal, QC H2X 0A9, Canada; Bioqual, Inc.; Rockville, MD 20850, USA; Department of Surgery, Duke University School of Medicine; Durham, NC, USA; Vaccine Research Center, National Institute of Allergy and Infectious Diseases, National Institutes of Health; Bethesda, MD 20892, USA; San Diego Biomedical Research Institute; San Diego, CA 92121, USA; Ragon Institute of MGH, Harvard and MIT; Cambridge, MA 02139, USA

**Keywords:** SCIV, V2-apex, broadly neutralizing antibodies, chimpanzee, immunofocusing, germline-targeting, occluded-open trimer

## Abstract

HIV-1 and its SIV precursors share a broadly neutralizing antibody (bNAb) epitope in variable loop 2 (V2) at the envelope glycoprotein (Env) trimer apex. Here, we tested the immunogenicity of germline-targeting versions of a chimpanzee SIV (SIVcpz) Env in human V2-apex bNAb heavy-chain precursor-expressing knock-in mice and as chimeric simian-chimpanzee immunodeficiency viruses (SCIVs) in rhesus macaques (RMs). Trimer immunization of knock-in mice induced V2-directed NAbs, indicating activation of V2-apex bNAb precursor-expressing mouse B cells. SCIV infection of RMs elicited high-titer viremia, potent autologous tier 2 neutralizing antibodies, and rapid sequence escape in the canonical V2-apex epitope. Six of seven animals also developed low-titer heterologous plasma breadth that mapped to the V2-apex. Antibody cloning from two of these identified multiple expanded lineages with long heavy chain third complementarity determining regions that cross-neutralized as many as 7 of 19 primary HIV-1 strains, but with low potency. Negative stain electron microscopy (NSEM) of members of the two most cross-reactive lineages confirmed V2 targeting but identified an angle of approach distinct from prototypical V2-apex bNAbs, with antibody binding either requiring or inducing an occluded-open trimer. Probing with conformation-sensitive, non-neutralizing antibodies revealed that SCIV-expressed Envs as well as some primary HIV-1 Envs adopted a more open conformation, thereby exposing a conserved V2 epitope that is occluded in closed SIVcpz and HIV-1 Env trimers. These results expand the spectrum of V2-apex targeted antibodies that can contribute to neutralization breadth and identify novel SIV Env platforms for further development as germline-targeting and immunofocusing immunogens.

**One sentence summary:** A cryptic V2 epitope in occluded-open HIV and SIV Env trimers is the target of a new class of V2-directed cross-neutralizing antibodies.

## Introduction

Broadly neutralizing antibodies (bNAbs) represent a key defense against viruses and an important correlate of immune protection of anti-viral vaccines (*1*). However, despite concerted efforts for nearly three decades, there are currently no HIV-1 immunogens that consistently elicit high titers of bNAbs in outbred animals or humans (*2–4*). While humans have the capacity to develop bNAbs, breadth and potency usually appear only after years of HIV-1 infection and only in a subset of individuals (*5–7*). This is because most bNAbs have atypical features such as exceptionally long heavy chain complementarity determining region 3 (HCDR3) segments, high levels of somatic hypermutation, insertions or deletions in variable regions, and low precursor frequencies, all of which pose substantial barriers to traditional vaccine approaches (*4, 8, 9*). Nonetheless, the identification of bNAbs in HIV-1 infected humans, and more recently in simian-human immunodeficiency virus (SHIV) infected rhesus macaques (RMs), has provided proof-of-principle that bNAb responses can be elicited and prompted studies to dissect the pathways of virus-antibody co-evolution as a blue print for vaccine design (*10–16*). It is widely believed that an effective vaccination strategy will need to stimulate rare precursor B cells of multiple bNAb lineages, which will then have to be affinity matured along desired pathways (*4, 17, 18*).

Env trimers display a large antigenic surface that can engage many different B cell receptors. Thus, immunogens that can focus B cell responses to specific epitopes may reduce unwanted off-target responses. We previously reported that certain strains of simian immunodeficiency viruses (SIVcpz and SIVgor) infecting chimpanzees (*Pan troglodytes*) and western gorillas (*Gorilla gorilla*) are exquisitely sensitive to neutralization by human variable loop 2 (V2)-directed bNAbs, but not bNAbs targeting V3 mannose patch, CD4 binding site (CD4bs) and interface epitopes (*19*). This antigenic conservation suggested that SIV-based immunogens might serve to immunofocus B cell responses to V2-apex epitopes shared by HIV-1. Indeed, a minimally-modified near-native soluble SIVcpz Env trimer MT145.Q171K (abbreviated MT145K) was shown to bind inferred germline precursors of several human V2-apex bNAbs and to stimulate one of these in a V2-apex bNAb germline heavy chain (CH01^gH^) expressing knock-in mouse (*20*). Subsequent boosting with a cocktail of V2-apex sensitive HIV-1 Env trimers induced heterologous neutralizing antibodies in a subset of mice (*20*).

There are different strategies to examine the bNAb induction potential of HIV-1 Envs, one of which is in the context of simian-human immunodeficiency virus (SHIV) infection of RMs (*12*). SHIVs express HIV-1 Envs as functional trimers on the surface of infected cells and virions. Moreover, SHIVs replicate continuously over the course of the infection, resulting in an evolving viral quasispecies that can drive antibody somatic hypermutation and maturation (*12, 21*). Indeed, the patterns of envelope-antibody (Env-Ab) co-evolution in SHIV-infected RMs are remarkably similar to those observed in HIV-1 infected humans, indicating similar mechanisms of epitope recognition and neutralizing antibody escape (*12*). Finally, SHIVs expressing V2-apex bNAb sensitive Envs commonly induce V2-directed neutralization breadth in RMs (*12, 21*). Thus, SHIV infection of RMs recapitulates V2-apex and other bNAb development in an animal model that closely approximates HIV-1 infected humans.

Immunofocusing is designed to reduce off-target responses by eliciting B cell recall responses to a shared epitope in otherwise antigenically different immunogens (*22–26*). Here, we examined the immunofocusing potential of Envs from diverse primate lentiviruses and found that several, including Envs from highly divergent SIVs infecting mustached (*Cercopithecus cephus*) and red-tailed (*Cercopithecus ascanius*) monkeys, were exquisitely sensitive to neutralization by mature human V2-apex bNAbs. However, these same Envs were resistant to neutralization by inferred ancestors of V2-apex bNAbs, even after the introduction of mutations previously shown to improve precursor binding (*20*). The exceptions were two V2-engineered SIVcpz Envs, CAM13.Q171K (CAM13K) and DP943.Q171K (DP943K) which, like the previously reported MT145K Env (*20*), were neutralized by the reverted unmutated ancestors (RUA) of PG9, PG16 and CH01. To examine their germline-targeting potential, we selected one of these Envs (CAM13K) together with a minimally modified derivative (CAM13.K169R.K170R.Q171K or CAM13RRK) for immunogenicity studies in human V2-apex bNAb germline-expressing knock-in mice and SCIV-infected RMs. Although the CAM13K SOSIP trimer expanded human V2-apex precursor-expressing mouse B cells and induced V2-directed neutralizing responses, neither SCIV.CAM13K nor SCIV.CAM13RRK induced prototypical V2-apex bNAbs in infected monkeys. Instead, both SCIVs elicited a novel V2-directed antibody specificity that cross-neutralized a number of tier 2 HIV-1 strains with low potency by recognizing a conserved V2 epitope in incompletely closed (occluded-open) Env trimers (*27, 28*).

## Results

### Antigenic conservation of the V2-apex among primate lentiviruses

The apex of the HIV-1 Env trimer in its closed, pre-fusion conformation is comprised of the V1V2 loops of three gp120 protomers, which are positioned around its axis and form the top layer of the glycoprotein (*29–32*). Canonical V2-apex bNAbs access this protein surface through their long anionic HCDR3s that penetrate the glycan shield surrounding N-linked glycans at positions 160 (N160) and 156 (N156) and contact positively charged lysine residues in strand C (position 168-171) of the V2 loop. Testing three human V2-apex bNAbs, we previously found that HIV-1 shares unexpected antigenic cross-reactivity in this epitope with certain SIVcpz and SIVgor strains (*19*). Here, we assembled 33 functional Envs from diverse primate lentiviruses and analyzed their sensitivity to a larger panel of V2-apex bNAbs, including the prototypic PG9, PG16, PGT145, PGDM1400, VRC26.25 and CH01 (*29, 33–36*) as well as to BG1 and VRC38, which have shorter and more compact HCDR3 loops (*37, 38*).

SIV Env expression plasmids were cotransfected in 293T cells with an *env*-minus SIVcpz proviral backbone and the neutralization sensitivity of the resulting pseudoviruses was tested in the TZM-bl assay (*39*). This analysis identified several SIVcpz and SIVgor Env pseudotypes that were uniquely sensitive to human V2-apex bNAbs, with half-maximal inhibitory concentrations (IC_50_) of less than 1 μg/ml (Fig. 1A). The most sensitive Envs (MB897, CAM13, MT145, EK505, DP943), all of which were potently neutralized by PG9, PG16, PGT145, PGDM1400 and VRC26.25, were derived from SIVcpz*Ptt* strains of central chimpanzees (*P. t. troglodytes*), which are the closest relatives of pandemic (group M) HIV-1. Envs from more distantly related gorilla viruses were also sensitive to V2-apex bNAbs, especially to PGDM1400 (IC_50_ <0.7μg/ml), although they were resistant to VRC26.25, CH01, BG1 and VRC38. Envs from the most distantly related SIVcpz*Pts* strains infecting eastern chimpanzees (*P. t. schweinfurthii*) were the least sensitive, although four were neutralized by BG1 (IC_50_ <5 μg/ml).

**Fig. 1.**
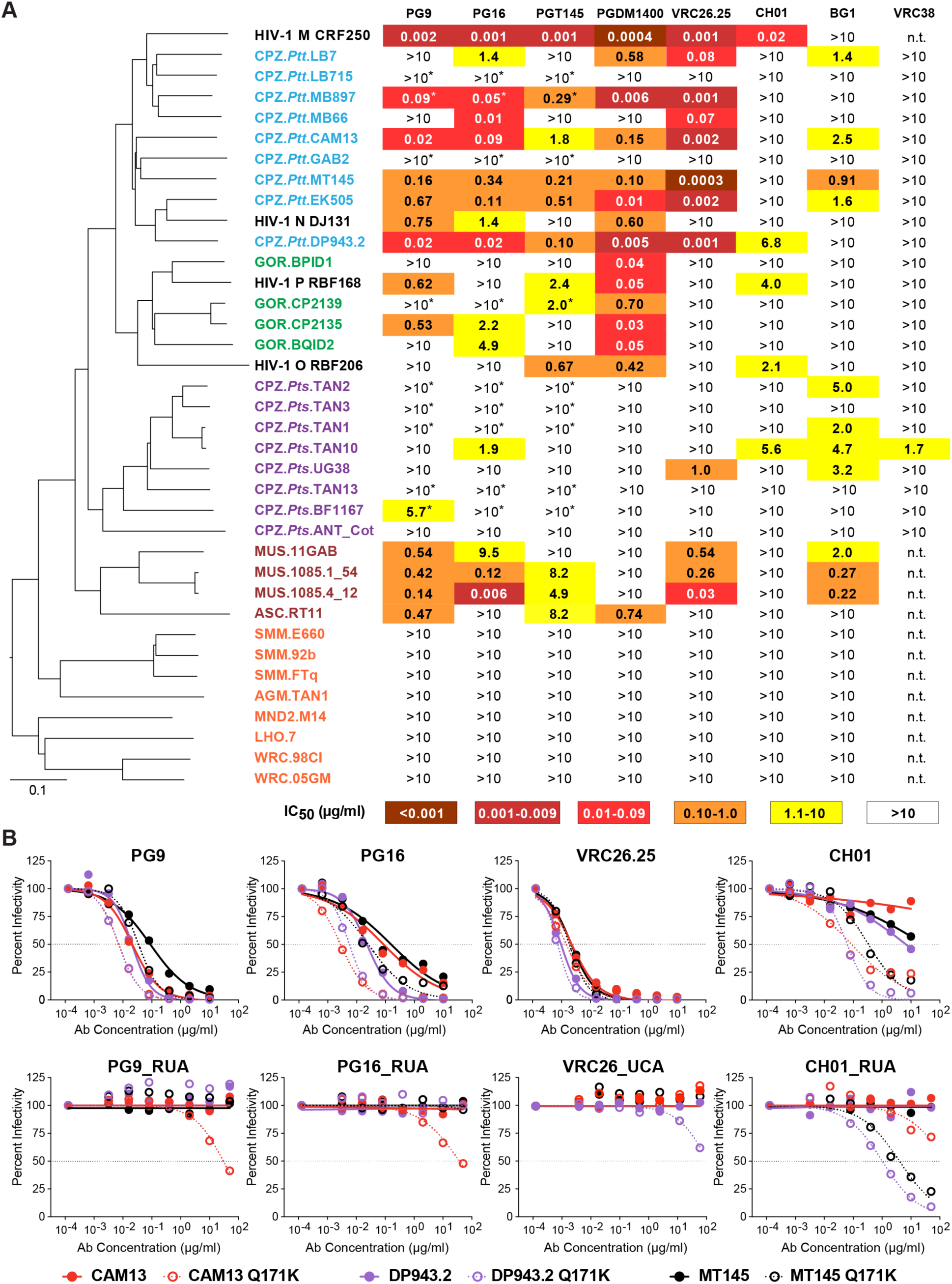
Antigenic conservation of the V2-apex among primate lentiviruses. (A) The ability of human V2-apex bNAbs (top) to neutralize pseudoviruses bearing Envs from diverse SIV strains (left) is shown, with numbers indicating IC_50_ values (μg/ml) as determined in the TZM-bl assay (*39*). Coloring indicates the relative neutralization potency. Values highlighted by asterisks have previously been reported (*19*). SIV Envs are color-coded to indicate their species/subspecies origin (central chimpanzees [CPZ.*Ptt*], blue; western gorillas [GOR], green; eastern chimpanzees [CPZ.*Pts*], purple; moustached [MUS] and red-tailed [ASC] monkeys, brown; sooty mangabeys [SMM], African green monkeys [AGM], mandrills [MND2], L’Hoest’s monkeys [LHO], and western red colobus [WRC], red) and their phylogenetic relationships are illustrated by a neighbor joining tree of full-length Env protein sequences to the left (the scale bar indicates 0.1 amino acid substitutions per site). HIV-1 strains CRF250 (group M), DJ131 (group N), RBF168 (group O) and RBF206 (group P) are shown for comparison (see Table S4 for a list of GenBank accession numbers of all SIV Envs analyzed). The highest antibody concentration used was 10 μg/ml (n.t., not tested). (B) Neutralization curves depicting the sensitivity of pseudoviruses containing wildtype (closed circles) and Q171K mutated (open circles) versions of the SIVcpz Envs CAM13 (red), DP943.2 (blue), and MT145 (black) to mature V2-apex bNAbs PG9, PG16, VRC26.25 and CH01 (top) and their inferred precursors (bottom). Dashed lines indicate 50% reduction in virus infectivity (the antibody concentration is shown on the x-axis in mg/ml).

In addition to SIVcpz and SIVgor strains, we also tested Envs from primate lentiviruses infecting various monkey species (Fig. 1A). These analyses revealed that Envs from some SIVs, such as SIVmus from moustached monkeys and SIVasc from red-tailed monkeys, were also potently neutralized by human V2-apex bNAbs. SIVmus Envs were particularly sensitive to PG9, PG16, VRC26.25 and BG1, while SIVasc was potently neutralized by PG9 and PGDM1400 (IC_50_ <1 μg/ml). In contrast, primate lentiviruses infecting other African monkey species, including SIVsmm from sooty mangabeys (*Cercocebus atys*), SIVtan from tantalus monkeys (*Chlorocebus tantalus*), SIVmnd2 from mandrills (*Mandrillus sphinx*), SIVlho from l’Hoest’s monkeys (*Allochrocebus lhoesti*) and SIVwrc from western red colobus (*Piliocolobus badius*) were resistant to V2-apex bNAbs (Fig. 1A). All of these lacked a potential N-linked glycosylation site (PNGS) at position 156, which is important for most V2 apex bNAbs (*36, 40, 41*), and contained other substitutions in the core V2 epitope consistent with neutralization resistance (Fig. S1). Thus, the antigenic conservation of the trimer apex is not limited to the immediate ape precursors of HIV-1 but is also observed among some SIVs infecting Old World monkeys.

### Germline-targeting potential of V2-apex bNAb sensitive SIV Envs

To examine whether any of the V2-apex bNAb sensitive SIV Envs had germline-targeting potential, we tested their neutralization sensitivity to previously reported reverted unmutated ancestors (RUAs) of PG9, PG16 and CH01 (*42*) as well as the inferred unmutated common ancestor (UCA) of VRC26 (*29*). As expected, none of the wildtype SIV Envs were neutralized by the precursor antibodies (Table S1). Since a glutamine-to-lysine mutation at position 171 (Q171K) of the MT145 Env had conferred the ability to bind to human V2-apex germline precursors (*20*), we tested the effect of this mutation in all other SIV Envs. Moreover, we modified select SIV Envs by introducing additional changes previously reported to improve germline-targeting of HIV-1 Envs (*20, 26*), including removing a PNGS at position 130 (N130K), adding a PNGS at position 156 (K156N + F158S), changing a lysine at position 166 to an arginine (K166R), and/or increasing the number of lysine residues in the C strand (Table S1).

While most of the V2 mutations increased the neutralization sensitivity of the SIV Envs to mature bNAbs (e.g., the combination of Q171K and N130K rendered the MB897 Env markedly more sensitive to PG9, PG16, VRC26.25 and CH01), this was not true for the corresponding precursors (Table S1). Of 20 SIVcpz, SIVgor, SIVmus and SIVasc Env mutants tested, only three were neutralized by the RUAs of PG9, PG16 and/or CH01 at IC_50_ values <50 μg/ml (Fig. 1B; Table S1). These included the previously reported MT145K Env (*20*) as well as two new minimally modified Envs, CAM13K and DP943K, from SIVcpz*Ptt* strains originally isolated from naturally infected chimpanzees in Cameroon (*43, 44*). Both DP943K and MT145K were only neutralized by the CH01_RUA, while CAM13K was neutralized by the RUAs of PG9, PG16 and CH01, although the latter did not reach 50% inhibition (Fig. 1B). Since the CAM13K Env appeared to engage more than one V2-apex bNAb precursor, it was selected for further studies.

### Immunogenicity of a CAM13K SOSIP trimer in CH01 germline heavy chain knock-in mice

To test the germline-targeting potential of the CAM13K Env, we expressed it as a soluble SOSIP trimer (Fig. S2A) using previously reported stabilization mutations (*45–48*). Biolayer interferometry (BLI) showed that the CAM13K trimer bound conformation-dependent V2-apex (PGT145, VRC26.25) as well as V3 (PGT128, PGT125) bNAbs, but not CD4 binding site (VRC01) and interface (PGT151) bNAbs (Fig. S2B). There also was no appreciable binding of non-neutralizing antibodies directed against V2 (CH58) and V3 (19b) as well as CD4-induced coreceptor (17b) and cluster A (A32) epitopes (Fig. S2C). Consistent with the neutralization results (Fig. 1B), the CAM13K SOSIP trimer bound the RUAs of PG9, PG16 and CH01 (Fig. S2B and C). Finally, negative stain electron microscopy (NSEM) showed a well-ordered trimer (Fig. S2D), indicating a native-like Env configuration.

To determine whether the CAM13K trimer could activate human V2-apex bNAb precursor-expressing B cells *in vivo*, we tested it in a previously described knock-in mouse (*20*). In this model, animals are homozygous for the prearranged heavy chain (V_H_DJ_H_^+/+^) of the inferred precursor of the V2-apex bNAb CH01, which then pairs with wildtype mouse light chains (*20*). Although this mouse model has an unphysiologically high precursor frequency, it has been used to test the priming capacity of V2-apex directed immunogens (*20*). Five mice were immunized with the CAM13K SOSIP trimer in glucopyranosyl lipid adjuvant-stable emulsion (GLA-SE), while three mice received only GLA-SE. After six immunizations at 2-week intervals, mice were necropsied and terminal bleeds tested for neutralizing activity (Fig. 2). Although adjuvant administration alone elicited some serum neutralization (reflecting the high precursor frequency), SOSIP immunization induced significantly higher titers to both autologous (CAM13K) and heterologous (C1080, CRF250, Q23) pseudoviruses, which were neutralized in an N160 glycan-dependent manner (Fig. 2). Since the three HIV-1 Envs are sensitive to neutralization by the CH01_RUA (Fig. S3), these findings do not indicate the development of heterologous breadth. However, the data show that a soluble CAM13K Env trimer, despite its extensive antigenic diversity from HIV-1, was able to expand human V2-apex bNAb precursor-encoding mouse B cells and induce a V2-apex directed neutralization response.

**Fig. 2.**
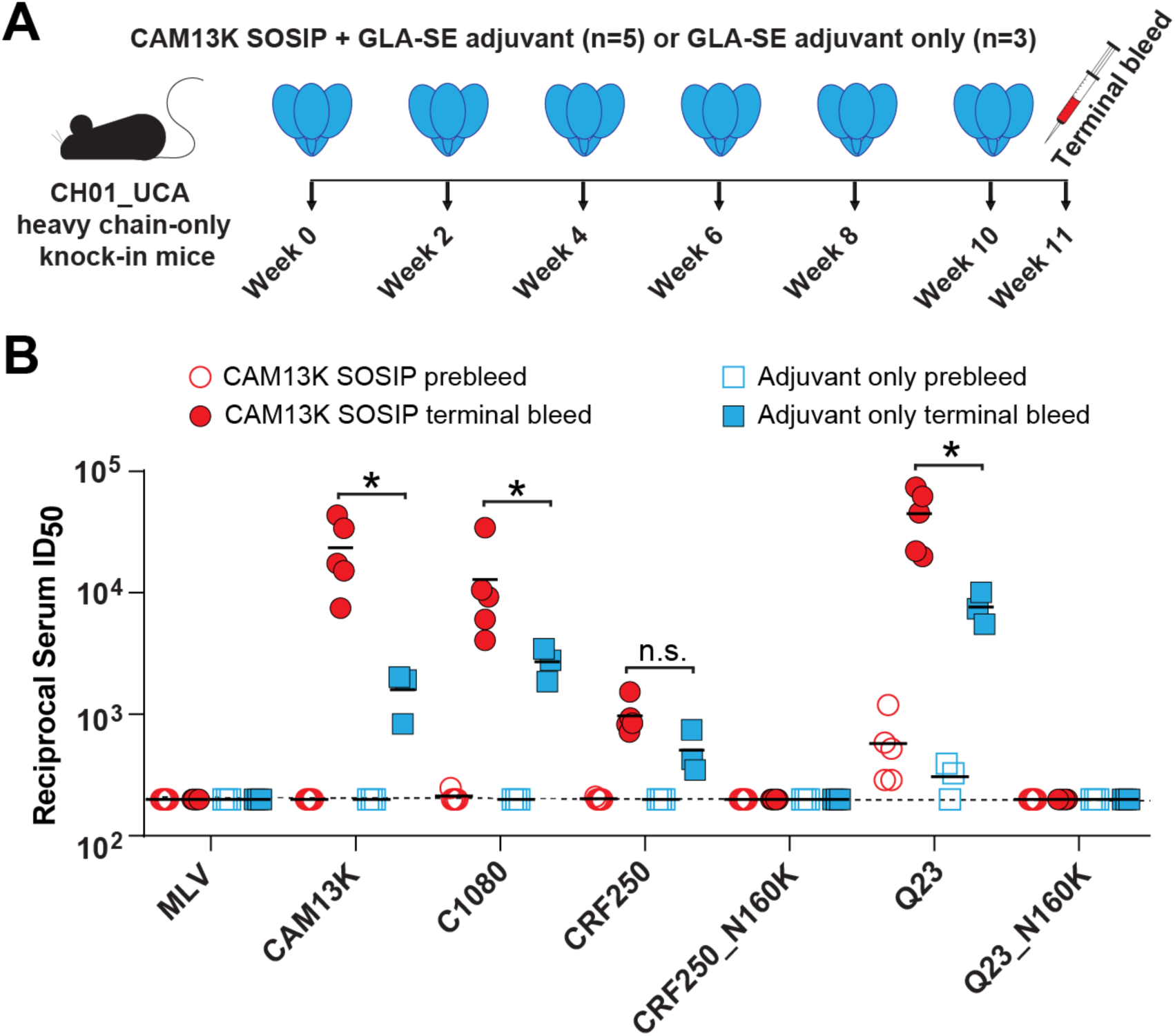
CAM13K SOSIP immunization elicits neutralizing antibody responses in V2-apex bNAb precursor expressing knock-in mice. (A) CH01_RUA heavy chain (HC)-only knock-in mice (*20*) were immunized with either 20 μg CAM13K SOSIP trimer in 5 μg GLA-SE adjuvant (n=5) or with 5 μg GLA-SE adjuvant alone (n=3). Time points of immunization and bleeds are indicated. (B) Comparison of serum neutralization titers (reciprocal 50% inhibitory dilutions, ID_50_) from trimer-immunized (red circles) and adjuvant only (blue squares) knock-in mice (pre-immunization and terminal bleeds are shown as open and filled symbols, respectively). Serum neutralizing activity was tested in the TZM-bl assay (*39*) against pseudoviruses carrying the immunogen-matched autologous CAM13K Env, three CH01_RUA sensitive HIV-1 Envs (C1080, CRF250, Q23) as well as N160 glycan knockout variants of CRF250 and Q23. MLV Env containing pseudovirus was used for control. Differences between immunized and adjuvant-only groups were assessed using a nonparametric Mann-Whitney test (Prism 9.4.0 GraphPad Software), with asterisks indicating p <0.05.

### SCIV induced autologous and heterologous NAb responses in RMs

SHIV infections provide insight into the immunogenicity of HIV-1 Env glycoproteins since these viruses express native trimers that bind and co-evolve with germline and intermediate B cell receptors, thereby driving somatic hypermutation and antibody maturation (*12*). To test the bNAb induction potential of the CAM13K Env in the context of a productive viral infection, we cloned its ectodomain (Fig. 3A) into an SIVmac766 vector previously optimized for SHIV construction (*21*). Since the amino acid residue at position 375 of the HIV-1 Env determines how efficiently the corresponding SHIV replicates in rhesus CD4+ T cells (*21, 49*), we created isogenic mutants of SCIV.CAM13K by changing the wildtype methionine (375M) to serine (375S), tyrosine (375Y), histidine (375H), tryptophan (375W) or phenylalanine (375F) residues. Individual constructs were sequence confirmed, tested for replication competence in rhesus CD4+ T cells *in vitro* (Fig. S4), and used as equal mixtures (based on p27 antigen content) to infect three naïve RMs (T925, T926, T927) by intravenous inoculation (Fig. 3B). Single genome sequencing (SGS) of plasma viral RNA four weeks post-infection identified the 375W variant as the predominant strain in all three animals (Fig. S5). Based on previous signature analyses (*50*), we also replaced two lysine residues at positions 169 and 170 in the C strand of the CAM13K Env with arginine residues, which markedly enhanced neutralization by the reverted unmutated ancestors of PG9, PG16 and CH01 (Fig. S6). By introducing these same mutations into the 375W variant of SCIV.CAM13K, we generated a second germline-targeting SCIV.CAM13RRK strain and infected four additional (repurposed) RMs (see methods for details). All seven animals were treated with anti-CD8 monoclonal antibodies to transiently deplete CD8+ T cells at the time of inoculation and then followed longitudinally for up to 88 weeks to assess viral replication, development of autologous and heterologous neutralizing antibodies, and Env sequence evolution (Figs. 3 and 4).

**Fig. 3.**
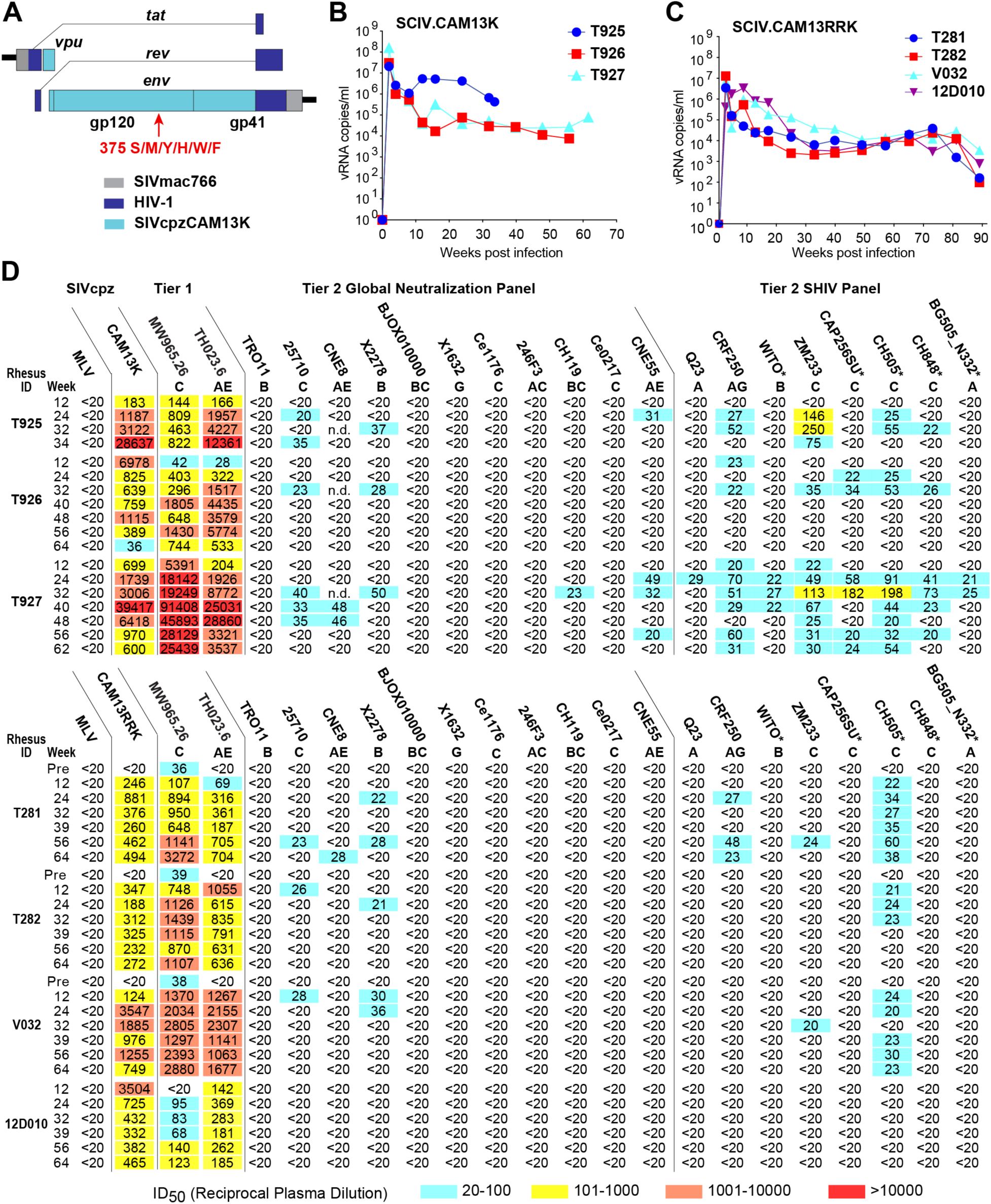
SCIV infected RMs develop low-titer heterologous neutralization breadth. (A) Design scheme of SCIV vectors expressing SIVcpz Env ectodomains. The SIVcpz CAM13K *vpu-env* region (teal) was cloned into an optimized SHIV vector (*21*) consisting of a SIVmac766 proviral backbone (grey) and HIV-1 derived *tat* and *rev* genes (dark blue). Six isogenic SCIV mutants were generated with serine (S), methionine (M), tyrosine (Y), histidine (H), tryptophan (W) or phenylalanine (F) at position 375 of the CAM13K Env. (B) Plasma vRNA kinetics in three RMs infected with SCIV.CAM13K. Animals were inoculated with equal mixtures of SCIV.CAM13K variants bearing all six Env375 mutants. SCIV.CAM13K.M375W emerged as the predominant strain in all animals. (C) Plasma vRNA kinetics in four RMs infected SCIV.CAM13RRK (generated from SCIV.CAM13K.375W by introducing K169R and K170R mutations). (D) Autologous and heterologous neutralization of longitudinal plasma samples from RMs infected with SCIV.CAM13K (top) or SCIV.CAM13RRK (bottom). Titers (reciprocal 50% inhibitory dilutions, ID_50_) are shown for autologous (wildtype CAM13K and CAM13RRK encoding 375M) and heterologous (tier 1 and tier 2) viruses representing different HIV-1 subtypes (A, AG, AE, AC, B, C, BC, G; indicated below virus name), with no reactivity observed against a murine leukemia virus (MLV) Env containing control (all ID_50_ <1:20). Coloring indicates relative neutralization potency. Both Env pseudotypes (CAM13K, CAM13RRK, tier 1 and tier 2 global panel) and replication competent SHIV strains were tested, all of which encoded the wildtype amino acid at position 375, except for SHIV.BG505_N332, which encoded a tyrosine instead of a serine. SHIVs expressing transmitted founder HIV-1 Envs are denoted by asterisks. For three SCIV.CAM13RRK infected animals, which were repurposed from prior HIV-1 immunization studies (see methods), pre-infection (pre) plasma neutralization titers are also shown.

**Fig. 4.**
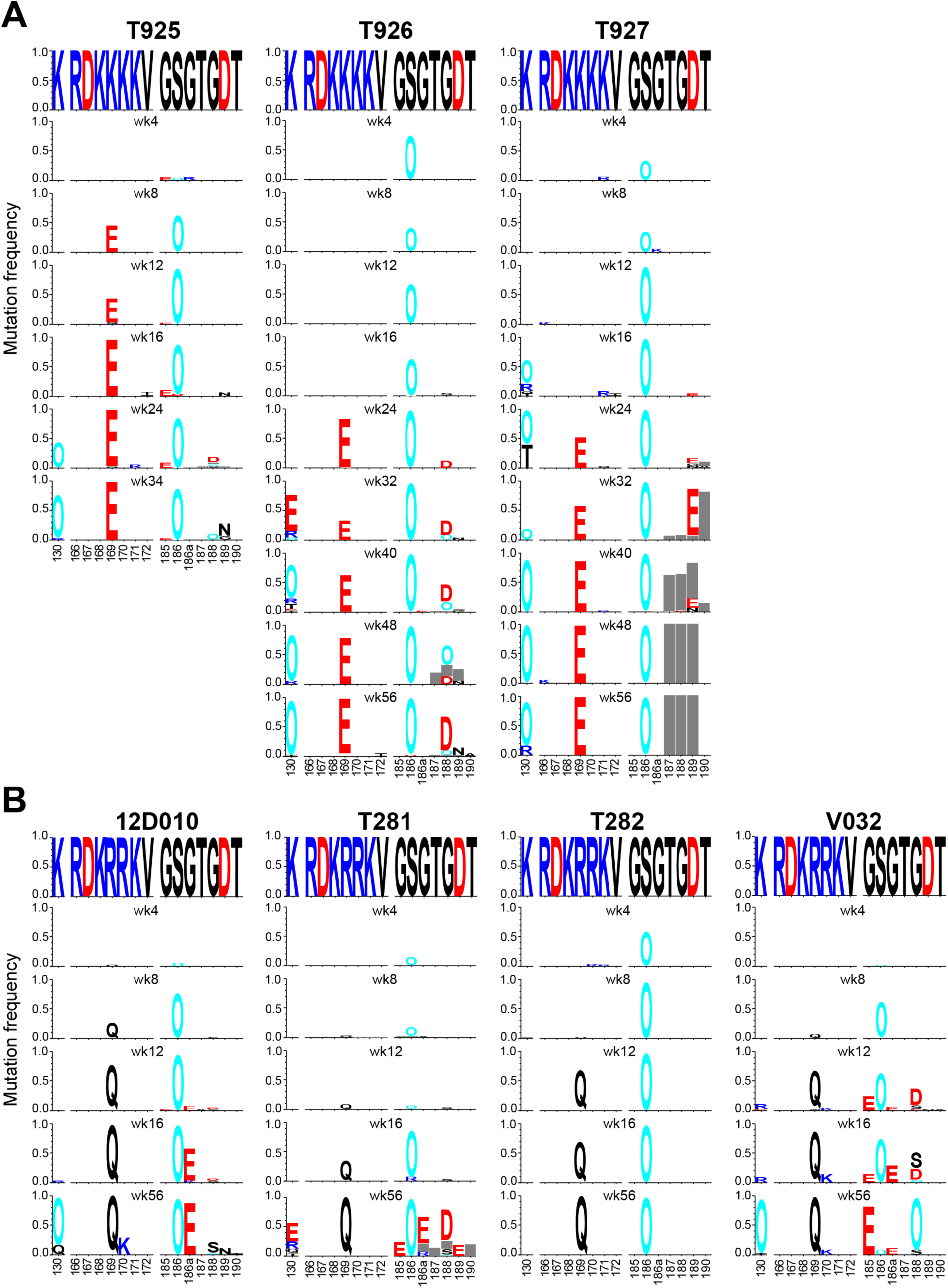
SCIV infection elicits strong V2-apex directed immune selection. (A, B) Longitudinal Env evolution at the V2-apex is shown over time for the N-linked glycosylation site at position 130, strand C (AA 166-172) and the hypervariable part of the V2 loop (AA 185-190) for all SCIV.CAM13K (A) and SCIV.CAM13RRK (B) infected RMs (see Fig. S5 for a similar analysis of the entire Env). Sequences are shown as logo plots, where the height of each amino acid is proportional to its frequency at the respective time point. For each RM, the top logo depicts the infecting SCIV, with subsequent logos representing longitudinal time points denoted in weeks (wk). To highlight mutations, the infecting virus sequence is blanked out for all longitudinal time points. Positively charged amino acids (R, H, K) are colored blue, negatively charged amino acids (D, E) are colored red, potential N-linked glycosylation sites are denoted as “O” and colored cyan, and grey bar indicate deletions. The webtool AnalyzeAlign from the Los Alamos HIV Databases was used for to generate the logos.

Both SCIV.CAM13K and SCIV.CAM13RRK established productive and persistent infections in all animals (Figs. 3B and C). Although there was a trend for higher peak (geometric mean [GM] 4.6 x 10^7^ vs 2.7 x 10^6^ vRNA/ml at week 2) and setpoint (GM 2.3 x 10^5^ vs 1.4 x 10^4^ vRNA/ml at week 24) viral loads in SCIV.CAM13K compared to SCIV.CAM13RRK infected animals, the group sizes were too small to reliably estimate differences. One SCIV.CAM13K infected RM (T925) with a particularly high set point viral load developed an AIDS-like illness and was euthanized at week 32. All SCIV-infected animals developed potent autologous neutralizing antibodies against wildtype (375M) CAM13K and CAM13RRK pseudoviruses, with peak 50% inhibitory dilutions (ID_50_) ranging from 1:350 to 1:39,000 (Fig. 3D). None of the plasma samples neutralized pseudoviruses bearing the murine leukemia virus (MLV) Env used for control. Overall, the SCIV-infected RMs exhibited *in vivo* replication kinetics and autologous neutralizing antibody responses that were similar to those of HIV-1 infected humans and SHIV-infected RMs (*12, 21*).

We also assessed heterologous plasma neutralization using tier 1 (n = 2) and tier 2 (n = 19) HIV-1 strains, which included global panel pseudoviruses (*51*) as well as replication competent SHIV strains (*12*). Plasma samples from all animals potently neutralized the two tier 1 strains, which are known to spontaneously adopt an open Env conformation and are thus susceptible to antibodies that recognize conserved epitopes normally occluded in closed Env trimers (Fig. 3D). Plasma samples from six animals also neutralized up to 11 (of 19) heterologous tier 2 strains, including several transmitted founder Env expressing SHIVs, which are usually neutralization resistant, except for antibodies that target bNAb epitopes (*52*). These heterologous plasma responses were observed very early in infection and mapped to the V2-apex epitope, since mutations at positions 166, 169 or 171 in susceptible Envs reduced their neutralization sensitivity (Fig. S7). Surprisingly, however, all heterologous plasma responses were only weakly cross-neutralizing, with most ID_50_ values just above the 1:20 cut-off. Moreover, there was no appreciable increase in breadth and potency over time (Fig. 3D), as is typically seen in SHIV-infected RMs or HIV-1-infected humans who develop canonical bNAbs. Thus, SCIV infection elicited atypical V2-directed neutralizing antibodies that did not represent V2 bNAbs.

### Env sequence evolution in SCIV infected rhesus macaques

To look for evidence of immune selection in viral variants, we characterized the evolving Env quasispecies in all SCIV infected RMs. This was done by limiting dilution PCR of plasma viral RNA, in which sequencing of single RNA templates allowed the tracking of sequence changes from the transmitted virus over time (Fig. S5). Longitudinal sequences were analyzed using a suite of computational tools designed to investigate different aspects of within-host Env evolution. One of these, the Longitudinal Antigenic Swarm Selection from Intrahost Evolution (LASSIE) program (*53*), identified Env residues where mutations altered the encoded amino acid in at least 80% of sequences at one or more time points. This approach identified several residues at the V2-apex that exhibited signs of immune selection (Fig. 4, Table S2).

The first mutation observed in all animals by week 4 involved the addition of a PNGS at position 186 (S186N) in the V2 loop (Fig. 4). This was followed as early as week 8 by changes at position 169 in the C strand, which is a known contact residue of V2-apex bNAbs (*30, 32, 40, 54, 55*). In all SCIV.CAM13K infected RMs, a lysine at this position was changed to a glutamic acid (K169E) (Fig. 4A), while in all SCIV.CAM13RRK infected RMs an arginine was changed to a glutamine (R169Q), in each case requiring a single nucleotide mutation (Fig. 4B). A third mutation that emerged as early as week 16 was a K130N substitution, which resulted in the addition of an N-linked glycosylation site at position 130. This change was observed in 5 of the 7 SCIV infected animals, and like the changes at position 169, is known to confer resistance to most V2-apex bNAbs (*30, 32, 40, 50, 54, 55*). Although these Env sites were not the only ones displaying signs of selection (Fig. S5, Table S2), they were among the first to change in all, or nearly all, SCIV-infected animals, indicating escape from strong V2-directed antibody pressures.

Since glycan holes can detract from bNAb development (*56*), we also investigated the evolution of the glycan shield in the SCIV-infected RMs. We found that CAM13K and CAM13RRK Envs have five strain-specific glycan holes, three of which were filled during the infection course (Fig. S8). Finally, we examined Env hypervariable loops, which evolve primarily by insertions and deletions, and thus are difficult to align. To preclude alignment artefacts, we used alignment-free characteristics such as length, number of glycans and net charge, which are known to be associated with bNAb sensitivity/resistance (*50*). This analysis showed very similar patterns of variable loop evolution across all RMs (Fig. S9). The most notable changes were deletions in the hypervariable part of V1 (HXB2 positions 132-152), which is exceptionally long in CAM13 (31 AA) and decreased by as many as 14 amino acids. This reduction in V1 loop size was accompanied by a loss of one or two glycosylation sites as well as a decrease in net charge (Fig. S9). There also was a trend of a net charge reduction in the V2 hypervariable loop, but this was not observed in all animals. In contrast, no or only minor changes in length, glycan number and net charge were observed for variable loops 4 and 5, respectively. Since long, negatively charged V1 and V2 loops are associated with resistance to V2-apex bNAbs (*50*), these findings provided further evidence for a strong V2-directed antibody response.

### Cloning of antigen-specific B cells from SCIV-infected rhesus macaques

To characterize the SCIV-induced plasma breadth at the monoclonal antibody (mAb) level, we selected RMs T925 and T927 for single B cell sorting and antibody expression analysis. Both animals were SCIV.CAM13K infected, not previously immunized, and exhibited the highest plasma neutralization titers (Fig. 3D). Matching the sorting probes to the plasma neutralizing activity (Fig. 3D), we used ZM197-ZM233 (*20*) and CAP256 (*55*) fluorophore-labeled SOSIP hooks to isolate antigen-specific B cells. For RM T927, we used both probes to sort peripheral blood mononuclear cells (PBMCs) collected 24 and 32 weeks post-infection but only the ZM197-ZM233 SOSIP to sort lymph node cells obtained at necropsy 62 weeks post-infection. Similarly, for RM T925, we only used the ZM197-ZM233 SOSIP (*20*) to sort B cells from PBMCs obtained 24 weeks post-infection. Starting with ∼10 x 10^6^ PBMC or lymph node cells per sort, we identified multiple expanded lineages in both animals, seven of which were selected for paired heavy and light chain immunoglobulin gene amplifications based on HCDR3 length and D gene usage, which were suggestive of V2-apex bNAbs (Table 1). Select lineage members were then tested for neutralization up to a concentration of 250 μg/ml against autologous (CAM13K) and heterologous (MT145K) SIVcpz Env pseudotypes as well as the same panel of tier 1 and tier 2 HIV-1 and SHIV strains used to characterize the plasma samples (Fig. 5A).

**Fig. 5.**
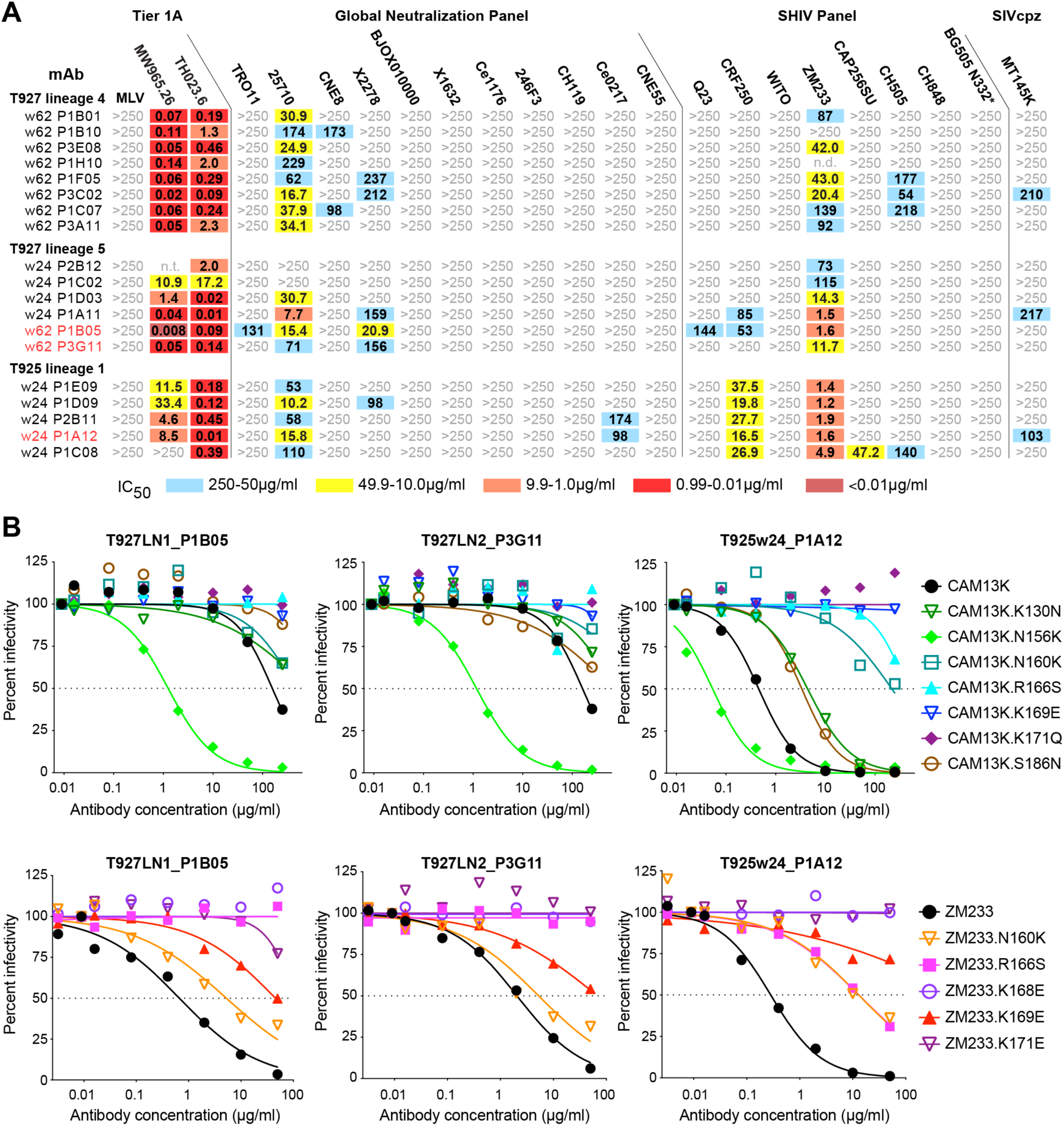
SCIV infections elicit low-potency V2-directed antibodies that cross-neutralize HIV-1. (A) Neutralization breadth and potency of SCIV-induced antibodies cloned from RMs T927 and T925 (indicated on the left). 50% inhibitory concentrations (IC_50_) are shown for representative lineage members (μg/ml) against pseudoviruses and SHIV strains (indicated on top). Only the most potent and cross-reactive lineages are shown (see Fig. S10 for similar results for the remaining lineages). The highest antibody concentration used was 250 μg/ml (coloring indicates relative neutralization potency, n.t., not tested). mAbs analyzed by negative stain electron microscopy are highlighted in red (heavy and light chain variable region sequences are shown in Fig. S12). (B) Epitope mapping of select cross-neutralizing mAbs. Neutralization curves are shown for three monoclonal antibodies (indicated on top and highlighted in red in panel A) against a panel of CAM13K and ZM233 mutants (indicated on the right). Dashed lines indicate 50% reduction in virus infectivity (corresponding IC_50_ values are shown in Fig. S11).

**Table 1.**
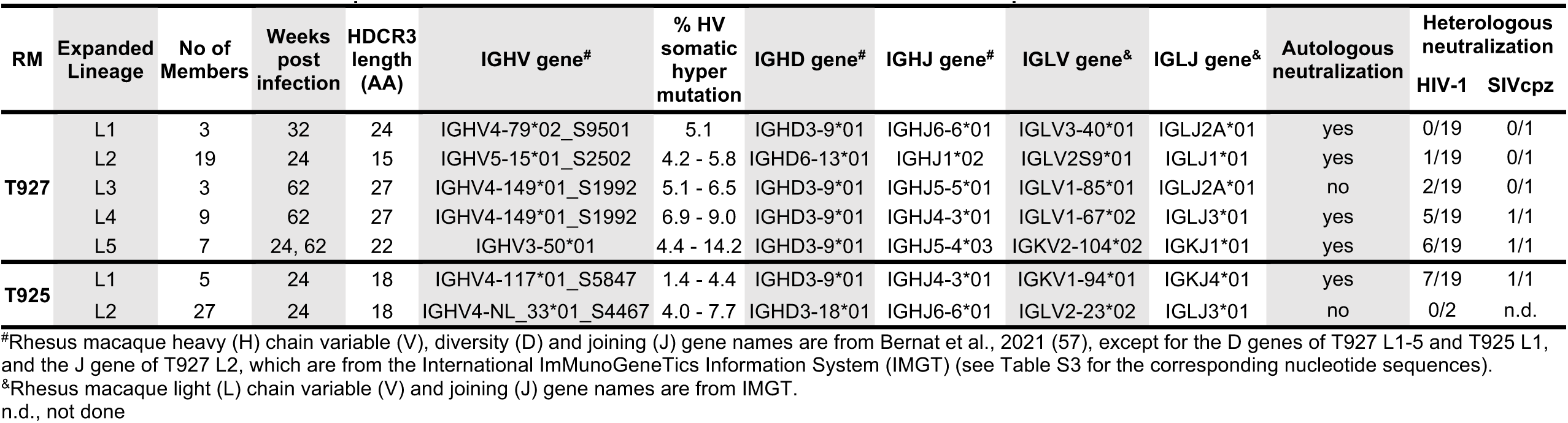
Immunogenetics of Env-specific monoclonal antibodies from SCIV infected rhesus macaques.

Of the five expanded lineages identified in RM T927, three exhibited no or very limited heterologous activity, with lineage 1 neutralizing no tier 2 HIV-1, lineage 2 neutralizing only ZM233, and lineage 3 neutralizing ZM233 and 25710, but not the autologous CAM13K strain (Figs. S10 and S11). The remaining two lineages, however, exhibited greater heterologous breadth, with lineage 4 neutralizing as many as 5 of 19 (26%) and lineage 5 as many as 6 of 19 (32%) tier 2 HIV-1 strains (Table 1, Fig. 5A). The viruses neutralized by both lineages were 25710, ZM233 and X2278, although most mAbs required very high antibody concentrations to achieve 50% inhibition. One member of each lineage also weakly neutralized the SIVcpz strain MT145K (IC_50_ >200 μg/ml). Antibody isolation from the second animal T925 yielded very similar results. Of the two expanded lineages, only lineage 1 exhibited heterologous breadth, with all members neutralizing 25710, CRF250 and ZM233, and a subset also neutralizing X2278, CE0217, CAP256SU, CH505 and MT145K strains, again with low potency (Fig. 5A). Surprisingly, almost all lineage members neutralized one or both tier 1 viruses with high potency, suggesting that they targeted conserved epitopes in incompletely closed HIV-1 Env trimers (Fig. 5A). Overall, the isolated mAbs accounted for most of the heterologous (tier 2) plasma activity. However, their potency was universally low despite the presence of several features characteristic for V2-apex bNAbs, such as long HCDR3s (18 to 27 AA), up to 14% heavy chain variable (V) gene somatic hypermutation, and the usage of the heavy chain diversity (D) gene HD3-9*01 (also known as HD3-15*01; Table S3) (*57, 58*), which encodes an EDDY motif predicted to be tyrosine sulfated (Table 1; Figure S12).

The epitopes targeted by the isolated mAbs were mapped using autologous (CAM13K) and heterologous (ZM233 and MW965.26) Env mutants. Given the patterns of early sequence evolution (Fig. 4) and the plasma mapping results (Fig. S7), we focused on amino acid substitutions in the central V2 loop. Testing only members of the most potent lineage in each animal, we found that substitutions at positions 166 (R166S), 168 (K168E), 169 (K169E), and/or 171 (K171E, K171Q) either reduced or abrogated neutralization of autologous and heterologous viruses (Figs. 5B, Fig. S11). Similarly, removal of the PNGS at position 160 (N160K) increased resistance of most lineage members, although this mutation did not completely abrogate neutralization. Addition of glycosylation sites at positions 130 (K130N) and 186 (S186N) in CAM13K, which represented early escape mutations in SCIV-infected RMs (Fig. 4), also reduced neutralization, but had overall only a modest effect (Figs. 5B, Fig. S11). Surprisingly, removal of the PNGS at position 156 (N156K), which is required by most V2-apex bNAbs (*36, 40, 41*), rendered CAM13K up to two orders of magnitude more sensitive to neutralization (both ZM233 and MW965.26 lack the N156 glycan). This was observed for all lineages, even the much less potent and cross-reactive ones that failed to neutralize the autologous CAM13K strain (Fig. S11). These results indicate that SCIV-induced cross-neutralizing antibodies target the V2 loop, but in a manner distinct from prototypic V2-apex bNAbs.

### Epitope mapping by negative stain electron microscopy

To characterize their modes of V2 interaction, we generated Fab fragments for members of the most cross-reactive lineage from RMs T927 (P3G11 and P1B05) and T925 (P1A12), respectively, and analyzed their binding to soluble Env trimers by negative stain electron microscopy (NSEM) using both CAM13K and ZM197-ZM233 SOSIP preparations. In all attempts to visualize Fab-Env complexes, we failed to observe binding of any of the three Fabs to SOSIP trimers, with most two-dimensional (2D) class averages showing trimers with no Fabs bound (Figs. S13A-15A). However, a smaller fraction of particles appeared to consist of single Fabs bound to single Env gp120/gp41 protomers (Fig. 6A), which allowed us to generate three-dimensional (3D) reconstructions (Fig. 6B). Although such structures have not previously been reported, Fab densities were readily resolved and a single gp120/gp41 protomer fit well into the remaining density. This analysis allowed us to infer the binding site and approximate angle of approach for each Fab, which in all three cases identified a region in the V2 loop near the N156 and N160 glycans (spanning Env positions 158-172) as the likely epitope (Figs. S13B-15B). However, unlike prototypical V2-apex bNAbs, all three Fabs used an angle of approach that when modeled onto a closed, prefusion trimer structure resulted in clashes with adjacent protomers (Figs. S13C-15C). This was true for P1B05 and P3G11, which used a more horizontal angle of approach compared to canonical bNAbs (Figs. S13C and S14C). This was also true for P1A12, which bound closer to the trimer axis very similar to PG9 (Fig. 6C), but appeared to be rotated, thereby resulting in clashes with neighboring protomers (Fig. S15C). Importantly, such clashes were not observed when Fab binding was modeled onto an occluded-open trimer (Figs. S13D-15D), in which the protomers are rotated away from the central axis, but the V1-V3 loops are still in their closed position shielding the V3 co-receptor binding site (*27, 28*). These data suggested that the SCIV-induced cross-neutralizing antibodies either required or induced an occluded-open Env trimer.

**Fig. 6.**
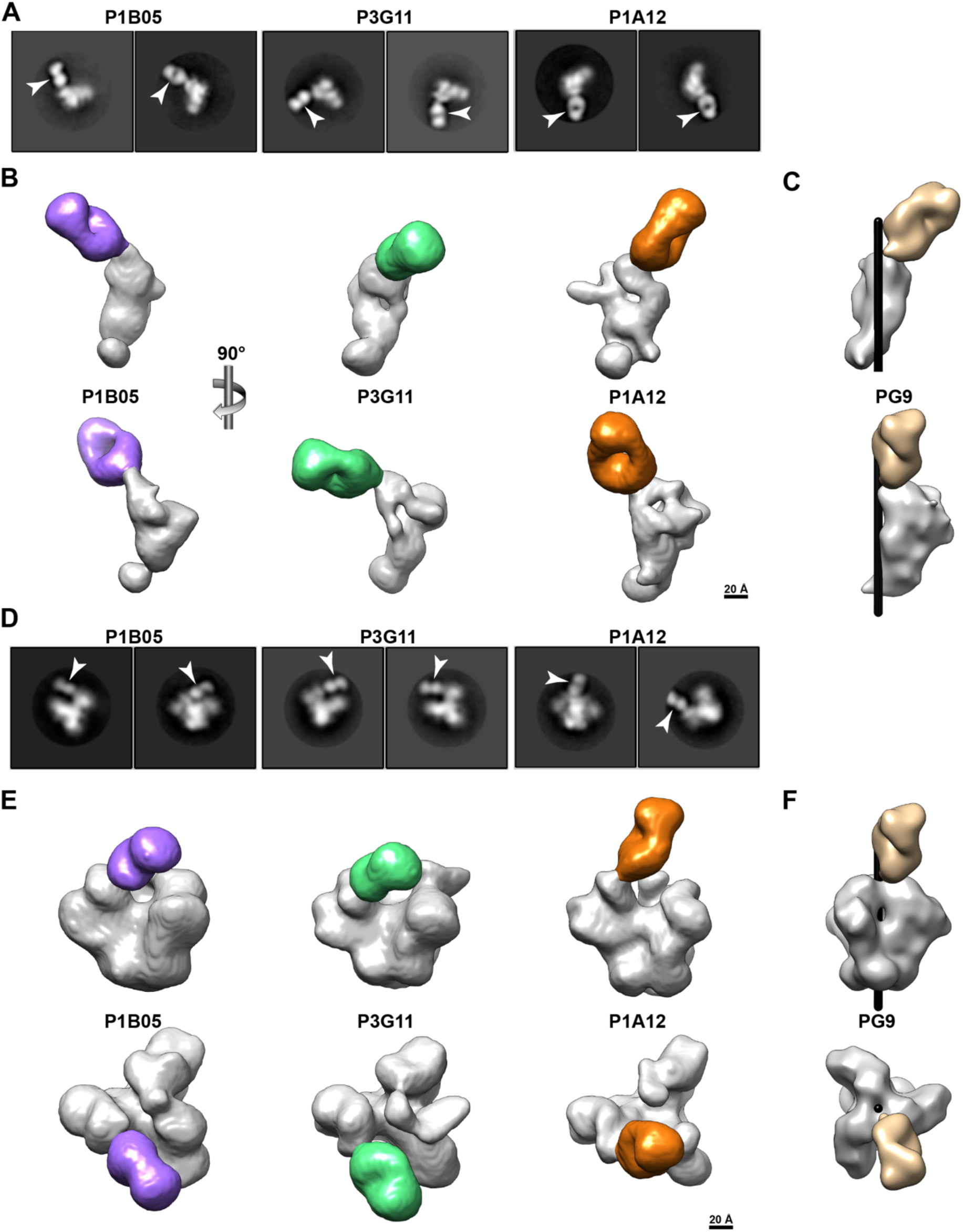
SCIV-induced V2-directed cross-neutralizing antibodies bind occluded-open but not closed trimers. (A) 2D class averages of NSEM images showing Fabs (arrows) bound to single Env protomers. Fabs (denoted on top) were generated for members of the most cross-reactive antibody lineage from RM T927 (P3G11 and P1B05) and RM T925 (P1A12), respectively (also see Fig. 5). Although both ZM197-ZM233 and CAM13K SOSIP preparations were used, only Fab-monomer complexes were observed. (B) 3D reconstructions of Fab-monomer complexes, shown in two orthogonal views, with the Env domain in gray and the Fab in color (P1B05, purple; P3G11, green; P1A12, gold). The scale bar indicates 20 Å. (C) Volume rendering of the PG9 Fab (tan, PDB 3U2S) in complex with a prefusion BG505 trimer (PDB 5FYL), of which only a single protomer (grey) is shown. The Env densities in (B) were aligned with the Env density in (C) and shown in the same orientation. The black cylinder denotes the position of the 3-fold trimer axis. (D) 2D class averages showing Fabs (arrows) bound to SOSIP trimers lacking the N156 glycan (CH505_N156Q SOSIP). (E) 3D reconstructions of Fabs bound to CH505_N156Q SOSIP trimers. Both side (upper) and top (lower) views are shown, with the Fabs highlighted in color as in (B). The densities of the CH505_N156Q SOSIP are consistent with an occluded-open trimer. The scale bar indicates 20 Å. (F) Volume rendering of PG9 (tan, PDB 3U2S) in complex with a closed, prefusion BG505 trimer (PDB 5FYL), shown in the same orientation as in (E). The black cylinder indicates the central 3-fold axis of the Env trimer.

The NSEM modeling data suggested that the SCIV-induced antibodies bound Env trimers that were neither completely open (*59*) nor completely closed (*60*). To demonstrate this directly, we performed NSEM using an N156 glycan-deficient SOSIP trimer. We reasoned that such a trimer was more likely to be partially open since the absence of the N156 glycan is known to reduce inter-V2-strand interactions and apex stability (*41*). Moreover, all SCIV-induced cross-reactive NAbs neutralized N156-deficient CAM13K with markedly increased potency (Fig. S11). We thus examined the binding of the three Fabs to a SOSIP trimer in which the asparagine at position 156 was changed to a glutamine (CH505.N156Q_SOSIP). In contrast to all previous attempts, this SOSIP yielded Fab-trimer complexes that were readily visualized in 2D class averages (Fig. 6D). Corresponding 3D reconstructions revealed Fab binding to the V2 region, with the overall trimer density consistent with an occluded-open (Fig. 6E) and not a closed (Fig. 6F) conformation. Importantly, the Fab-monomer structures fit well into the corresponding Fab-trimer structures (Figs. S13E-15E), thus validating their utility to infer putative epitopes and approximate angles of approach. Indeed, the Fab-trimer structures suggested the same epitopes as identified by the Fab-monomers, i.e., in the V2 loop near residues 158-172 and glycans N156 and N160 (Figs. S13F-15F). Thus, weakly cross-neutralizing antibodies from two different SCIV-infected rhesus macaques bound to a cryptic V2 epitope that appeared to only be accessible in an occluded-open trimer conformation.

### Probing SCIV-expressed and wildtype Envs with conformation-sensitive antibodies

To examine whether the CAM13K and CAM13RRK Envs inherently adopt a more open conformation, we tested their sensitivity to conformation-sensitive V2i (697-D, 1393A) (*61*), V2p (CH58, CAP228-3D) (*42, 62, 63*), linear V3 (3074, 447-52D) (*64, 65*) and CD4i (17b, A32) (*66, 67*) antibodies as well as the V2-apex bNAb PG9 for control. These antibodies were selected because they recognize epitopes that are occluded in prefusion trimers and thus only neutralize viruses whose Envs are not completely closed. Using antibodies up to a concentration of 100 μg/ml, we found that pseudoviruses carrying full-length CAM13K and CAM13RRK Envs with a wildtype methionine at position 375 (PV_CAM13K_375M, PV_CAM13RRK_375M) were completely resistant to the above conformation-sensitive V2, V3 and CD4i antibodies (Fig. 7A). This was also observed for a SCIV construct that encoded a methionine at position 375 (SCIV.CAM13K_375M), thus excluding the possibility that the chimeric SCIV gp41 domain (Fig. 3A) influenced the stability of the timer apex. In contrast, SCIV constructs with a tryptophan at position 375 (SCIV.CAM13K_375W, SCIV.CAM13RRK_375W), which represented the preferred *in vivo* variant, as well as a CAM13RRK pseudovirus in which the 375M was mutated to a 375W (PV_CAM13RRK_375W) were sensitive to monoclonal antibodies CAP228-3D, 3074, 447-52D and 17b, which neutralized all three strains with remarkably consistent potencies (Fig. 7A). These results indicate that the amino acid substitution at position 375 rendered CAM13K and CAM13RRK Env conformationally more flexible, with the bulky 375W shifting the equilibrium toward a more open state. Interestingly, the SIVcpz MT145K Env, which naturally encodes a histidine at position 375, was resistant to neutralization by CAP228-3D, 3074, 447-52D and 17b, both as a pseudovirus and as a SCIV construct (Fig. 7A). Thus, the propensity to adopt a more open conformation is not an inherent characteristic of SIVcpz Envs or SCIV constructs, but is an Env-specific property.

**Fig. 7.**
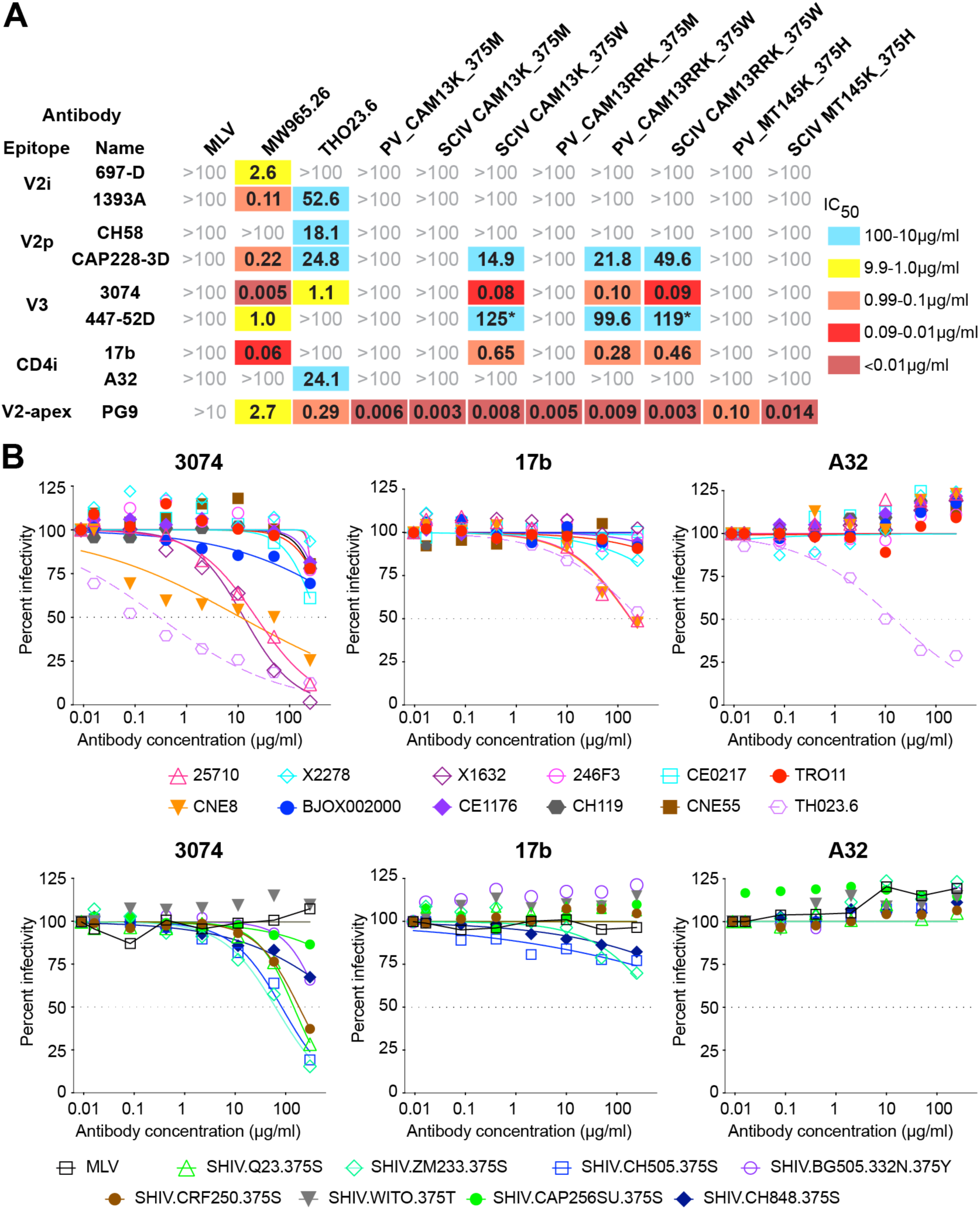
SCIV expressed, but not wildtype SIVcpz Env, as well as some primary HIV-1 Envs adopt a more open conformation. (A) The sensitivity of CAM13K and CAM13RRK Envs to non-neutralizing V2p, V2i, linear V3 and CD4i antibodies as well as the V2-apex bNAb PG9 (indicated on the left) is shown for pseudovirus (PV) and SCIV constructs encoding either the wildtype methionine (M) or the RM-selected tryptophan (W) at position 375 (indicated on top). 50% inhibitory concentrations (IC_50_) are shown in μg/ml (coloring indicates relative neutralization potency, with asterisks identifying IC_50_ values estimated by the Prism software). Also shown are IC_50_ values for pseudovirus and SCIV expressed MT145K Env, which encodes a histidine at position 375. MLV and tier 1 HIV-1 (MW965.26, TH023.6) Env pseudoviruses are shown for control. (B) Neutralization curves are shown for non-neutralizing V3 (3074) and CD4i (17b, A32) antibodies (indicated on top) against global panel pseudoviruses (upper panel) and SHIV strains (lower panel; with the amino acid residue at position 375 indicated). Dashed lines indicate 50% reduction in virus infectivity.

Finally, we examined the conformational state of global panel pseudovirus Envs and SHIV strains that were used to characterize the SCIV-induced cross-neutralizing antibodies (Fig. 7B). As reported previously (*52*), we found that several global panel pseudoviruses, including 25710 that was neutralized by all SCIV-induced cross-neutralizing antibodies, were sensitive to the linear V3 antibody 3074. This was also true for SHIV.Q23, SHIV.ZM233, SHIV.CRF250 and SHIV.CH505, which despite encoding a wildtype amino acid residue at position 375 were sensitive to 3074 neutralization, albeit only at high antibody concentrations (IC_50_ >50 μg/ml). Although none of the tier 2 viruses were neutralized by A32, global panel strains 25710 and CNE8 were sensitive to 17b, as was SHIV.ZM233, although the latter virus did not reach an IC_50_ (Fig. 7B). These data confirm that some primary HIV-1 Envs are prone to sample a more open conformation and to expose an otherwise cryptic V2 epitope in the context of an occluded-open trimer. Although the V1V2 region is highly variable, several sites in the C strand are relatively conserved (Fig. S1), thus explaining the sensitivity of these HIV-1 strains to neutralization by SIVcpz Env-induced antibodies.

## Discussion

The HIV-1 trimer apex elicits both non-neutralizing and neutralizing antibodies, including some of the most potent bNAbs identified to date (*68*). We previously reported that some SIVcpz strains share unexpected antigenic cross-reactivity with HIV-1 at this epitope (*19*) and that a C-strand modified SIVcpz Env bound inferred human V2-apex bNAb precursors with high affinity (*20*). These observations suggested that SIV Envs, which are otherwise antigenically highly divergent from HIV-1, could be used to focus B cell responses to this bNAb epitope. Here, we used our extensive collection of primate lentiviral Envs to systematically analyze the antigenic conservation of the V2-apex across divergent SIV lineages. We found that SIVs infecting African apes and some Old World monkeys are exquisitely sensitive to neutralization by mature human V2-apex bNAbs, indicating that the quaternary structure of the trimer apex is highly conserved. This sensitivity was further increased by modifications in the C strand, which also rendered two new SIVcpz Envs susceptible to neutralization by inferred human V2-apex bNAb precursors. Thus, SIV Envs, and especially SIVcpz Envs, may have utility as germline-targeting and/or immunofocusing components of an AIDS vaccine. While priming with an SIV trimer would generate B cell responses to the entire Env surface, a subsequent boost with an HIV-1 trimer that shared only the V2-apex bNAb site should focus the B cell recall response to this epitope. In addition, the backbone diversity of the SIV Envs may reduce germinal center competition for B cells with long HCDR3s that target the V2-apex. There is increasing evidence that sequential immunization with divergent HIV-1 Env immunogens favors B cell recall responses to shared epitopes (*22-24, 26, 69*), and a derivative of the SIVcpz MT145K trimer is being produced for safety and immunogenicity studies in humans (*4*). Thus, the additional SIV Envs characterized here provide a unique collection to develop a wider array of priming and boosting immunogens.

To test the utility of an SIVcpz Env priming immunogen, we generated a soluble CAM13K SOSIP trimer and evaluated its germline-targeting properties in knock-in mice that expressed the reverted unmutated ancestor of the human V2-apex bNAb CH01 (*20*). We found that CAM13K SOSIP immunization of these mice elicited antibodies that neutralized CH01_RUA sensitive HIV-1 strains in an N160 dependent manner, demonstrating the successful stimulation and expansion of precursor antibody expressing mouse B cells. However, this knock-in mouse model expresses only a single prearranged inferred germline heavy chain at an unphysiologically high precursor frequency. Moreover, the RUA of CH01, like that of PG9 or PG16, was inferred from very few lineage members and thus represents only an approximation of the unmutated common ancestor (*35, 68*). Given these limitations, it seems clear that the germline-targeting potential of the CAM13K Env and future derivatives will have to be confirmed in additional, more relevant mouse models, such as long human CDR3 rearranging mice, as well as ultimately in outbred animals and humans (*70–72*).

SHIVs evolve continuously in infected RMs, thus allowing for the selection of Env variants with enhanced affinity through sequential rounds of antibody binding and virus escape, leading in some animals to a broadening of the neutralizing response (*12*). Indeed, we recently reported that SHIVs expressing transmitted founder Envs, which in humans elicited bNAbs, did the same in RMs, with the molecular patterns of Env-Ab co-evolution mirroring those observed in HIV-1 infected humans (*12*). To determine whether a similar approach could be used to examine the bNAb induction potential of SIVcpz Envs, we cloned CAM13K and CAM13RRK Envs into an optimized SHIV vector and used the resulting SCIV constructs to infect RMs. We found that both SCIVs closely resembled SHIVs in their *in vivo* replication kinetics and other biological properties. Both SCIVs grew to high titers in human and macaque CD4+ T cells *in vitro* and caused persistent infections *in vivo*, resulting in high peak (10^6^-10^8^ copies/ml) and setpoint (10^3^-10^5^ copies/ml) viral loads. Both SCIVs preferred a bulky aromatic amino acid at position 375, which has been shown to confer enhanced affinity to the rhesus CD4 receptor and thus an improved ability to replicate *in vivo* (*21, 73*). Finally, both SCIVs induced potent autologous tier 2 neutralizing antibodies, consistent with a high antigenic burden and the expression of intact Env trimers. Recent data have shown that SHIV infections are not only useful for tracing bNAb development, but also represent an important outbred animal model for HIV-1 transmission, prevention, immunopathogenesis and cure studies (*74–81*). Our SCIV data show that this model is not restricted to HIV-1 Envs but can be extended to more divergent Envs from primate lentiviruses.

Given that a subset of SHIV infected RMs develop V2-apex bNAbs that closely resemble those of humans (*12*), we reasoned that SCIV infection may induce similar responses. Indeed, six of seven RMs developed limited plasma breadth that mapped to the C strand and was accompanied by rapid viral escape at positions 169 and 186 in the V2 loop, suggesting nascent bNAb development (*12*). However, neither SCIV.CAM13K nor SCIV.CAM13RRK infected RMs ultimately developed appreciable V2-apex bNAbs, although B cell cloning identified antibodies that cross-neutralized a number of tier 2 HIV-1 strains with low potency. Negative stain electron microscopy, coupled with conformational probing of SCIV and wildtype SIVcpz Envs, provided an explanation: the introduction of a tryptophan at position 375, which mediated efficient *in vivo* replication, rendered SCIV-expressed CAM13K and CAM13RRK Envs conformationally more open, as exemplified by an increased sensitivity to V2p, linear V3 and CD4i antibodies. This propensity may have impeded the development of bNAbs, even if appropriate V2-apex bNAb precursors were, in fact, stimulated as suggested by the expansion of long HCDR3 antibody lineages, including many that encoded the HD3-9*01 gene thus far identified in all RM-derived V2-apex bNAbs.

The finding of a cryptic V2 epitope that is exposed on occluded-open, but not closed, Env trimers is consistent with recent structural analyses of antibodies detected in sequentially SOSIP-immunized RMs (*28*). Examining immunization-elicited neutralizing antibodies that target the CD4 binding site, Yang and colleagues found that Fab binding was only observed when Envs were not constrained in their closed prefusion conformation (*28*). Instead, these CD4bs NAbs bound an occluded-open trimer, which also exposed V1V2 regions that were buried in closed trimers. Although no antibody specific for these regions was isolated, the cryo-EM data suggested the presence of a cryptic epitope (*28*). Our results confirm and extend these findings, showing that cryptic V2 epitopes are not only present on SOSIP trimers, but also on membrane-anchored Envs on the surface of infected cells or virions, which has implications for immunogen design. However, higher resolution structures of Fab/trimer complexes will be required to fine map the respective epitopes and identify contact residues.

In addition to prototypic V2-apex bNAbs, two classes of V2-directed non-neutralizing antibodies have been described. One of these, termed V2p, includes both human and macaque antibodies that recognize the V2 loop in a helix-coil conformation and require a glutamic acid-aspartic acid (ED) LCDR2 motif for V2 peptide binding (*42, 62, 63*). The other class, designated V2i, recognizes conformational epitopes on gp120 (*82*) and has recently been shown to belong to the large family of CD4i antibodies (*61*). Our SCIV-derived antibodies lack the ED LCDR2 motif and bind the V2 domain in an occluded-open trimer. Moreover, they neutralize primary viruses, whereas V2p and V2i antibodies only neutralize tier 1 isolates (*42, 62, 83*). Thus, the SCIV-derived monoclonal antibodies comprise a new, fourth class of V2-directed antibodies that is weakly cross-neutralizing.

Although the breadth and potency of the SCIV-induced NAbs are limited, these findings do not indicate that SIVcpz Envs are unsuitable immunogens. CAM13K and CAM13RRK Envs with a methionine at position 375 were resistant to V2, linear V3 and CD4i antibodies, indicating a closed conformation, and this was also true for the SIVcpz MT145K Env, both in the context of pseudovirus and SCIV constructs. Moreover, SHIV-expressed Envs are not inherently more open than the corresponding wildtype Envs from which they are derived. A recent study showed that amino acid substitutions at position 375 did not alter the antigenicity of 10 transmitted founder Env expressing SHIVs (*21*). Nonetheless, it is clear that the HIV-1 envelope glycoprotein is conformationally flexible (*84, 85*), and that some HIV-1 strains sample open Env conformations more readily than others (*52, 86*). Since the latter are likely to elicit responses similar to SCIV.CAM13K, it will be necessary to use conformation sensitive probes, such as the antibodies described here, to examine immunogens and SHIV/SCIV constructs prior to vaccination and infection studies. In particular, mRNA-expressed membrane-bound Envs should be tested since more open conformations exposing cryptic epitopes are not limited to soluble SOSIP trimers. On the other hand, greater conformational flexibility may also have benefits, e.g., by enhancing Env binding to bNAb precursors. It will thus be important to explore both the advantages and disadvantages of Env immunogens that reversibly expose some surfaces that are buried in closed Env trimers.

In summary, we examined the antibody response to germline-targeting versions of the SIVcpz CAM13 Env, both as a soluble SOSIP immunogen and as infectious SCIVs in RMs. Although SCIV infection failed to elicit canonical V2-apex bNAbs, we found that C-strand modified versions of the CAM13 Env bound several human V2-apex bNAb precursors and stimulated one of these in knock-in mice. We also characterized a new class of V2-directed antibodies that targets a conserved epitope in occluded-open SIVcpz and HIV-1 Env trimers. Although the breadth and potency of this new specificity are limited, our findings expand the spectrum of V2-apex targeted antibodies that can contribute to neutralization breadth and identify a novel SIVcpz Env platform for further development as germline-targeting and immunofocusing immunogens.

## Materials and Methods

### Study design

In this study we examined the bNAb induction potential of two germline-targeting versions of a chimpanzee SIV Env (CAM13K and CAM13RRK) by (i) immunizing human V2-apex bNAb precursor-expressing knock-in mice with a soluble CAM13K SOSIP trimer and by (ii) infecting seven rhesus macaques with chimeric simian-chimpanzee immunodeficiency viruses expressing CAM13K and CAM13RRK Env ectodomains. The number of immunized and infected animals were dependent on availability and not predetermined by power calculations. The number of samples collected and the tests performed are detailed in Figs. 2 and 3.

### Knock-in mouse immunogenicity study

CH01_RUA homozygous “HC only” (V_H_DJ_H_^+/+^) knock-in (KI) mice were generated on the C57BL/6 background as described (*20*). All vaccinations were performed using 6-12-week-old mice. Five mice received 20 μg of the CAM13K SOSIP trimer plus 5 μg of glucopyranosyl lipid adjuvant-stable emulsion (GLA-SE), while three mice received 5 μg of GLA-SE alone. All animals were immunized six times at 2-week intervals by intramuscular injection at two sites (25 μl each) in the hind limbs (Fig. 2A). Blood samples were collected at pre-bleed (pre) and one week after the last immunization and sera were tested for neutralizing activity using a panel of autologous and CH01_RUA sensitive heterologous pseudoviruses. Serum samples were heat inactivated for potential complement activity at 56°C for 30 min. All mice were housed at Duke University and cared for in an AAALAC accredited facility. All animal procedures were approved by the Duke Institutional Animal Care and Use Committee (IACUC).

### SCIV infection of nonhuman primates

Indian rhesus macaques were housed at Bioqual, Inc., Rockville, MD, according to guidelines of the Association for Assessment and Accreditation of Laboratory Animal Care (AAALAC) standards. Experiments were approved by the University of Pennsylvania and Bioqual Institutional Animal Care and Use Committees (IACUC). All RMs were socially housed with a variety of recommended environmental enrichments. They were observed daily and any signs of disease or discomfort was reported to the veterinary staff. RMs were sedated for blood draws, anti-CD8 mAb infusions and SCIV inoculations. All animals received an intravenous infusion of 25 mg/kg of the anti-CD8b mAb CD8beta255R1 at the time of SCIV inoculation to transiently reduce CD8+ cells, thereby allowing for higher peak and set point viral loads. RMs T925, T926 and T927 received 1 ml of a SCIV.CAM13K infection stock (a total of 300 ng of p27 antigen) containing six different Env alleles encoding either the wildtype methionine (M), or serine (S), tyrosine (Y), histidine (H), tryptophan (W) or phenylalanine (F) at position 375 (50ng of p27 antigen each) by intravenous injection. RMs T281, T282, V032 and 12D010 received 1 ml of SCIV.CAM13RRK, which encoded a W at position 375 (50 ng of p27 antigen). Animals underwent sequential blood draws (Fig. 3) to obtain plasma and PBMCs. Cells were counted, tested for viability by trypan blue exclusion, and aliquoted at a concentration of 5 to 10 x 10^6^ cells/ml in CryoStor CS5 cryopreservation media (Sigma-Aldrich). Mononuclear cells collected from lymph nodes (LN) harvested at necropsy were processed in a similar manner (*12, 21*). Plasma viral loads were determined as previously described (*12, 21*). Three SCIV.CAM13RRK infected animals were repurposed from prior HIV-1 immunization studies. T281 and T282 received HIV-1 subtype C peptides containing liposomes, while V032 received CH505 SOSIP immunizations as described (*87*). Although pre-infection plasmas of these animals weakly neutralized the tier 1 strain MW965.26, none had detectable tier 2 neutralizing antibodies prior to SCIV inoculation (Fig. 3D). Animal 12D010 developed progressive pancytopenia.

### SIV Env expression plasmids and pseudovirus generation

The nucleotide sequences of the SIV Envs used for V2-apex neutralization studies have previously been reported (see Table S4A for GenBank accession numbers and references), except for the CPZ.Pts.TAN10 Env, which was amplified from the spleen of a naturally SIVcpz infected chimpanzee (Ch-036) who died of an AIDS-like illness in Gombe National Park (*88*) and the CPZ.Pts.ANT_Cot Env, which was derived from a limiting dilution plasma viral isolate obtained from an SIVcpzANT infected captive chimpanzee (*89*). Briefly, SIV *env* genes were synthesized (some as wildtype and others as codon-optimized sequences, see Table S4A) and cloned into the pcDNA3.1 expression vector (Sigma-Aldrich). SIV Env pseudotypes were produced in 293T cells by co-transfection of the SIV *env* expression vectors with an Env-deficient SIVcpzMB897-EnvFS backbone, titered on TZM-bl cells as described (*19*), and tested for sensitivity to the mature V2-apex bNAbs PG9, PG16, PGT145, PGDM1400, VRC26.25, CH01, BG1 and VRC38 (*29, 33–36*) as well as the PG9_RUA, PG16_RUA, CH-01_RUA (*42*) and the VRC26_UCA (*29*) in the TZM-bl assay (GenBank accession numbers of heavy and light chain variable domain sequences are listed in Table S4B). A subset of SIV Envs was further modified by site-directed mutagenesis (Q5 Site-Directed Mutagenesis Kit; New England BioLabs) to incorporate amino acid changes previously reported to improve the germline-targeting capacity of HIV-1 Envs (*20, 26*). Mutations included the removal of a PNGS at position 130 (N130K), the addition of a PNGS at position 156 (K156N + F158S), the change of a lysine at position 166 to an arginine (K166R), and/or the addition of positively charged C strand residues (K166R, T169K, T170K, E170K, S170K, A170K, Q171K, H171K), alone and/or in combination (GenBank accession numbers of mutant Envs are listed in Table S4C).

### Neutralizing antibody assay

The neutralization capacity of rhesus macaque plasma and monoclonal antibodies was assessed using the TZM-bl assay as described (*39*). Briefly, 96-well plates were seeded with TZM-bl cells (15,000 cells per well) overnight in Dulbecco’s modified Eagle’s medium (DMEM) containing 10% fetal bovine serum (FBS) and 100 U/ml Penicillin-Streptomycin-Glutamine (Gibco). Serial 5-fold dilutions of RM plasma (1:20, 1;100, 1:2,500, 1:12,500, 1:62,500, 1:312,500) or monoclonal antibodies (e.g., 250, 50, 10, 2, 0.04, 0.08, 0.016, 0.00 μg/ml) were incubated with transfection-derived virus at a multiplicity of infection (MOI) of 0.3 in a total volume of 100 μl in the presence of DEAE-dextran (40 μg/ml) for 1 h at 37°C, and this mixture was then added to TZM-bl cells. After 48 h, TZM-bl cells were analyzed for luciferase expression using a Synergy Neo2 Multimode Microplate reader (Bio-Tek) with Gen5 version 1.11 software. Uninfected cells were used to correct for background luciferase activity. The infectivity of each virus without plasma or antibodies was set at 100%. The 50% inhibitory concentration (IC_50_) is the antibody concentration that reduces by 50% the relative light units (RLUs) compared with the no Ab control wells after correction for background. Nonlinear regression curves were determined and IC_50_ values calculated by using variable slope (four parameters) function in Prism software (v8.0). All monoclonal antibodies were tested in duplicate on at least two independent occasions. Since amounts of mouse and rhesus macaque plasma were limited, samples were generally tested only once in duplicate.

CAM13K, CAM13RRK and global panel pseudovirus (*51*) as well as SHIV/SCIV (*12, 21*) viral stocks were generated by transfection of 293T cells. Briefly, 100-mm tissue culture dishes were seeded with 4×10^6^ 293T cells overnight in DMEM containing 10% FBS and 100 U/ml Penicillin-Streptomycin-Glutamine. Cells were transfected by adding 0.5 ml of a preincubated DMEM solution containing 4.5 μg of SIVcpz (SIVcpzMB897-EnvFS; see Table S4A for GenBank accession number) or HIV-1 (SG3Δenv) backbone plasmids and 30 ng of codon optimized or 1.5 μg of wildtype SIVcpz and HIV-1 Env plasmids, respectively, or 6 μg of SCIV or SHIV construct DNA, and 18 μl of FuGENE 6 transfection reagent (Promega), according to manufacturer’s recommendations. The cells were incubated at 37°C in a CO2 incubator for 48-72 h, and supernatant was harvested and stored at −80 °C in 0.5-ml aliquots.

### SCIV construction and infectious stock preparation

SCIV.CAM13K was generated using a previously reported (second generation) SHIV vector (pCRXTOPO.SHIV.V2.backbone1) (*21*), which consists of an SIVmac766 proviral backbone (a transmitted founder clone derived from the SIVmac251 isolate) that contains partial *tat/rev* and gp41 sequences from the transmitted founder HIV-1 strain D.191859 (Fig. 3A). This vector allowed the unidirectional cloning of a CAM13K *vpu-env* fragment (*env* nucleotides 1 to 2153, HXB2 numbering) into unique BsmBI restriction enzyme sites (since BsmBI cleaves outside its recognition site, the enzyme recognition sequence is not conserved after insert ligation in the final construct). The CAM13K *vpu-env* fragment was amplified from the SIVcpz CAM13 proviral clone (*43*), followed by mutation of a glutamine at position 171 to a lysine (Q171K) to improve V2-apex bNAb germline-targeting (*20*). The SCIV construct was then used to create allelic variants at position 375 using the QuikChange II XL site-directed mutagenesis kit (Agilent). Wild-type and mutant plasmids were transformed into MAX Efficiency Stbl2 competent cells (ThermoFisher Scientific) for maxi-DNA preparations and each SCIV genome was sequenced to confirm its integrity. The SCIV.CAM13RRK construct was generated by replacing two wildtype lysine residues at positions 169 and 170 in the Env of SCIV.CAM13K_375W with arginine residues (see Table S4A for GenBank accession numbers of SCIV.CAM13K and SCIV.CAM13RRK). Infectious stocks were generated by transfecting 293T cells (*21*).

### SCIV replication in RM CD4+ T cells

RM CD4^+^ T cells were isolated from PBMCs using the non-human primate CD4+ T cells isolation kit (Miltenyi Biotec), activated using the NHP T-Cell Activation/Expansion kit (Miltenyi Biotec) and used to determine the replication kinetics of each SCIV.CAM13K Env375 variant (*12, 21*). For each variant, transfection derived supernatant containing 300 ng p27Ag was added to 2 x 10^6^ activated RM CD4 T cells in the presence of DEAE-dextran (20 μg/ml) using an estimated MOI of 0.01 to 0.05. Cell and virus mixtures were incubated for 2 hours at 37°C to facilitate infection, washed three times, and resuspended in RPMI1640 medium containing 15% FBS, 1% Penicillin-Streptomycin and 30U/ml IL2. Cells were plated into 24-well plates at 2 x 10^6^ cells in 1 ml and cultured for 13 days, with sampling of 0.2 ml supernatant and media replacement every 2-3 days. Supernatants were assayed for p27 antigen production by ELISA (Zeptometrix).

### Single genome amplification

Single genome amplification of 3’ half genomes or viral *env* genes from plasma RNA and proviral DNA was performed as previously described (*90, 91*). Briefly, ∼20,000 copies of viral RNA were extracted from plasma using QIAamp Viral RNA kit (Qiagen) and reverse transcribed using SuperScript III Reverse Transcriptase (Invitrogen). Viral cDNA was then endpoint diluted and amplified using nested PCR with primers and conditions as previously reported (*90, 91*). Geneious software was used for alignments and sequence analysis (see Table S4D for GenBank accession numbers of SCIV.CAM13K and SCIV.CAM13RRK longitudinal *env* gene sequences).

### Longitudinal Env evolution in SCIV infected RMs

Longitudinal Env evolution analyses were performed as described (*12*). Briefly, Longitudinal Antigenic Swarm Selection from Intrahost Evolution (LASSIE) (https://github.com/phraber/lassie) was used to identify amino acid/glycan mutations under selection in each RM, using 80% or higher loss of the transmitted virus sequence as the cutoff. LASSIE-selected sites shared across RMs were identified (Table S2). The webtool AnalyzeAlign was used to calculate sequence logos (https://www.hiv.lanl.gov/content/sequence/ANALYZEALIGN/analyze_align.html) that show evolution at such sites. Glycan shield mapping was performed using the Glycan Shield Mapping tool (https://www.hiv.lanl.gov/content/sequence/GLYSHIELDMAP/glyshieldmap.html) (*56*). This method maps PNGS to a reference trimer structure and assumes that a 10Å cutoff around each PNGS is glycan shielded. Rare glycan holes are identified by comparison with M-group Envs that are >50% or >80% glycan shielded. For longitudinal glycan shield mapping, consensus glycans (>50%) and median hypervariable loop lengths were used from each host and each time point. For hypervariable loop evolution, we first investigated if there were any repeated indels across RMs. Because of the stochastic nature of hypervariable loop indels and the inherent difficulty of aligning these loops, such an analysis is only feasible by careful inspection of repeated patterns of indels in early longitudinal Envs. Since this analysis did not identify repeated indels in different RM hosts, hypervariable loops from longitudinal Envs were characterized using alignment-free characteristics (https://www.hiv.lanl.gov/content/sequence/VAR_REG_CHAR/index.html), such as length, number of glycans and net charge. Hypervariable loop positions were identified based on HXB2 numbering: 132-152 for hypervariable V1, 185-190 for hypervariable V2, 396-410 for hypervariable V4, and 460-465 for hypervariable V5.

### Production of biotinylated trimer probes for B cell sorting

AVI-tagged ZM197-ZM233V1V2 (*20*) and CAP256-wk34c80-RnS-3mut-2G (*55*) SOSIP trimers were produced using transient transfection in FreeStyle 293F cells (ThermoFisher Scientific) by pre-mixing C-terminal AVI-tagged ZM197-ZM233V1V2 or C-terminal AVI-tagged CAP256-wk34c80-RnS-3mut-2G with a furin expression plasmid using Polyethylenimine (PEI) 40K (Polysciences, Inc.) or Turbo293 (Speed BioSystems) transfection reagents, respectively. The cells were cultured for 4-6 days before AVI-tagged trimers were purified from supernatants using either PGT145 or 2G12 affinity chromatography. The Protein A resin captured AVI-tagged ZM197-ZM233V1V2 was biotinylated with BirA enzyme (Avidity) and cleaved from the Fc purification tag with HRV3C protease. After overnight incubation, the cleaved and biotinylated ZM197-ZM233V1V2 was concentrated using Amicon Ultra-15-30K (Millipore), and purified using a Superdex 200 16/600 gel filtration column (Cytiva) equilibrated with PBS. The 2G12 affinity column captured AVI-tagged CAP256-wk34c80-RnS-3mut-2G protein was eluted with 3M magnesium chloride solution, concentrated using Centricon Plus-70-30K (Millipore), followed by a Superdex 200 16/600 gel filtration column. The corresponding trimer fractions were pooled, negatively selected using a V3 cocktail column containing six V3-directed antibodies (1006-15D, 2219, 2557, 2558, 3074, and 50.1), and biotinylated with BirA enzyme followed by final purification over a Superdex 200 16/600 gel filtration column.

### Rhesus single cell B cell sorting

Single HIV-1 Env–specific B cells were isolated from the blood (weeks 24 and 32) and lymph node (week 62) of RM T927 and from the blood (week 24) of RM T925, both of which were infected with SCIV.CAM13K. Cryopreserved cells collected at these timepoints were thawed and stained with LIVE/DEAD Fixable Aqua Dead Cell Stain (Life Technologies). Cells were washed and stained with an antibody cocktail against CD3 (clone SP34-2, BD Biosciences), CD4 (clone OKT4, BioLegend), CD8 (clone RPA-T8, BioLegend), CD14 (clone M5E2, BioLegend), CD20 (clone 2H7, BioLegend), IgG (clone G18-145, BD Biosciences), IgD (polyclonal, Dako), and IgM (clone G20-127, BD Biosciences) at room temperature in the dark for 30 min. Cells were then washed and incubated with avi-tagged and biotinylated ZM197-ZM233V1V2 (*20*) and CAP256-wk34c80-RnS-3mut-2G (*55*) SOSIP trimers conjugated to phycoerythrin (PE) or allophycocyanin (APC) for 30 min at room temperature. The stained cells were then washed three times with PBS, resuspended in 1 ml of PBS containing 5% FBS and 0.1% sodium azide, and passed through a 70 μm cell mesh (BD Biosciences). Since double positive cells were rare, memory B cells that bound to one or the other of these probes (CD3-CD4-CD8-CD14-CD20+IgD-IgM-IgG+probe+) were isolated with a FACSAria cell sorter using the FACSDiva software (BD Biosciences) and flow cytometric data were subsequently analyzed using FlowJo (v10.8.1). B cells were sorted at 1 cell per well in a 96-well plate containing 20 μl lysis buffer as described (*12*). Plates were frozen on dry ice and stored at −80°C.

### Rhesus macaque monoclonal antibody production

Heavy and light chain variable region genes were amplified using single cell PCR approaches as described (*92*). Briefly, immunoglobulin gene transcripts from single B cells were reverse transcribed with Superscript III (Invitrogen) using random hexamer primers (Thermo Scientific). The complementary DNA was then used as a template for two rounds of nested PCR for heavy and light chain amplification, with amplicons examined by agarose gel electrophoresis. Amplicons were Sanger sequenced (Genewiz), and gene sequences were computationally analyzed using SONAR (*93*), IgBLAST (*94*), and IMGT V-QUEST (*95*). SONAR provided initial gene assignment, sequence annotation, and clonotype clustering based on CDR3 similarity, while IgBLAST and V-QUEST provided additional annotations. For V-QUEST, the IMGT rhesus macaque reference database was used, while for SONAR and IgBLAST an alternate rhesus gene reference database with additional diversity was used (*58*). Final lineage assignments were made after review of all automatic annotations and grouping results by looking for evidence of common descent, such as similar junctions and shared somatic hypermutation. As a final step of germline gene assignment, all heavy chain sequences were compared to a recent macaque reference (*57*), and the closest-matching gene was retained as the final germline gene.

To produce select antibodies, paired heavy (VDJ) and light (VJ) chain variable gene sequences were commercially synthesized (GenScript) and cloned into antibody expression plasmids (see Table S4E for GenBank accession numbers of heavy and light chain variable regions). Recombinant antibodies were produced by cotransfecting paired heavy and light chain expression plasmids into Expi293F cells using ExpiFectamine 293 transfection reagents (ThermoFisher Scientific), purified from culture supernatants using the Protein A/Protein G GraviTrap kit (GE Healthcare), and buffer-exchanged into PBS as described (*12*).

Fab fragments were generated by inserting a stop codon six amino acids upstream of the hinge region (CPPCP) of the heavy chain expression plasmid using the Q5 Site-Directed Mutagenesis Kit (New England BioLabs). This mutagenized plasmid was co-transfected with the respective light chain plasmid, harvested by centrifugation to remove debris, and the supernatant was filtered through 0.22 µm filter and then passed through a CaptureSelect™ CH1-XL Affinity Matrix column (ThermoFisher Scientific). Recombinant Fab protein was eluted with 50 mM acetate buffer at pH 5.0 and concentrated using Ultra-15 Centrifugal Filters (Millipore). Protein concentration was determined using the Qubit protein assay (Invitrogen).

### Biolayer interferometry

The antigenicity of the CAM13K SOSIP trimer was examined by biolayer interferometry (BLI) using an OctetRed96 platform (Sartorius) and a panel of monoclonal antibodies (mAbs), including (i) bNAbs that bind to the CD4-binding site (VRC01), glycan-V3 (PGT128, PGT125, 2G12), V2-apex (PGT145, VRC26.25), and the gp120-gp41 interface (PGT151), (ii) conformation sensitive non-bNAbs that bind linear V3 (19b), V2 (CH58), and CD4i (17b, A32) epitopes, as well as (iii) reverted unmutated ancestors of bNAbs (PG16_RUA, PG9_RUA, CH01_RUA). Antibodies were diluted to 20μg/mL and captured onto anti-hIgG Fc capture (AHC) biosensors for a period of 300s and then washed with PBS pH7.4 running buffer. The antibody captured biosensors were then submerged into wells containing CAM13K SOSIP protein diluted to 50μg/ml for 400s followed by a dissociation period of 600s in 1x PBS. Non-specific binding was assessed using a control anti-influenza haemagglutinin mAb (CH65) for reference subtraction and the binding curves were analyzed using the ForteBio Data Analysis Software 10.0 (Sartorius). Binding responses (nm) of the SOSIP to each antibody were measured at 390-395s of the association phase after reference subtraction and the binding response was normalized (binding ratio = binding in nm of mAb/binding in nm of PGT151) relative to the reference mAb PGT145.

### CAM13K and CH505.N156Q SOSIP trimer expression

To generate a soluble CAM13K Env trimer immunogen, we introduced several stabilizing mutations, including the original SOSIP.664 mutations (A501C-Y605C; I559P; truncation at residue 664) (*45, 46*), the DS stabilizing mutations (I201C-G433C) (*47*), and a hexa-arginine furin cleavage site (6R) (*48*) (Fig. S2A). The gene was codon optimized using GenArt software (ThermoFisher), synthesized and cloned into VRC8400 at SalI and BamHI sites (GenScript). Similarly, to generate a N156Q mutant CH505 SOSIP trimer, we mutagenized the CH505 transmitted founder Env and stabilized it with a chimeric gp41 as well as E64K and A316W substitutions to reduce V3 loop exposure (*96*). SOSIP expression plasmids were transfected into Freestyle 293F cells (ThermFisher) using 293Fectin together with a furin-encoding plasmid to improve gp120/gp41 cleavage. Six days post transfection the supernatant was cleared of cells, concentrated, filtered, and subjected to PGT145 affinity chromatography by gravity flow columns or on a AKTA Pure (Cytvia) as previously described (*96*). Protein eluted from the PGT145 affinity column was 0.2 μm-filtered and concentrated prior to performing size exclusion chromatography. SOSIP trimers were purified in 10 mM Tris pH8, 500 mM NaCl on a HiLoad Superose6 16/600 column (Cytvia). All proteins were snap-frozen and stored at −80°C. SOSIP trimer quality was assessed by antibody binding and NSEM.

### Negative stain electron microscopy and single particle reconstructions

Fab-SOSIP complexes were formed by mixing 10 µg of SOSIP with 36 µg of Fab, and bringing the final volume to 100 µl with buffer containing 20 mM HEPES, 150 mM Na_2_SO_4_, 5 mM NaN_3_, pH 7.4, and incubating overnight at 22 °C. The next day, samples were diluted with 900 µl of buffer, then 1 ml of 0.16% glutaraldehyde in the same buffer was added and incubated for 5 minutes. 160 µl of 1 M Tris buffer, pH 7.4, was added to quench unreacted glutaraldehyde and samples were transferred to a 2 ml, 100 kDa Amicon centrifugal concentrator and spin-concentrated to ∼50 µl. Concentrated samples were then diluted to a nominal SOSIP concentration of 0.2 mg/ml with buffer containing 20 mM HEPES, 150 mM NaCl, and 5 g/dl glycerol, applied to a glow-discharged carbon-coated EM grid for 8-10 second, blotted and stained with 2 g/dL uranyl formate for 1 min, and then blotted and air-dried. Grids were examined on a Philips EM420 electron microscope operating at 120 kV and nominal magnification of 49,000x, and ∼100 images for each sample were collected on a 76 Mpix CCD camera at 2.4 Å/pixel. Images were analyzed by 2D class averages and 3D reconstructions calculated using standard protocols with Relion 3.0 (*97*).

### Statistical analyses

Differences between immunized and adjuvant-only knock-in mouse groups were assessed using a nonparametric Mann-Whitney test using GraphPad Prism 9.4.0 software. We chose a nonparametric rank-based test because the distributions of antibody titers can be highly variable and we had small sample sizes. Statistical significance of hypervariable loop characteristics was determined using a non-parametric two-sided Kendall Tau test.

## Supplementary Materials

Fig. S1. Sequence conservation in the V2-apex bNAb core epitope among primate lentiviruses.

Fig. S2. Generation and characterization of a CAM13K SOSIP trimer.

Fig. S3. Sensitivity of HIV-1 Env pseudoviruses to neutralization by the inferred human V2-apex precursor CH01_RUA.

Fig. S4. Replication kinetics of SCIV.CAM13K allelic variants *in vitro*.

Fig. S5. Env sequence evolution in SCIV-infected rhesus macaques.

Fig. S6. Sensitivity of CAM13RRK to neutralization by V2-apex bNAb precursors.

Fig. S7. Low-titer heterologous plasma breadth in SCIV-infected RMs maps to the V2-apex.

Fig. S8. Glycan shield evolution in SCIV.CAM13K and SCIV.CAM13RRK infected RMs.

Fig. S9. Env hypervariable loop evolution in SCIV.CAM13K and SCIV.CAM13RRK infected RMs.

Fig. S10. Expanded antibody lineages isolated from SCIV.CAM13K infected RMs with no or very limited tier 2 neutralization breadth.

Fig. S11. SCIV-induced cross-neutralizing antibodies target the V2 region.

Fig. S12. Heavy and light chain variable region sequences of SCIV-induced cross-neutralizing antibodies.

Fig. S13. The SCIV-induced cross-neutralizing antibody P1B05 from RM T927 binds a V2 epitope that is occluded in a closed, prefusion trimer.

Fig. S14. The SCIV-induced cross-neutralizing antibody P3G11 from RM T927 binds a V2 epitope that is occluded in a closed, prefusion trimer.

Fig. S15. The SCIV-induced cross-neutralizing antibody P1A11 from RM T925 binds a V2 epitope that is occluded in a closed, prefusion trimer

Table S1. Neutralization sensitivity of wildtype and modified SIV Envs to V2-apex bNAbs and their inferred precursors.

Table S2. Env sites under selection in SCIV.CAM13K and SCIV.CAM13RRK infected rhesus macaques.

Table S3. Varying nomenclature in different databases for the same rhesus macaque germline heavy (H) chain variable (V), diversity (D) and joining (J) gene sequences.

Table S4. GenBank Accession Numbers.

## Acknowledgments

We thank F.-H. Lee and N. Chohan for expert technical assistance; M. Gondim for generating a plasma virus isolate from an SIVcpzANT infected chimpanzee; R. Rudicell for amplifying the SIVcpz*Pts* TAN10 *env* gene; P. Hallberg, S. Tian and the staff of the Penn Cytomics and Cell Sorting Resource Laboratory for assistance with B cell sorting; T. Denny, T. Demarco, and N. DeNaeyer and members of the Nonhuman Primate Virology Core Laboratory at the Duke Human Vaccine Institute for SCIV plasma vRNA measurements; R. Kaufman, H. Chen, and E. Lee for recombinant protein production and biochemical assays; the staff at Bioqual for exceptional care and assistance with nonhuman primates; and K. Ruffin for preparation of the manuscript.

## Author contributions

F.B.R., R.M.R., G.M.S., and B.H.H. conceived, planned and executed the study; H.L. and R.S.R. designed the SCIV constructs; F.B.R, R.M.R., W.D., J.R., and K.C generated virus stocks and performed neutralization assays; R.M.R. and A.N.S. performed single cell B cell sorting; W.L, Y.L., and Y.P. cloned, expressed and purified monoclonal antibodies; D.W, R.H., G.E.H., E.M., G.S.S., I-T.T., R.A. and K.O.S produced SOSIP trimers and probes; K.M. and R.J.E generated and analyzed NSEM data; S.W., R.B., and A.J.R. performed sequencing and computational analyses, E.L., K.C., W.D., G.S.S., and S.D. processed blood samples and performed experiments; E.E.G, K.W., and B.T.K. performed evolutionary analyses; S.S.M. performed statistical analyses; A.N., D.W.C, L.V. and K.O.S. designed and conducted the knock-in mouse experiments; S.M.A performed BLI analyses; A.F., R.A., D.C.M., K.J.W., M.G.L., P.D.K., D.R.B., K.O.S., and B.F.H provided critical expertise and contributed reagents. F.B.R., R.M.R., G.M.S., and B.H.H. coordinated the contributions of all authors and wrote the manuscript.

## Data and materials availability

All data associated with this study are in the paper or supplementary materials. Newly generated viral and antibody sequences have been deposited in GenBank with accession numbers listed in Table S4.

