## Supplement material for "A germline-targeting chimpanzee SIV envelope glycoprotein elicits a new class of V2-apex directed cross-neutralizing antibodies"

**Supplementary Material**

| | 156 160 166 171 |
|---|---|
| **HIV-1 M CRF250** | EMKNCSFNVTTELRDKKKKEYSFF |
| CPZ.*Ptt*.LB7 | T.......T.........QT..L. |
| CPZ.*Ptt*.LB715 | DLR.....AG-DI.....RVH.L. |
| CPZ.*Ptt*.MB897 | VQY...............QV..L. |
| CPZ.*Ptt*.MB66 | D.Y.....A......R..QV..L. |
| CPZ.*Ptt*.CAM13 | Q....T..T.........QV..L. |
| CPZ.*Ptt*.GAB2 | D.R..T..T........QQI..L. |
| CPZ.*Ptt*.MT145 | D.R..............RQV..L. |
| CPZ.*Ptt*.EK505 | Q.Q...............QV..L. |
| **HIV-1 N DJO0131** | H.......I...IH....QA..L. |
| CPZ.*Ptt*.DP943.2 | MV.K.F..M.......Q.QV..L. |
| GOR.BPID1 | G.LT.N.....V.K...EQKQAL. |
| **HIV-1 P RBF168** | .LFK....T..V.K...QEQQAL. |
| GOR.CP2139 | DIY........V.K..TT.QQAL. |
| GOR.CP2135 | DIY........V.K...T.QQAL. |
| GOR.BQID2 | .VY..T.....V.K...SQQQAL. |
| **HIV-1 O RBF206** | P..K.E.....V.K..QE.KQAL. |
| CPZ.*Pts*.TAN2 | .IF.....Q...F.....QI..L. |
| CPZ.*Pts*.TAN3 | .LF....QQ...F.....QI..L. |
| CPZ.*Pts*.TAN1 | .VY.....Q...F.....QI..L. |
| CPZ.*Pts*.TAN10 | .VY.....Q...FK....QI..L. |
| CPZ.*Pts*.UG38 | KVY.....Q...F.....QI..L. |
| CPZ.*Pts*.TAN13 | KVL.....Q...F.....NI..L. |
| CPZ.*Pts*.BF1167 | ..YE.F..Q...F.....QI..L. |
| CPZ.*Pts*.ANT_Cot | QLQQ.N.TQK.GF..R.QNITGI. |
| MUS.11GAB | PVY.....Q...F.....QM..L. |
| MUS.1085.1_54 | P.F.....Q...F....RQM..L. |
| MUS.1085.4_12 | P.F.....Q...F....RQM..L. |
| ASC.RT11 | SLL.....M.PGFK.R.AHYWAP. |
| SMM.E660 | P.IG.K..M.GLK...RIEYNETW |
| SMM.92b | P.IG.Q..M.GLKK.Q.RQYNETW |
| SMM.FTq | .LVS.K..M.GLK.....EYSETW |
| AGM.TAN1 | NSSI...AMAGYR..V...YN.TW |
| MND2.M14 | .NRV.K..T.GLC..C.IEIKES. |
| LHO.7 | RNAE.QY...GLC..CRTEIKQS. |
| WRC.98CI | INHV.R..T.GMCK.CRIEIKES. |
| WRC.05GM | TNHV.R....GLC..C.QEIVES. |

**Fig. S1. Sequence conservation in the V2-apex bNAb core epitope among primate lentiviruses.** V2 loop amino acid sequences are aligned to an HIV-1 group M reference strain (CRF250), with potential N-linked glycosylation sites at positions 156 and 160 (HXB2 numbering), and positively charged residues at positions 166 through 171 (HXB2 numbering) highlighted in red (strands B and C of the V2 loop are underlined). SIV Env sequences are labeled as in Figure 1 and color-coded to indicate their species/subspecies origin: central chimpanzees (CPZ.*Ptt*), blue; western gorillas (GOR) green; eastern chimpanzees (CPZ.*Pts*), magenta; moustached monkeys (MUS) and red-tailed monkeys (ASC), brown; sooty mangabeys (SMM), African green monkeys (AGM), mandrills (MND2), L'Hoest's monkeys (LHO), and western red colobus (WRC), red. HIV-1 reference strains CRF250 (group M), DJ131 (group N), RBF168 (group O) and RBF206 (group P) are shown in black (see Table S4 for GenBank accession numbers of all HIV-1 and SIV Envs analyzed). Dots indicate sequence identity to HIV-1 CRF250, with conservation of core epitope residues and glycosylation sites highlighted in yellow.

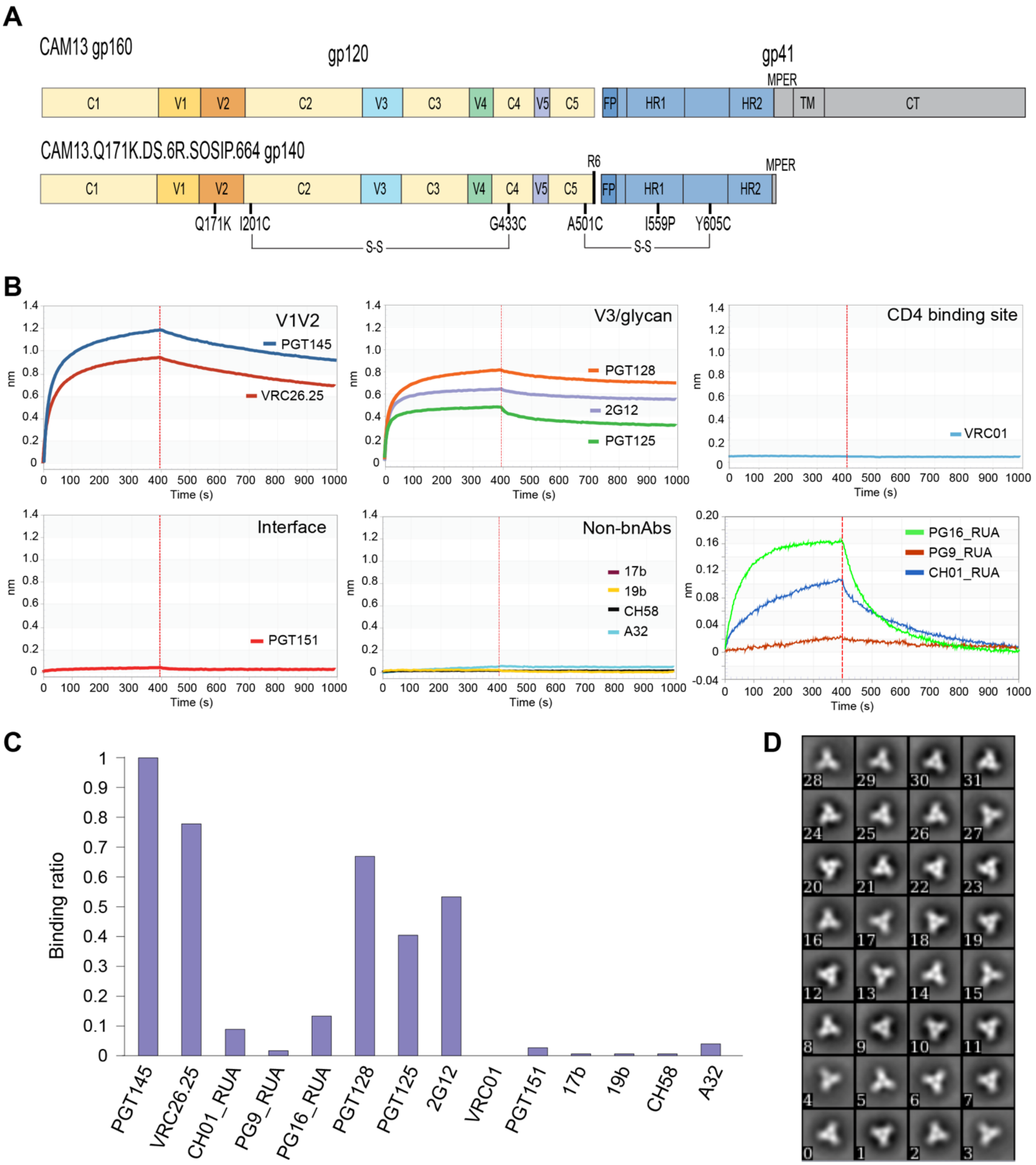


**Fig. S2. Generation and characterization of the CAM13K SOSIP trimer.** (A) The CAM13K SOSIP construction scheme is shown in relation to the wildtype SIVcpz CAM13 gp160, with the location of constant (C) and variable (V) gp120 segments as well as functional gp41 domains (FP, fusion peptide; HR1, heptad repeat 1; HR2, heptad repeat 2, MPER, membrane proximal external region; TM, transmembrane domain; CT, cytoplasmic domain) indicated. In addition to the Q171K mutation, the CAM13K SOSIP trimer was designed to include the original SOSIP.664 mutations (A501C-Y605C; I559P; truncation at residue 664) (*45, 46*), the DS stabilizing mutation (I201C-G433C) (*47*), as well as a hexa-arginine furin cleavage site (R6) (*48*). (B) Biolayer interferometry (BLI) sensorgrams depicting the binding of the CAM13K SOSIP trimer (at 50 μg/ml) to anti-human IgG Fc captured mature V2-apex (PGT145, VRC26.25), V3-glycan (PGT128, 2G12, PGT125), CD4 binding site (VRC01) and interface (PGT151) bNAbs as well as non-neutralizing V2 (CH58), V3 (19b), and CD4-induced (17b, A32) antibodies. Sensorgrams depicting the binding of the V2-apex bNAb precursors PG16_RUA, PG9_RUA, and CH01_RUA are also shown. (C) Antibody binding of the CAM13K SOSIP trimer relative to PGT145. Binding ratios were calculated relative to the binding of PGT145 which showed the highest level of binding (ratio = binding to mAb [nm] / binding to PGT145 [nm]). (D) Two-dimensional class averages of NSEM images of the CAM13K SOSIP protein showing the expected trimer morphology.

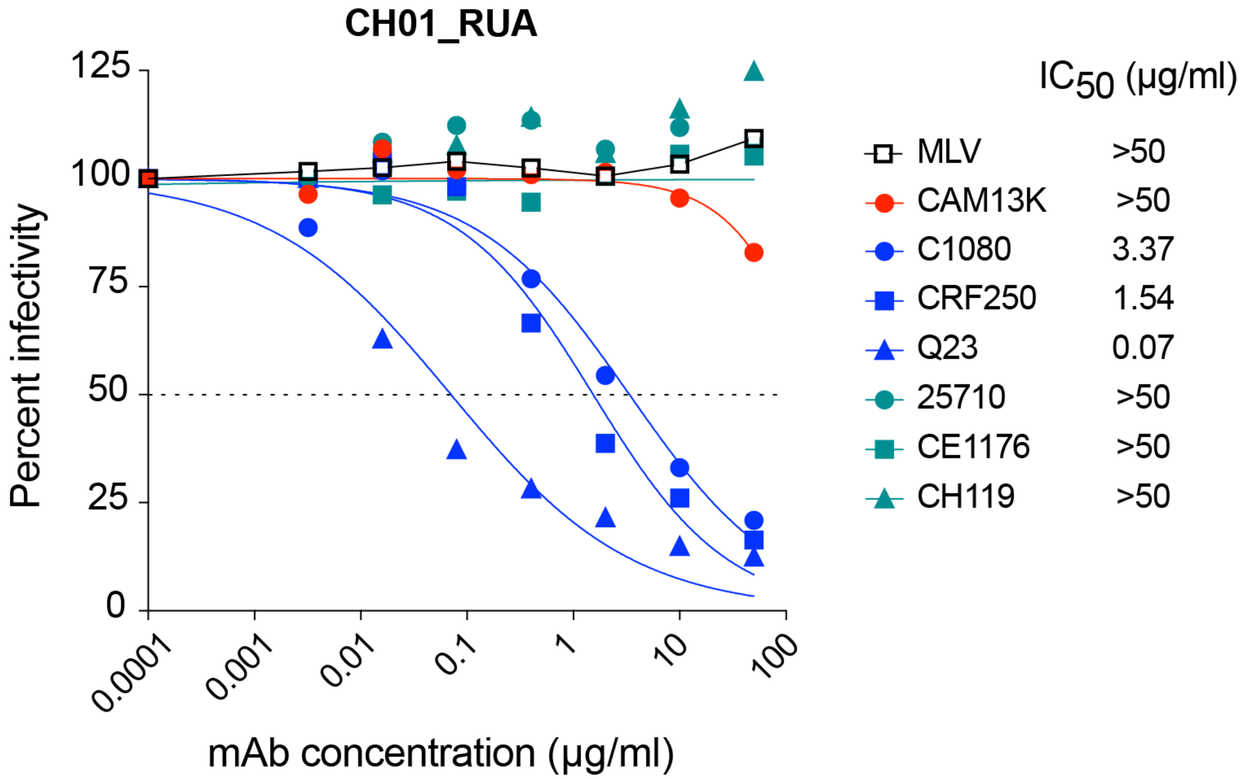


**Fig. S3. Sensitivity of HIV-1 Env pseudoviruses to neutralization by the human V2-apex precursor CH01_RUA.** Pseudoviruses carrying different HIV-1 Envs (C1080, CRF250, Q23, 25710, CE1176, CH119) were tested for their sensitivity to neutralization by the CH01_RUA precursor. The dashed line indicates 50% reduction in virus infectivity and the corresponding 50% inhibitory concentrations ($IC_{50}$ values in μg/ml) are listed on the right. HIV-1 pseudotypes sensitive to CH01_RUA are colored in blue, while resistant strains are shown in green. CAM13K (red) and MLV (black) pseudotypes are shown for control.

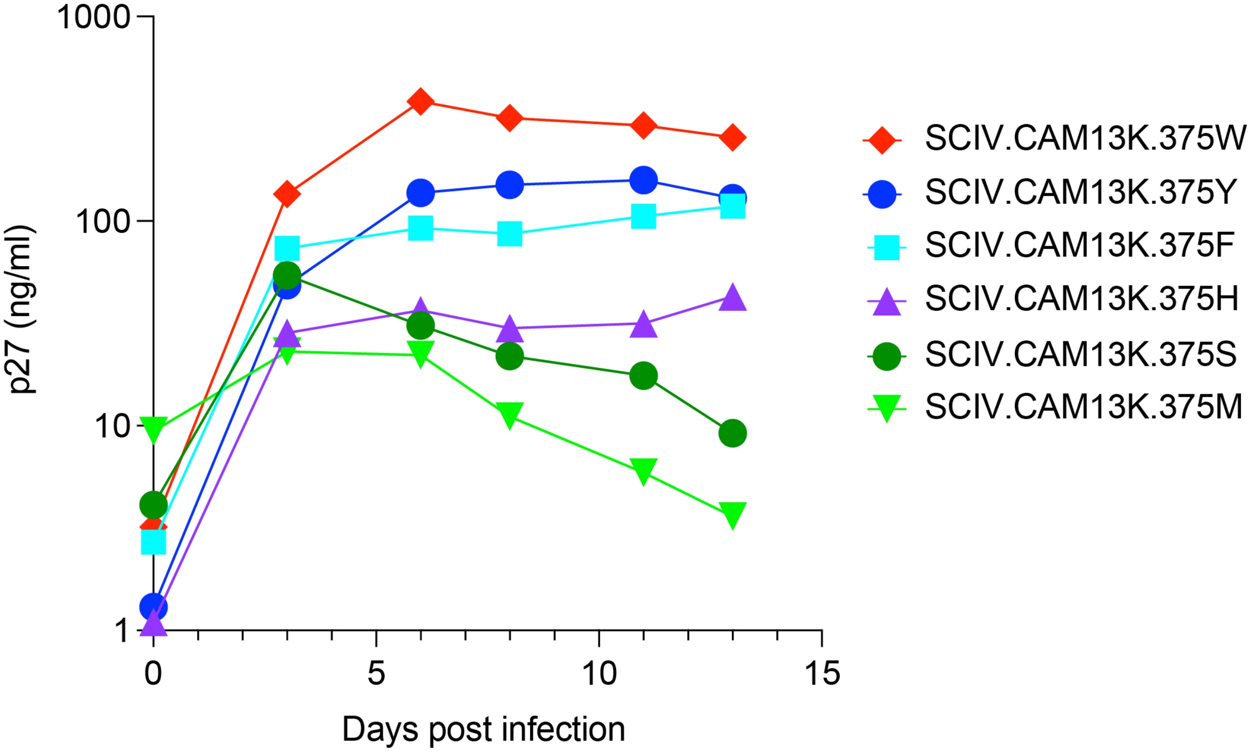


**Fig. S4. Replication kinetics of SCIV.CAM13K allelic variants *in vitro*.** Six isogenic mutants of SCIV.CAM13K differing only in the amino acid residue at Env position 375 were used to infect primary activated rhesus CD4+ T cells. Cultures were sampled at the indicated time points and virus replication was monitored by measuring p27 antigen content in the supernatant. SCIV.CAM13K.375W, which grew to the highest titers *in vitro*, also outcompeted all other mutants in infected rhesus macaques *in vivo*.

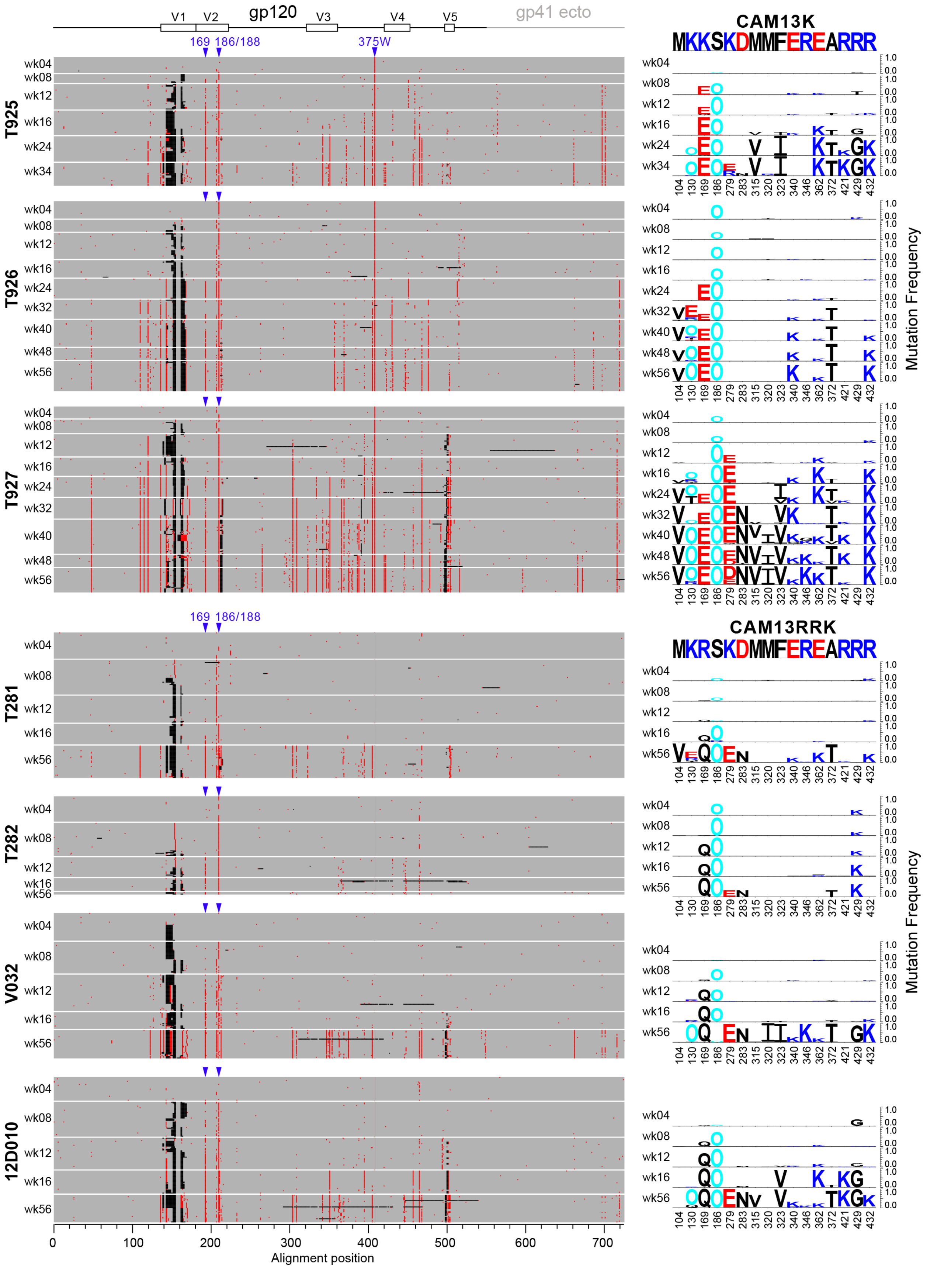
gp120
V1
V2
V3
V4
V5
gp41 ecto
169
186/188
375W
T925
T926
T927
T281
T282
V032
12D010
wk04
wk08
wk12
wk16
wk24
wk32
wk34
wk40
wk48
wk56
Alignment position
0
100
200
300
400
500
600
700
CAM13K
MKKSKDMMFEREARRR
CAM13RRK
MKRSKDMMFEREARRR
104
130
169
186
279
283
315
320
323
340
346
362
372
421
429
432
Mutation Frequency
1.0
0.0

**Fig. S5. Env sequence evolution in SCIV-infected rhesus macaques.** A Highlighter plot of longitudinal Env amino acid sequences obtained by single genome sequencing of plasma viral RNA is shown on the left for each infected RM. Sequences are grouped for each time point (SCIV.CAM13K infected animals are shown on the top, SCIV.CAM13RRK infected animals are shown on the bottom). Each row within a time block represents a single sequence, depicted as a string of horizontal pixels. Thus, each pixel represents a single amino acid in the alignment, which is colored grey if it matches the infecting virus, red if it represents a mutated residue, and black if it is inserted or deleted relative to the infecting virus. Sequences are shown up to HXB2 site 683 (gp41 ectodomain), with a schematic map depicted at the top and an alignment position scale depicted at the bottom. A single alignment was generated from sequences of all RMs and all time points, so that perfectly vertical "stripes" indicate the same HXB2 site in different Envs. One of these stripes represents the outgrowth of the 375W variant from the initial inoculum of six isogenic SCIV.CAM13K mutants (indicated on the top). Positions 169 and 186/188, which are mutated in all animals, are also highlighted. The right panels show mutation frequencies for each time point and RM at sites identified by LASSIE to be selected in at least two rhesus macaques (also see Table S2; sites beyond HXB2 position 683 are not shown). Mutations are depicted as logos, where the height of the letter is proportional to its frequency in the Env sequences from the corresponding time point (amino acids of the infecting virus are blanked out). Red, dark blue, and black indicate acidic, basic, and neutral residues, respectively. Cyan colored "O" indicates asparagine (N) embedded in an N-linked glycosylation motif (Asn-X-Ser or Asn-X-Thr, where X can be any amino acid except Pro). Numbers at the bottom indicate residue positions (HXB2 numbering).

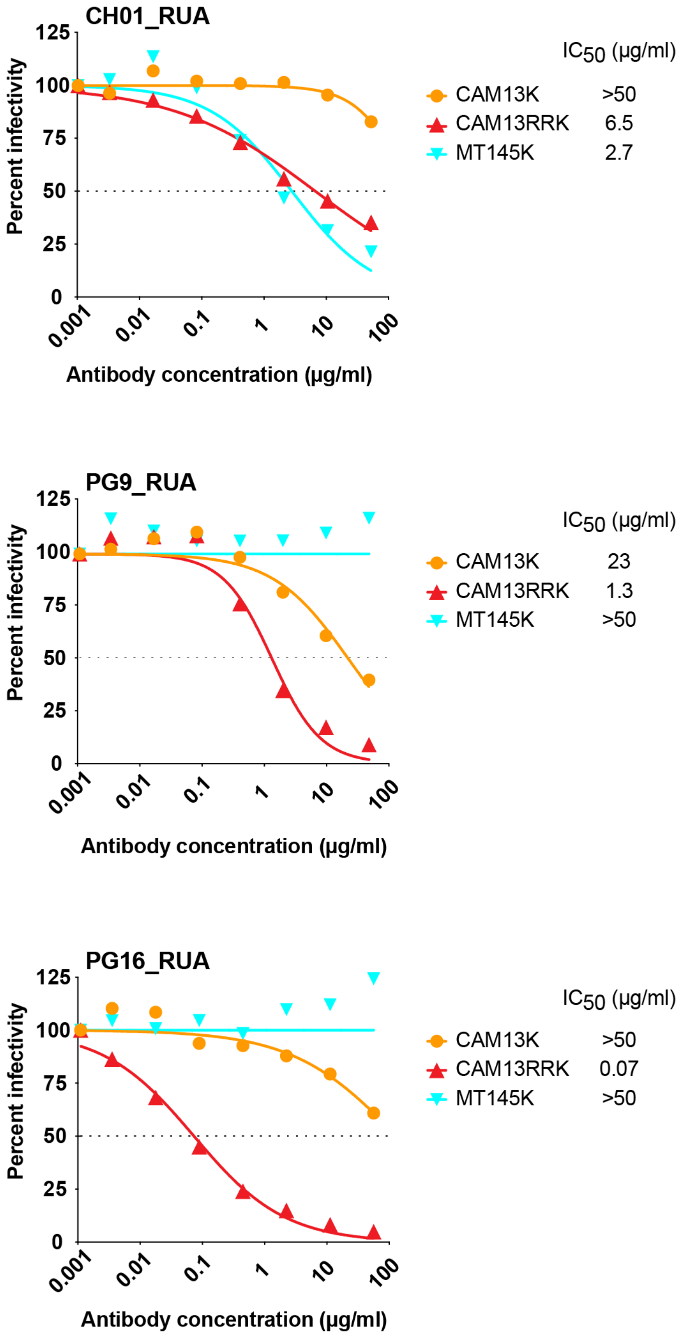


**Fig. S6. Sensitivity of CAM13RRK to neutralization by V2-apex bNAb precursors.** Neutralization curves are shown for human V2-apex bNAb precursors (indicated on top) against CAM13K (CAM13_Q171K), CAM13RRK (CAM13_Q171K_K169R_K170R) and MT145K (MT145_Q171K) Env bearing pseudoviruses, all of which encode the wildtype amino acid at position 375. The dashed line indicates 50% reduction in virus infectivity and the corresponding 50% inhibitory concentrations ($IC_{50}$ values in μg/ml) are listed on the right.

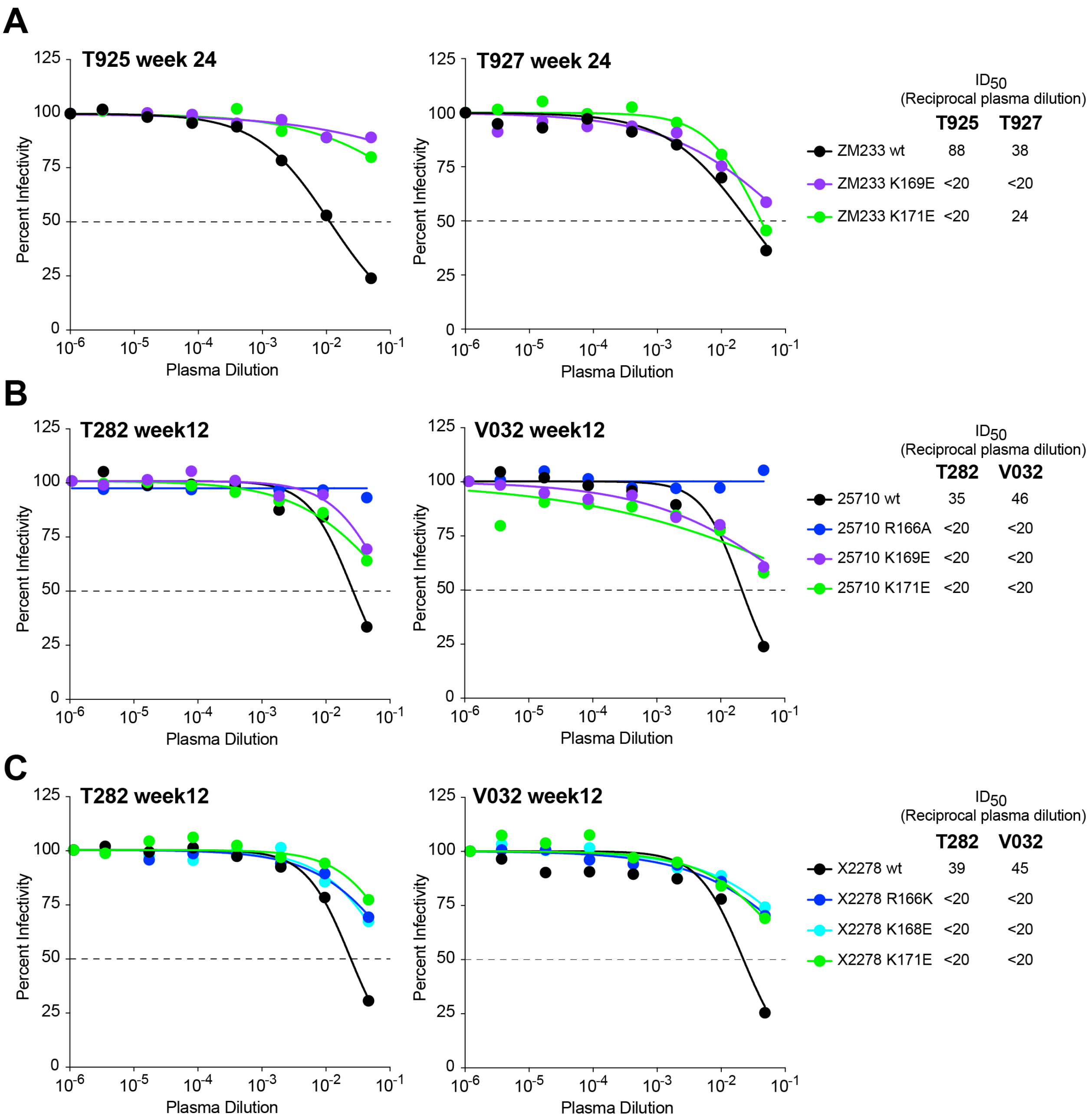


**Fig. S7. Low-titer heterologous plasma breadth in SCIV-infected RMs maps to the V2-apex.** Neutralization curves are shown for plasma samples from four RMs that developed low-titer heterologous neutralization breadth (also see Fig. 3). Site-directed mutants of ZM233, 25710 and X2278 Envs (indicated on the right) show that changes at residues 166, 169 and/or 171 reduce plasma neutralization, indicating a V2-targeted response. Corresponding $ID_{50}$ values are shown on the right.

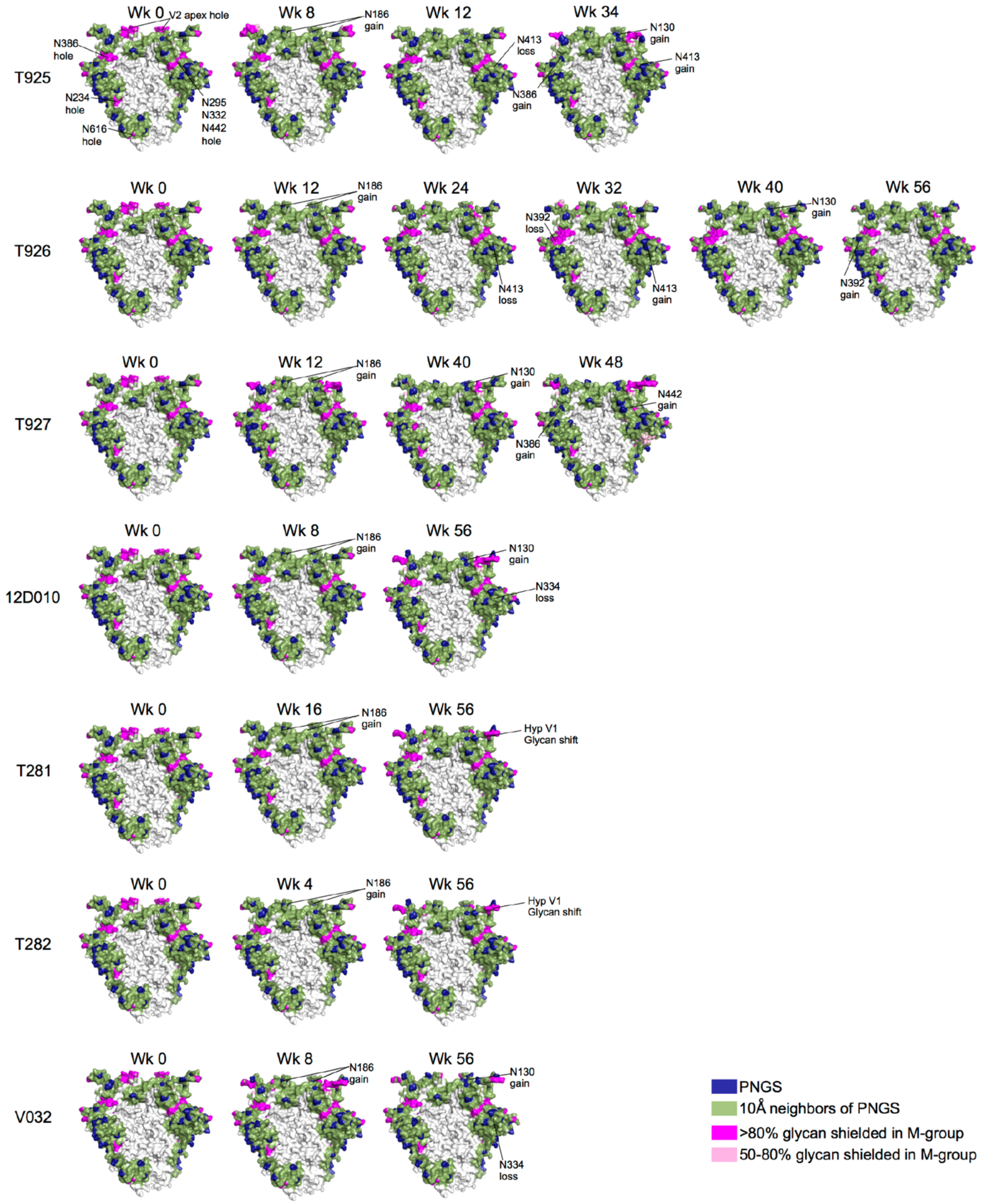


**Fig. S8. Glycan shield evolution in SCIV.CAM13K and SCIV.CAM13RRK infected RMs.** For each RM (indicated on the left), the evolution of the glycan shield is shown, starting with the Env of the transmitted virus and followed by glycan shield calculations of subsequent time points using time-point consensus glycans. The glycan shield is mapped on to a trimeric Env structure oriented with the V2 apex at the top and the viral membrane at the bottom (not shown). Blue regions indicate the location of potential N-linked glycosylation sites, green regions indicate predicted glycan shield coverage, and pink or magenta indicate rare “glycan holes” that are shielded in more than 50% or 80% M-group Envs, respectively. Glycan shield calculations were performed using the Glycan Shield Mapping webtool on the Los Alamos HIV Database (https://www.hiv.lanl.gov/content/sequence/GLYSHIELDMAP/glyshieldmap.html)with consensus glycans (50% or higher frequency for the time point) and hypervariable loop lengths chosen for longitudinal Envs as previously described (*56*). Glycan gains, losses and shifts are indicated.

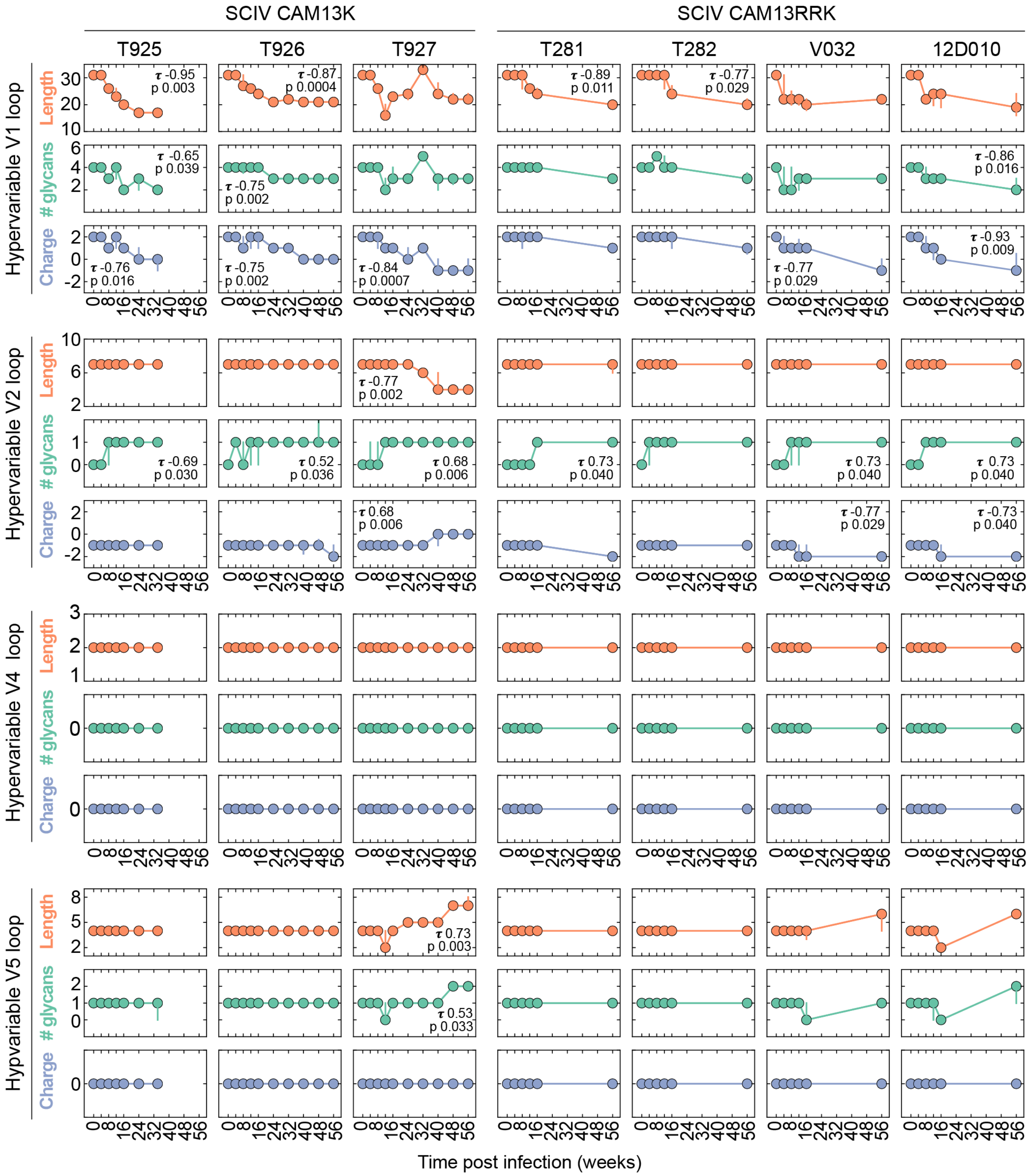


**Fig. S9. Env hypervariable loop evolution in SCIV.CAM13K and SCIV.CAM13RRK infected RMs.** The longitudinal evolution of SCIV.CAM13K and SCIV.CAM13RRK hypervariable loops was examined using alignment-free characteristics, including length, number of glycans and net charge, so as to not bias results by insertion and deletion-related alignment ambiguities. Each

RM is depicted as a column and hypervariable loop characteristics are depicted in rows. Individual plots show the longitudinal evolution of the indicated hypervariable loop characteristic (on the left), with time since infection (in weeks) indicated on the bottom. Points show the median of the characteristic calculated using Env sequences from the respective time point, with vertical bars spanning the inter-quartile range ($25^{th}$ to $75^{th}$ percentiles). Hypervariable loop net charge is measured in amino acid charge units, where His, Lys and Arg have +1 charge, and Asp and Glu have -1 charge. To measure statistical significance of longitudinal trends, the median values for each hypervariable loop characteristic from each time point and RM were subjected to a non-parametric Kendall-Tau test for associations with time post infection; for statistically significant ($p < 0.05$) associations, the Kendall Tau ($\tau$) coefficient and p-value are indicated in the plot. Hypervariable loop positions in HXB2 are: 132-152 for hypervariable V1, 185-190 for hypervariable V2, 396-410 for hypervariable V4 and 460-465 for hypervariable V5. Note that V3 does not have a hypervariable region. All calculations were performed using the Variable Region Characteristics tool (https://www.hiv.lanl.gov/content/sequence/VAR_REG_CHAR/index.html) available at the Los Alamos HIV Database.

| mAb | MLV | Tier 1A: MW965.26 | Tier 1A: TH023.6 | Global Neutralization Panel: TRO11 | 25710 | CNE8 | X2278 | BJOX010000 | X1632 | Ce1176 | 246F3 | CH119 | Ce0217 | CNE55 | SHIV Panel: Q23 | CRF250 | WITO | ZM233 | CAP256SU | CH505 | CH848 | BG505 N332* | SIVcpz: MT145K |
|---|---|---|---|---|---|---|---|---|---|---|---|---|---|---|---|---|---|---|---|---|---|---|---|
| **T927 lineage 1** | | | | | | | | | | | | | | | | | | | | | | | |
| w32 P2D12 | >250 | >250 | 6.6 | >250 | >250 | >250 | >250 | >250 | >250 | >250 | >250 | >250 | >250 | >250 | >250 | >250 | >250 | >250 | >250 | >250 | >250 | >250 | >250 |
| **T927 lineage 2** | | | | | | | | | | | | | | | | | | | | | | | |
| w24 P1D08 | >250 | 0.04 | >250 | >250 | >250 | >250 | >250 | >250 | >250 | >250 | >250 | >250 | >250 | >250 | >250 | >250 | >250 | 194 | >250 | >250 | >250 | >250 | >250 |
| w24 P2D06 | >250 | 0.02 | >250 | >250 | >250 | >250 | >250 | >250 | >250 | >250 | >250 | >250 | >250 | >250 | >250 | >250 | >250 | 119 | >250 | >250 | >250 | >250 | >250 |
| w24 P1D09 | >250 | 0.02 | >250 | >250 | >250 | >250 | >250 | >250 | >250 | >250 | >250 | >250 | >250 | >250 | >250 | >250 | >250 | 119 | >250 | >250 | >250 | >250 | >250 |
| **T927 lineage 3** | | | | | | | | | | | | | | | | | | | | | | | |
| w62 P1C05 | >250 | 0.05 | 0.80 | >250 | 59 | >250 | >250 | >250 | >250 | >250 | >250 | >250 | >250 | >250 | >250 | >250 | >250 | 82 | >250 | >250 | >250 | >250 | >250 |
| w62 P1C01 | >250 | 0.04 | 6.9 | >250 | 20.2 | >250 | >250 | >250 | >250 | >250 | >250 | >250 | >250 | >250 | >250 | >250 | >250 | 40.5 | >250 | >250 | >250 | >250 | >250 |
| **T925 lineage 2** | | | | | | | | | | | | | | | | | | | | | | | |
| w24 P1G03 | >250 | | | | >250 | | | | | | | | | | >250 | | | >250 | | | | | |
| w24 P1E11 | >250 | | | | >250 | | | | | | | | | | >250 | | | >250 | | | | | |
| w24 P1F01 | >250 | | | | >250 | | | | | | | | | | >250 | | | >250 | | | | | |
| w24 P2C12 | >250 | | | | >250 | | | | | | | | | | >250 | | | >250 | | | | | |
| w24 P1F12 | >250 | | | | >250 | | | | | | | | | | >250 | | | >250 | | | | | |

$IC_{50}$ 250-50µg/ml 49.9-10.0µg/ml 9.9-1.0µg/ml 0.99-0.01µg/ml

**Fig. S10. Expanded antibody lineages isolated from SCIV.CAM13K infected RMs with no or very limited tier 2 neutralization breadth.** 50% inhibitory concentrations ($IC_{50}$) are shown (µg/ml) for representative members of T927 lineages 1-3 and T925 lineage 2 (listed on the left) against pseudovirus panels carrying HIV-1 and SIVcpz Envs as well as replicating SHIVs (indicated on top). The highest antibody concentration used was 250 µg/ml (coloring indicates the relative neutralization potency).

| mAb | CAM13K | CAM13K-K130N | CAM13K-N156K | CAM13K-N160K | CAM13K-R166S | CAM13K-K169E | CAM13K-K171Q | CAM13K-S186N |
|---|---|---|---|---|---|---|---|---|
| **T927 lineage 1** | | | | | | | | |
| w32 P2D12 | 188 | 140 | 2.3 | 141 | 87 | 162 | >250 | 233 |
| **T927 lineage 2** | | | | | | | | |
| w24 P1D08 | 11.5 | 56 | 0.05 | 0.36 | >250 | >250 | >250 | 40.4 |
| w24 P2D06 | 9.2 | 34.6 | 0.03 | 0.12 | >250 | >250 | 174 | 30.1 |
| w24 P1D09 | 10.1 | 12.4 | 0.01 | 0.57 | >250 | >250 | 78 | 27.4 |
| **T927 lineage 3** | | | | | | | | |
| w62 P1C05 | >250 | >250 | 9.5 | >250 | >250 | >250 | >250 | >250 |
| w62 P1C01 | >250 | >250 | 11.4 | >250 | >250 | >250 | >250 | >250 |
| **T927 lineage 4** | | | | | | | | |
| w62 P1B01 | >250 | >250 | 6.9 | >250 | >250 | >250 | >250 | >250 |
| w62 P1B10 | >250 | >250 | 17.1 | >250 | n.d. | >250 | >250 | >250 |
| w62 P3E08 | >250 | >250 | 7.2 | 170 | 165 | 169 | >250 | >250 |
| w62 P1H10 | >250 | >250 | 19.0 | >250 | >250 | >250 | >250 | >250 |
| w62 P1F05 | >250 | >250 | 6.1 | >250 | >250 | >250 | >250 | >250 |
| w62 P3C02 | 215 | 142 | 1.7 | 141 | 110 | 231 | 233 | 198 |
| w62 P1C07 | >250 | >250 | 5.0 | >250 | >250 | >250 | >250 | >250 |
| w62 P3A11 | >250 | >250 | 5.8 | >250 | >250 | >250 | >250 | >250 |
| **T927 lineage 5** | | | | | | | | |
| w24 P2B12 | >250 | >250 | 50.0 | >250 | >250 | >250 | >250 | >250 |
| w24 P1C02 | >250 | >250 | 41.2 | >250 | >250 | >250 | >250 | >250 |
| w24 P1D03 | 100 | >250 | 5.8 | >250 | >250 | >250 | >250 | >250 |
| w24 P1A11 | 13.6 | 100 | 1.4 | >250 | >250 | >250 | >250 | 48.0 |
| w62 P1B05 | 157 | >250 | 1.3 | >250 | >250 | >250 | >250 | >250 |
| w62 P3G11 | 161 | >250 | 1.2 | >250 | >250 | >250 | >250 | >250 |
| **T925 lineage 1** | | | | | | | | |
| w24 P1E09 | 1.2 | 3.3 | 0.04 | >250 | >250 | >250 | >250 | 6.6 |
| w24 P1D09 | 1.1 | 6.8 | 0.09 | >250 | >250 | >250 | >250 | 4.6 |
| w24 P2B11 | 2.3 | 14.0 | 0.10 | 102 | >250 | >250 | >250 | 5.9 |
| w24 P1A12 | 0.6 | 4.4 | 0.05 | 219 | >250 | >250 | >250 | 3.7 |
| w24 P1C08 | 6.6 | 48.3 | 0.46 | >250 | >250 | >250 | >250 | 45.4 |
| **T925 lineage 2** | | | | | | | | |
| w24 P1G03 | >250 | | | | | | | |
| w24 P1E11 | >250 | | | | | | | |
| w24 P1F01 | >250 | | | | | | | |
| w24 P2C12 | >250 | | | | | | | |
| w24 P1F12 | >250 | | | | | | | |

| mAb | ZM233 | ZM233-N160K | ZM233-R166S | ZM233-K168E | ZM233-K169E | ZM233-K171E | MW965.26 | MW965.26-N160K | MW965.26-R166A | MW965.26-K168E | MW965.26-K169E | MW965.26-Q171K |
|---|---|---|---|---|---|---|---|---|---|---|---|---|
| **T927 lineage 5** | | | | | | | | | | | | |
| w24 P2B12 | 14.0 | 48.0 | >250 | >250 | >250 | >250 | 1.4 | >10 | >10 | >10 | >10 | 0.91 |
| w24 P1C02 | 68 | 61 | >250 | >250 | >250 | >250 | 3.5 | >10 | >10 | >10 | >10 | 1.0 |
| w24 P1D03 | 10.1 | 21.3 | >50 | >50 | >50 | >50 | 0.76 | 6.8 | >10 | >10 | >10 | 0.22 |
| w24 P1A11 | 0.52 | 1.2 | >50 | >50 | >50 | >50 | 0.09 | 0.73 | >10 | >10 | >10 | 0.01 |
| w62 P1B05 | 0.62 | 5.3 | >50 | >50 | 46.0 | >50 | 0.01 | 0.30 | >10 | >10 | 0.01 | 0.003 |
| w62 P3G11 | 2.0 | 5.3 | >50 | >50 | >50 | >50 | 0.14 | 2.5 | >10 | >10 | 0.37 | 0.02 |
| **T925 lineage 1** | | | | | | | | | | | | |
| w24 P1E09 | 0.78 | 9.5 | 17.8 | >50 | >50 | >50 | 4.0 | >10 | >10 | >10 | >10 | 0.007 |
| w24 P1D09 | 5.0 | 2.5 | >50 | >50 | >50 | >50 | 9.0 | >10 | >10 | >10 | >10 | 0.009 |
| w24 P2B11 | 0.77 | 18.7 | 17.0 | >50 | >50 | >50 | 4.9 | >10 | >10 | >10 | >10 | 0.02 |
| w24 P1A12 | 0.28 | 13.0 | 13.0 | >50 | >50 | >50 | 22.9* | >10 | >10 | >10 | >10 | 0.006 |
| w24 P1C08 | 2.0 | 9.7 | >50 | >50 | >50 | >50 | >10 | >10 | >10 | >10 | >10 | 0.05 |
| **T925 lineage 2** | | | | | | | | | | | | |
| w24 P1G03 | >250 | | | | | | | | | | | |
| w24 P1E11 | >250 | | | | | | | | | | | |
| w24 P1F01 | >250 | | | | | | | | | | | |
| w24 P2C12 | >250 | | | | | | | | | | | |
| w24 P1F12 | >250 | | | | | | | | | | | |

$IC_{50}$: 250-50µg/ml; 49.9-10.0µg/ml; 9.9-1.0µg/ml; 0.99-0.01µg/ml; <0.01µg/ml

**Fig. S11. SCIV-induced cross-neutralizing antibodies target the V2 region.** Representative members of all isolated antibody lineages (listed on the left) were tested against panels of CAM13K, ZM233 and/or MW965.26 mutants (indicated on top) to map their respective epitopes. Given the patterns of early sequence evolution, we mutagenized residues at or near the V2-apex, with an emphasis on key N-linked glycosylation sites and C strand residues. 50% inhibitory concentrations ($IC_{50}$) are shown (µg/ml). An asterisk denotes an estimated $IC_{50}$ value (as determined by the Prism software). Only the two most potent and cross-reactive lineages were tested against mutants of the heterologous ZM233 and MW965.26 Envs. All viruses represented pseudotypes generated by transfection of 293T cells. The highest antibody concentration used was 250 µg/ml (colors indicate the relative neutralization potency).

**A**

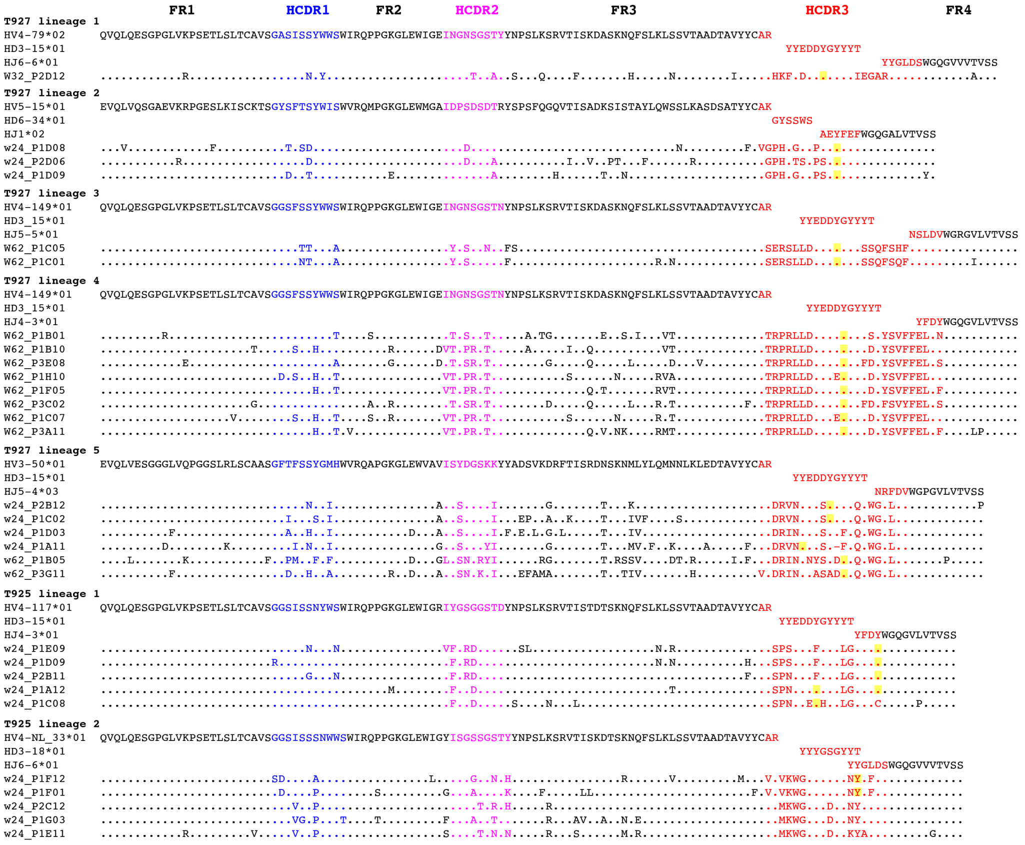
FR1
HCDR1
FR2
HCDR2
FR3
HCDR3
FR4
T927 lineage 1
HV4-79*02
HD3-15*01
HJ6-6*01
W32_P2D12
T927 lineage 2
HV5-15*01
HD6-34*01
HJ1*02
w24_P1D08
w24_P2D06
w24_P1D09
T927 lineage 3
HV4-149*01
HD3_15*01
HJ5-5*01
W62_P1C05
W62_P1C01
T927 lineage 4
HV4-149*01
HD3_15*01
HJ4-3*01
W62_P1B01
W62_P1B10
W62_P3E08
W62_P1H10
W62_P1F05
W62_P3C02
W62_P1C07
W62_P3A11
T927 lineage 5
HV3-50*01
HD3-15*01
HJ5-4*03
w24_P2B12
w24_P1C02
w24_P1D03
w24_P1A11
w62_P1B05
w62_P3G11
T925 lineage 1
HV4-117*01
HD3-15*01
HJ4-3*01
w24_P1E09
w24_P1D09
w24_P2B11
w24_P1A12
w24_P1C08
T925 lineage 2
HV4-NL_33*01
HD3-18*01
HJ6-6*01
w24_P1F12
w24_P1F01
w24_P2C12
w24_P1G03
w24_P1E11

**B**

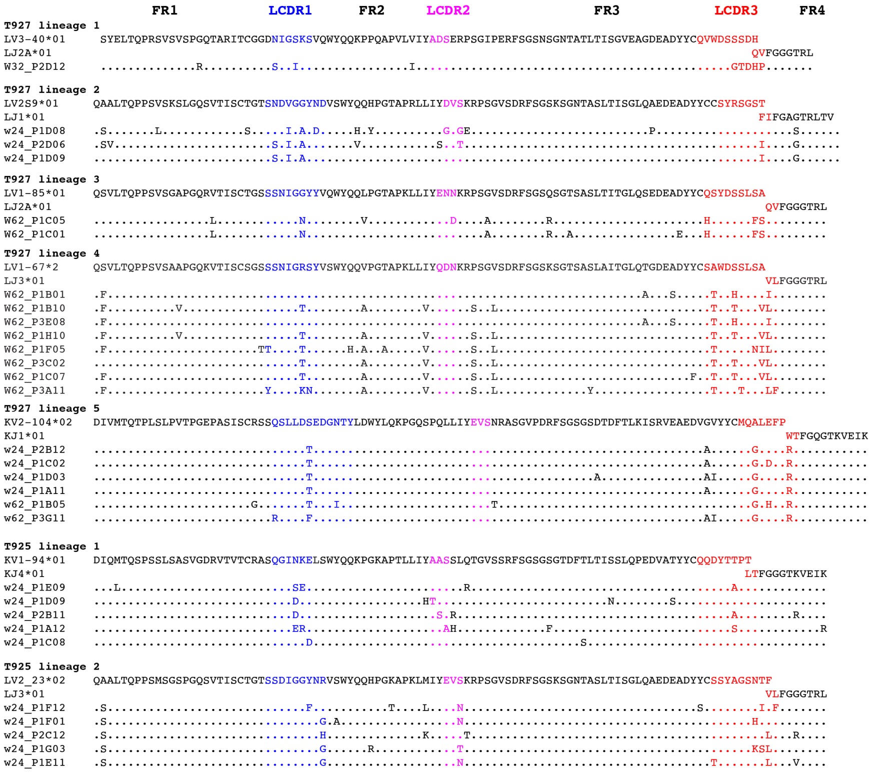
FR1
LCDR1
FR2
LCDR2
FR3
LCDR3
FR4
T927 lineage 1
LV3-40*01
LJ2A*01
W32_P2D12
T927 lineage 2
LV2S9*01
LJ1*01
w24_P1D08
w24_P2D06
w24_P1D09
T927 lineage 3
LV1-85*01
LJ2A*01
W62_P1C05
W62_P1C01
T927 lineage 4
LV1-67*2
LJ3*01
W62_P1B01
W62_P1B10
W62_P3E08
W62_P1H10
W62_P1F05
W62_P3C02
W62_P1C07
W62_P3A11
T927 lineage 5
KV2-104*02
KJ1*01
w24_P2B12
w24_P1C02
w24_P1D03
w24_P1A11
w62_P1B05
w62_P3G11
T925 lineage 1
KV1-94*01
KJ4*01
w24_P1E09
w24_P1D09
w24_P2B11
w24_P1A12
w24_P1C08
T925 lineage 2
LV2_23*02
LJ3*01
w24_P1F12
w24_P1F01
w24_P2C12
w24_P1G03
w24_P1E11

**Fig. S12. Heavy and light chain variable region sequences of SCIV-induced cross-neutralizing antibodies**. Heavy (H, panel A) and light (lambda [L] and kappa [K], panel B) chain variable region sequences of mature lineage members isolated from RMs T927 and T925 (listed on the left) are aligned to the deduced amino acid sequences of their respective germline variable (V), diversity (D), and joining (J) genes (indicated on the top). Framework (FR) and complementary determining regions (CDR) are color coded, with HCDR1 and LCDR1, HCDR2 and LCDR2, and HCDR3 and LCDR3 highlighted in blue, pink and red, respectively. Predicted sulfated tyrosines in the HCDR3 region (using the GPS-TSP 1.0 software at a high threshold setting) are highlighted in yellow (*98*). Expanded lineages (listed on the left) were identified using SONAR's automated "unseeded" lineage assignment tool, followed by manual inspection of gene assignments, HCDR3 regions, and non-templated nucleotides to confirm membership (Genbank accession numbers are listed in Table S4).

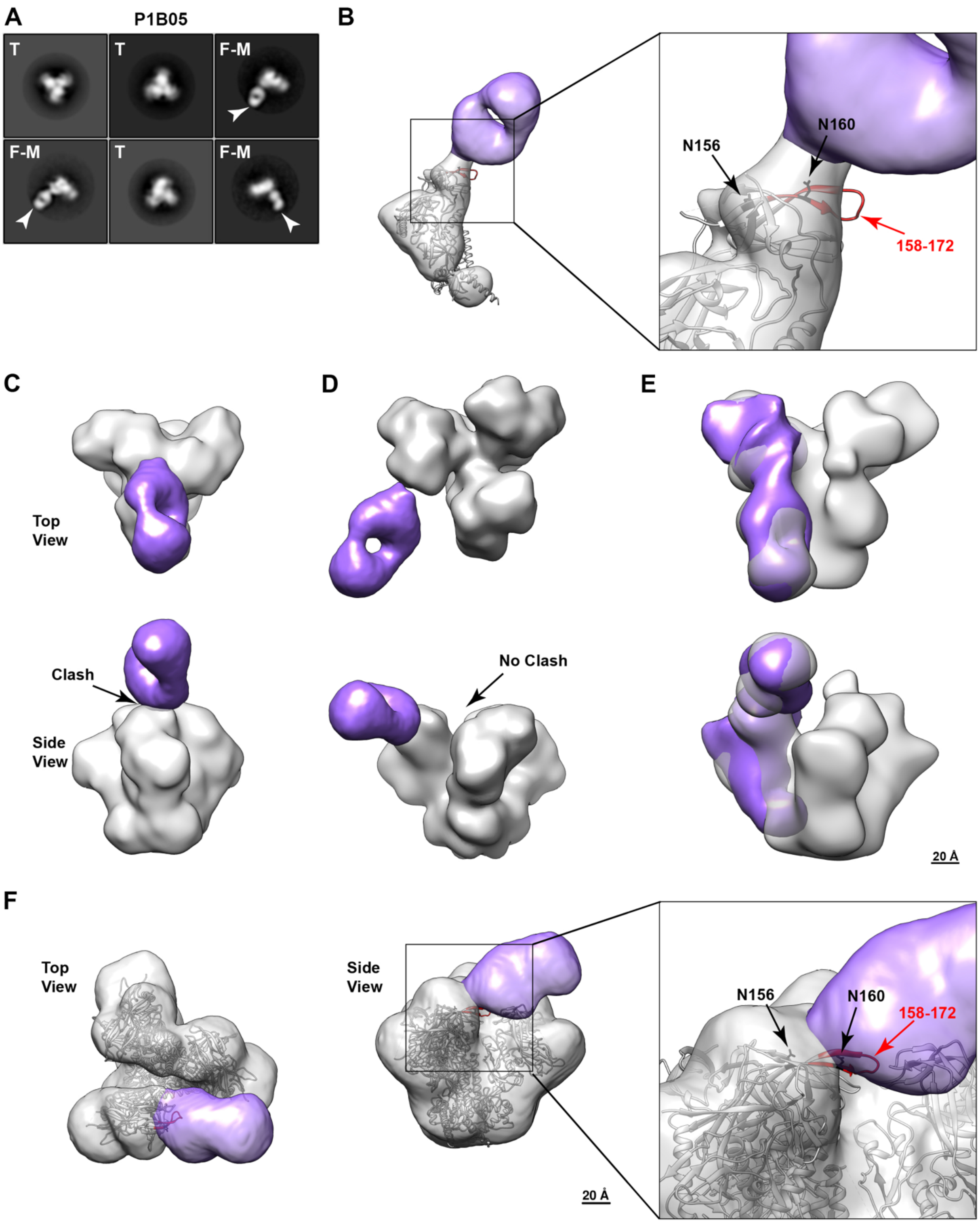

A
P1B05
T
T
F-M
F-M
T
F-M
B
N156
N160
158-172
C
Top View
Clash
Side View
D
No Clash
E
20 Å
F
Top View
Side View
20 Å
N156
N160
158-172


Figure S13

**Fig. S13. The SCIV-induced cross-neutralizing antibody P1B05 from RM T927 binds a V2 epitope that is occluded in a closed, prefusion trimer**. (A) 2D class averages showing intact SOSIP trimers without bound Fabs (T) and Fab-monomers (F-M), but no Fab-trimer complexes. Fabs in Fab-monomer complexes are indicated by an arrow. (B) 3D reconstruction of a P1B05 Fab-monomer fits the model of a single Env protomer (PDB 5FYL), with the Env domain in gray and the Fab in purple. A higher magnification inset identifies residues 158-172 in the V2 loop (highlighted in red) as the likely epitope (the positions of glycans at positions 156 and 160 are shown). (C) Modeling of P1B05 Fab binding onto a closed, prefusion trimer. Both top and side views are shown, revealing a clash (arrow) with an adjacent protomer. (D) Modeling of P1B05 Fab binding onto an occluded-open trimer identifies no clashes with adjacent protomers. (E) Overlay of Fab-monomer (colored) and Fab-trimer structures. The Fab-monomer fits well within the density of the corresponding trimer, thus validating the utility of Fab-monomer complexes to infer putative epitopes and approximate angles of approach. (F) P1B05 Fab-trimer structures identify the same putative V2 epitope as the corresponding Fab-monomer (residues 158-172; highlighted in red) shown in (B).

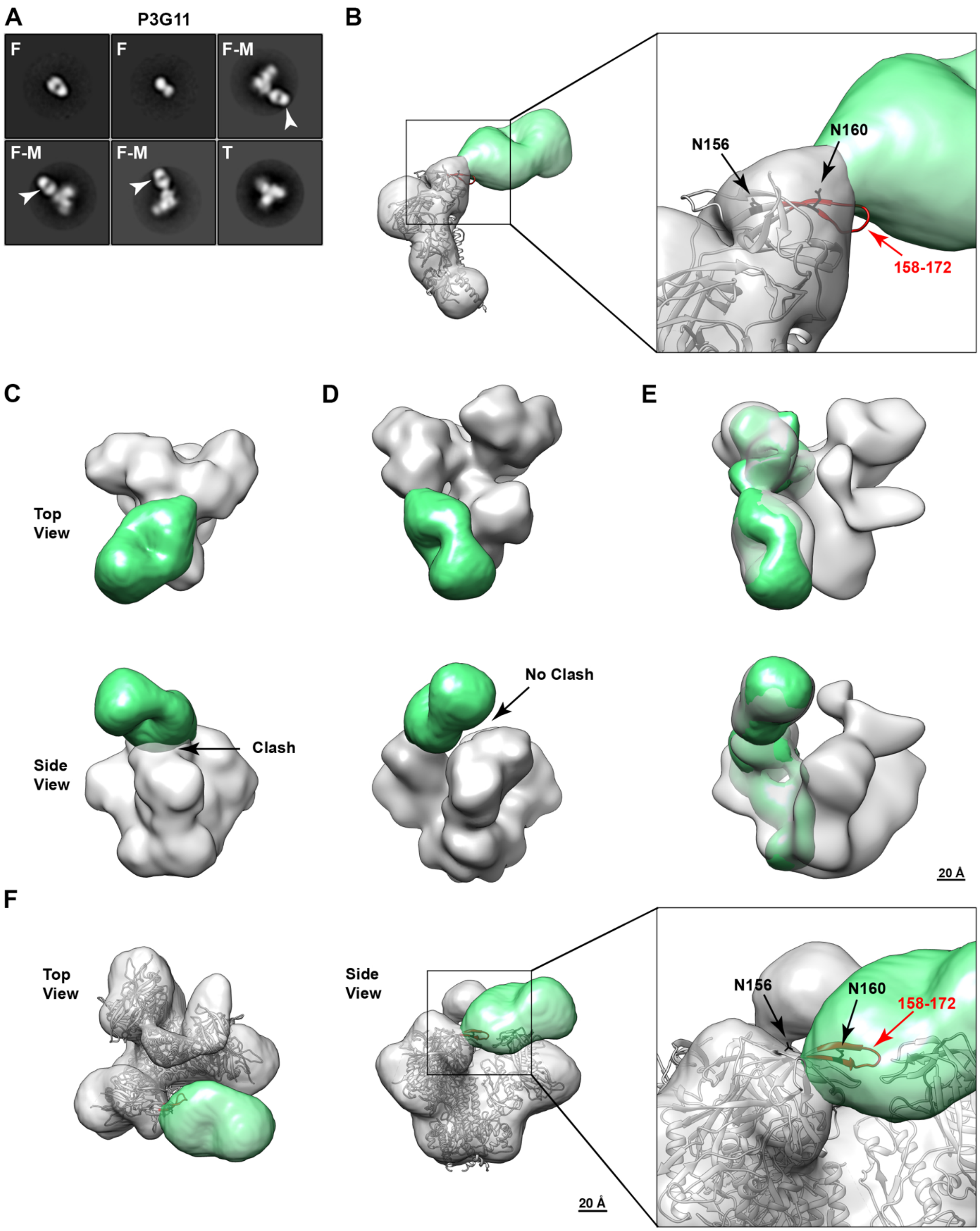


Figure S14

**Fig. S14. The SCIV-induced cross-neutralizing antibody P3G11 from RM T927 binds a V2 epitope that is occluded in a closed, prefusion trimer**. (A) 2D class averages showing Fabs

(F), am intact SOSIP trimer without a bound Fab (T) and Fab-monomers (F-M), but no Fab-trimer complexes. Fabs in Fab-monomer complexes are indicated by an arrow. (B) 3D reconstruction of a P3G11 Fab-monomer fits the model of a single Env protomer (PDB 5FYL), with the Env domain in gray and the Fab in green. A higher magnification inset identifies residues 158-172 in the V2 loop (highlighted in red) as the likely epitope (the positions of glycans at positions 156 and 160 are shown). (C) Modeling of P3G11 Fab binding onto a closed, prefusion trimer. Both top and side views are shown, revealing a clash (arrow) with an adjacent protomer. (D) Modeling of P3G11 Fab binding onto an occluded-open trimer identifies no clashes with adjacent protomers. (E) Overlay of Fab-monomer (colored) and Fab-trimer structures. The Fab-monomer fits well within the density of the corresponding trimer, thus validating the utility of Fab-monomer complexes to infer putative epitopes and approximate angles of approach. (F) P3G11 Fab-trimer structures identify the same putative V2 epitope as the corresponding Fab-monomer (residues 158-172; highlighted in red) shown in (B).

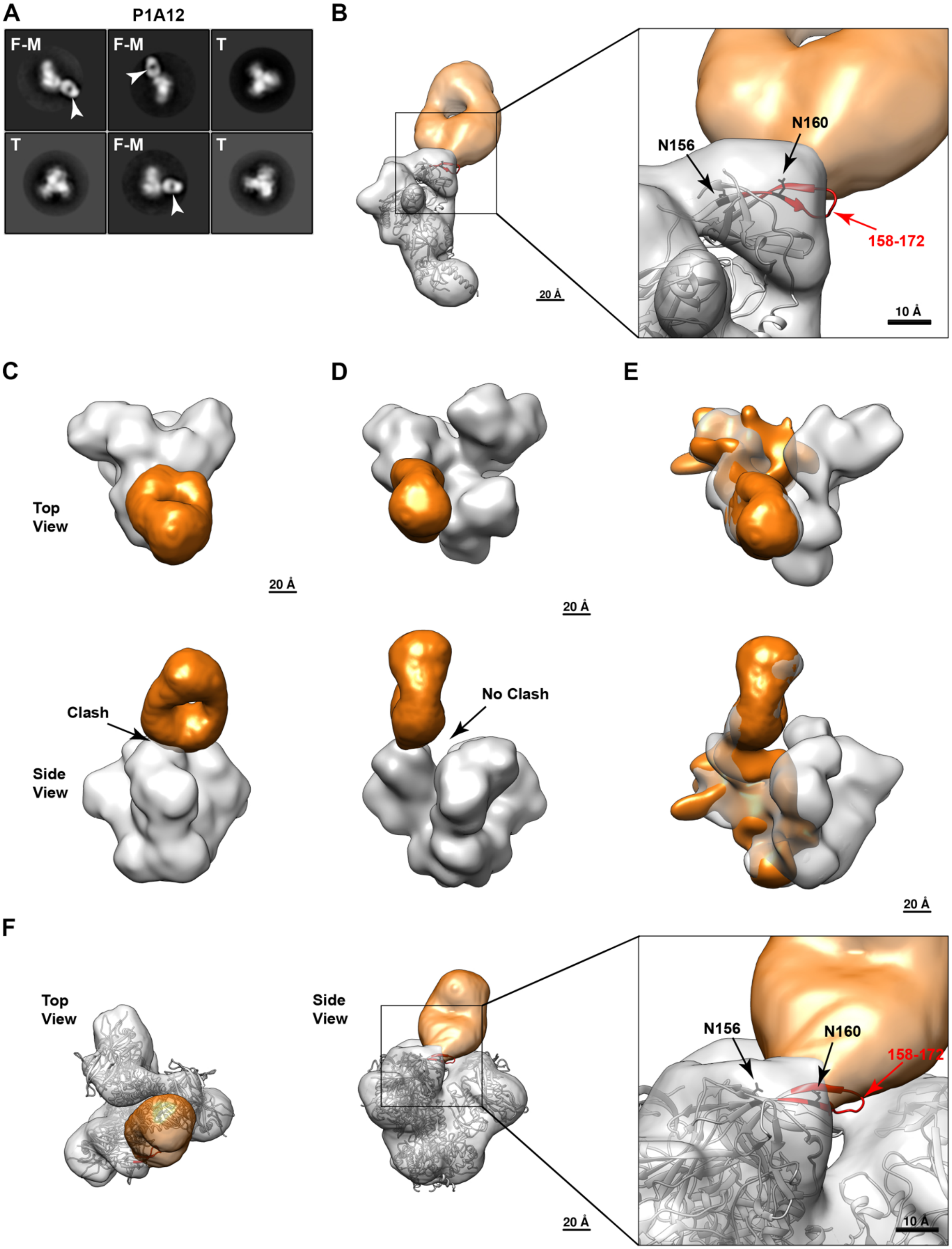
A
P1A12
F-M
F-M
T
T
F-M
T
B
N156
N160
158-172
20 Å
10 Å
C
Top View
20 Å
Clash
Side View
D
20 Å
No Clash
E
20 Å
F
Top View
Side View
N156
N160
158-172
20 Å
10 Å

Figure S15

**Fig. S15. The SCIV-induced cross-neutralizing antibody P1A12 from RM T925 binds a V2 epitope that is occluded in a closed, prefusion trimer**. (A) 2D class averages showing intact SOSIP trimers without a bound Fab (T) and Fab-monomers (F-M), but no Fab-trimer complexes. Fabs in Fab-monomer complexes are indicated by an arrow. (B) 3D reconstruction of a P1A12 Fab-monomer fits the model of a single Env protomer (PDB 5FYL), with the Env domain in gray and the Fab in gold. A higher magnification inset identifies residues 158-172 in the V2 loop (highlighted in red) as the likely epitope (the positions of glycans at positions 156 and 160 are shown). (C) Modeling of P1A12 Fab binding onto a closed, prefusion trimer. Both top and side views are shown, revealing a clash (arrow) with an adjacent protomer. (D) Modeling of P1A12 Fab binding onto an occluded-open trimer identifies no clashes with adjacent protomers. (E) Overlay of Fab-monomer (colored) and Fab-trimer structures. The Fab-monomer fits well within the density of the corresponding trimer, thus validating the utility of Fab-monomer complexes to infer putative epitopes and approximate angles of approach. (F) P1A12 Fab-trimer structures identify the same putative V2 epitope as the corresponding Fab-monomer (residues 158-172; highlighted in red) shown in (B).

**Table S1. Neutralization sensitivity of wildtype and modified SIV Envs to V2-apex bNAbs and their inferred precursors**

| Neutralization sensitivity[1] | PG9 RUA[2] | PG16 RUA[2] | VRC26 UCA[2] | CH01 RUA[2] | PG9 | PG16 | VRC26.25 | CH01 |
|---|---|---|---|---|---|---|---|---|
| SIVcpz.LB7 | >50 | >50 | >50 | >50 | >10 | 1.4 | 0.08 | >10 |
| SIVcpz.LB7 Q171K | >50 | >50 | >50 | >50 | 0.73 | >10 | 0.07 | 7.6 |
| SIVcpz.MB897 | >50 | >50 | >50 | >50 | 0.09 | 0.05 | 0.001 | >10 |
| SIVcpz.MB897 Q171K | >50 | >50 | >50 | >50 | 0.03 | 0.01 | 0.001 | 1.2 |
| SIVcpz.MB897 Q171K + N130K | >50 | >50 | >50 | >50 | 0.003 | 0.003 | 0.0007 | 0.18 |
| SIVcpz.MB66 | >50 | >50 | >50 | >50 | >10 | 0.01 | 0.07 | >10 |
| SIVcpz.MB66 Q171K | >50 | >50 | >50 | >50 | >10 | 5.8 | 0.04 | >10 |
| SIVcpz.CAM13 | >50 | >50 | >50 | >50 | 0.02 | 0.09 | 0.002 | >10 |
| SIVcpz.CAM13 Q171K | 28.2 | 37.7 | >50 | >50 | 0.007 | 0.003 | 0.002 | 0.09 |
| SIVcpz.MT145 | >50 | >50 | >50 | >50 | 0.16 | 0.34 | 0.0003 | >10 |
| SIVcpz.MT145 Q171K | >50 | >50 | >50 | 2.7 | 0.04 | 0.03 | 0.002 | 0.34 |
| SIVcpz.DP943.2 | >50 | >50 | >50 | >50 | 0.02 | 0.02 | 0.001 | 6.8 |
| SIVcpz.DP943.2 Q171K | >50 | >50 | >50 | 1.0 | 0.007 | 0.006 | 0.0007 | 0.06 |
| SIVcpz.DP943.2 Q171K + N156 glycan (K156N+F158S) | >50 | >50 | >50 | 49.1 | 0.01 | 0.009 | 0.009 | 0.15 |
| SIVgor.BPID1 | >50 | >50 | >50 | >50 | >10 | >10 | >10 | >10 |
| SIVgor.BPID1 E170K + Q171K | >50 | >50 | >50 | >50 | >10 | 4.3 | >10 | >10 |
| SIVgor.CP2139 | >50 | >50 | >50 | >50 | >10 | >10 | >10 | >10 |
| SIVgor.CP2139 T169K + T170K | >50 | >50 | >50 | >50 | 0.12 | 0.76 | >10 | 2.8 |
| SIVgor.CP2139 T169K + T170K + K166R | >50 | >50 | >50 | >50 | 0.14 | 8.1 | 0.008 | >10 |
| SIVgor.CP2139 T169K + T170K + K166R + N130K | >50 | >50 | >50 | >50 | 0.01 | 9.6 | 0.004 | >10 |
| SIVgor.BQID2 | >50 | >50 | >50 | >50 | >10 | 4.9 | >10 | >10 |
| SIVgor.BQID2 S170K + Q171K | >50 | >50 | >50 | >50 | 0.45 | 0.79 | >10 | >10 |
| SIVgor.BQID2 S170K + Q171K + N130T | >50 | >50 | >50 | >50 | 3.2 | >10 | >10 | >10 |
| SIVcpz.TAN10 | >50 | >50 | >50 | >50 | >10 | 1.9 | >10 | 5.6 |
| SIVcpz.TAN10 Q171K | >50 | >50 | >50 | >50 | 0.13 | >10 | >10 | 7.0 |
| SIVmus.11GAB | n.t | n.t | n.t | n.t. | 0.54 | 9.5 | 0.54 | >10 |
| SIVmus.11GAB Q171K | >50 | >50 | >50 | >50 | 0.12 | 1.8 | 0.92 | 6.5 |
| SIVmus.1085 4-12 | n.t | n.t | n.t | n.t. | 0.14 | 0.006 | 0.03 | >10 |
| SIVmus.1085 4-12 Q171K | >50 | >50 | >50 | >50 | 0.03 | 0.04 | 0.02 | 1.17 |
| SIVasc.RT11 | n.t | n.t | n.t | n.t. | 0.47 | >10 | >10 | >10 |
| SIVasc.RT11 H171K | >50 | >50 | >50 | >50 | 0.09 | >10 | >10 | 9.3 |
| SIVasc.RT11 H171K + K166R | >50 | >50 | >50 | >50 | 0.05 | 0.55 | >10 | 2.6 |
| SIVasc.RT11 H171K + A170K | >50 | >50 | >50 | >50 | 0.31 | >10 | >10 | >10 |

[1]Numbers indicating $IC_{50}$ values (μg/ml) as determined in the TZM-bl assay (n.t., not tested). The highest antibody concentration used was 50 μg/ml (colors indicate relative neutralization potency).
[2]RUA, reverted unmutated ancestor; UCA, unmutated common ancestor.

**Table S2. Env sites under selection in SCIV.CAM13K and SCIV.CAM13RRK infected rhesus macaques**

| Amino acid position | SCIV CAM13K/ CAM13RRK | RM T925 | RM T926 | RM T927 | RM T281 | RM T282 | RM V032 | RM 12D010 |
|---|---|---|---|---|---|---|---|---|
| **169** | K/R | **E** | **E** | **E** | **Q** | **Q** | **Q** | **Q** |
| **186/188** | S | **O** | **O** | **O** | **O**,R | **O** | **O**,S,D | **O** |
| **372** | A | **T,V** | **T** | **T,V** | **T** | | **T** | **T** |
| **130** | K | | **O,R**,E | **O,R**,T | | | **O** | **O,R**,Q |
| **279** | K | | | **E**,D | **E** | | **E** | **E** |
| **323** | F | **I** | | **I,**V | | | **I** | V |
| **432** | R | **K** | **K** | **K** | | | **K** | |
| **429** | R | G,T | | | | | **G** | **G** |
| **362** | E | **K** | | **K** | | | | **K** |
| **749*** | L | | **P**,R | **P** | **P** | | | |
| **830*** | H | | | | **Y,R** | | **Y**,L,**R**,Q,N | |
| **104** | M | | | **V** | **V** | | | |
| **283** | D | | | **N**,I | | | | **N** |
| **315** | M | | | V | | | | |
| **320** | M | | | **I** | | | **I** | |
| **340** | E | | **K** | **K,**G | | | | |
| **346** | R | | | **K**,G | | | **K** | |
| **421** | R | **K** | | | | | | **K** |

The Longitudinal Antigenic Sequences and Sites from Intra-Host Evolution (LASSIE) program was used to identify Env residues where mutations altered the encoded amino acid in at least 80% of sequences at one or more time points (O denotes potential N-linked glycosylation sites). Sites are sorted by the number of animals in which they were detected. Bold faced amino acids were found in two and more animals. An asterisk indicates positions outside of the SIVcpz CAM13 Env ectodomain in the SCIV gp41 region.

### Table S3. Varying nomenclature in different databases for the same rhesus macaque germline heavy (H) chain variable (V), diversity (D) and joining (J) gene sequences

| Segment | Bernat* | Ramesh# | IMGT^ | Sequence (5'-3') |
|---|---|---|---|---|
| HV | IGHV4-117*01_S5847 | | IGHV4-160*01 | CAGGTGCAGCTGCAGGAGTCGGGCCCAGGACTGG<br>TGAAGCCTTCGGAGACCCTGTCCCTCACCTGCGC<br>TGTCTCTGGTGGCTCCATCAGCAGTAACTACTGG<br>AGCTGGATCCGCCAGCCCCCAGGGAAGGGGCTGG<br>AGTGGATTGGACGTATCTATGGTAGTGGTGGGAG<br>CACCGACTACAACCCCTCCCTCAAGAGTCGAGTC<br>ACCATTTCAACAGACACGTCCAAGAACCAGTTCT<br>CCCTGAAGCTGAGCTCTGTGACCGCCGCGGACAC<br>GGCCGTGTATTACTGTGCGAGAGA |
| HV | IGHV4-NL_33*01_S4467 | | | CAGGTGCAGCTGCAGGAGTCGGGCCCAGGACTGG<br>TGAAGCCTTCAGAGACCCTGTCCCTCACCTGCGC<br>TGTCTCTGGTGGCTCTATCAGCAGTAGTAACTGG<br>TGGAGCTGGATCCGCCAGCCCCCAGGGAAAGGGC<br>TGGAGTGGATTGGGTATATCAGTGGTAGTAGTGG<br>TAGCACCTACTACAACCCCTCCCTCAAGAGTCGA<br>GTCACCATTTCAAAAGACACGTCCAAGAACCAGT<br>TCTCCCTGAAGCTGAGCTCTGTGACCGCCGCGGA<br>CACGGCCGTGTATTACTGTGCGAGAGA |
| HV | | | IGHV4-80*01 | CAGGTGCAGCTGCAGGAGTCGGGCCCAGGACTGG<br>TGAAGCCTTCGGAGACCCTGTCCCTCACCTGCGC<br>TGTCTCTGGTGCCTCCATCAGTAGTTACTGGTGG<br>AGCTGGATCCGCCAGCCCCCAGGGAAGGGACTGG<br>AGTGGATTGGGGAGATCAATGGTAATAGTGGTAG<br>CACCTACTACAACCCCTCCCTCAAGAGTCGAGTC<br>ACCATTTCAAAAGACGCGTCCAAGAACCAGTTCT<br>CCCTGAAGCTGAGCTCTGTGACCGCCGCGGACAC<br>GGCCGTGTATTACTGTGCGAGATA |
| HV | IGHV5-15*01_S2502 | | | GAGGTGCAGCTGGTGCAGTCTGGAGCAGAGGTGA<br>AAAGGCCCGGGGAGTCTCTGAAGATCTCCTGTAA<br>GACTTCTGGATACAGCTTTACCAGCTACTGGATC<br>AGCTGGGTGCGCCAGATGCCCGGAAAAGGCCTGG<br>AGTGGATGGGGGCGATTGATCCTAGTGATTCTGA<br>TACCAGATACAGCCCGTCCTTCCAAGGCCAGGTC<br>ACCATCTCAGCCGACAAGTCCATCAGCACCGCCT<br>ACCTGCAGTGGAGCAGCCTGAAGGCCTCGGACTC<br>CGCCACGTATTACTGTGCGAAAGA |
| HV | IGHV4-149*01_S1992 | | | CAGGTGCAGCTGCAGGAGTCGGGCCCAGGACTGG<br>TGAAGCCTTCGGAGACCCTGTCCCTCACCTGCGC<br>TGTCTCTGGTGGCTCCTTCAGCAGTTACTGGTGG<br>AGCTGGATCCGCCAGCCCCCAGGGAAGGGACTGG<br>AGTGGATTGGGGAGATCAATGGTAATAGTGGGAG<br>CACCAACTACAACCCCTCCCTCAAGAGTCGAGTC<br>ACCATTTCAAAAGACGCGTCCAAGAACCAGTTCT<br>CCCTGAAGCTGAGCTCTGTGACCGCCGCGGACAC<br>GGCCGTGTATTACTGTGCGAGAAA |
| HV | IGHV3-50*01 | | | GAGGTGCAGCTGGTGGAGTCTGGAGGAGGCTTGG<br>TTCAGCCTGGGGGGTCCCTGAGACTCTCCTGTGC<br>AGCCTCTGGATTCACCTTCAGTAGCTATGGCATG<br>CACTGGGTCCGCCAGGCTCCAGGGAAGGGGCTGG<br>AGTGGGTGGCAGTTATATCGTATGATGGAAGTAA<br>GAAATACTACGCAGACTCTGTGAAGGACCGATTC<br>ACCATCTCCAGAGACAATTCCAAGAACATGCTAT<br>ATCTTCAAATGAACAACCTGAAATTGGAGGACAC<br>GGCCGTGTATTACTGTGCGAGAGA |
| HD | IGHD3-15*01 | IGHD3-9*01 | IGHD3-9*01 | GTATTACGAGGATGATTACGGTTACTATTACACC |

| | | | | |
|---|---|---|---|---|
| HD | IGHD3-18*01 | IGHD3-26*01 | | GTATTACTATGGTAGTGGTTATTACACC |
| HD | IGHD6-34*01 | IGHD6-11*01 | IGHD6-13*01 | GGGTATAGCAGCTGGTCC |
| HJ | IGHJ4-3*01 | IGHJ4*01 | IGHJ4*01 | ACTACTTTGACTACTGGGGCCAGGGAGTCCTGGT CACCGTCTCCTCAG |
| HJ | IGHJ6-6*01 | IGHJ6*01 | IGHJ6*01 | ATTACTACGGTTTGGATTCCTGGGGCCAAGGGGT CGTCGTCACCGTCTCCTCAG |
| HJ | | IGHJ1*02 | IGHJ1*02 | GCTGAATACTTCGAGTTCTGGGGCCAGGGCGCCC TGGTCACCGTCTCCTCCG |
| HJ | IGHJ5-5*01 | IGHJ5-2*01 | IGHJ5-2*02 | ACAACTCATTGGATGTCTGGGGCCGGGGAGTTCT GGTCACCGTCTCCTCAG |
| HJ | IGHJ5-4*03 | IGHJ5-1*01 | IGHJ5-1*01 | ACAACCGGTTCGATGTCTGGGGCCCGGGAGTCCT GGTCACCGTCTCCTCAG |

*Bernat: *Immunity* **54**, 355-366.e4 (2021).
#Ramesh: *Front Immunol* **8**, 1407 (2017).
^IMGT: International ImMunoGeneTics Information System
Latest sequence at http://kimdb.gkhlab.se/datasets/

**Table S4. GenBank accession numbers**

**A. Nucleotide sequences of viral strains and *env* genes used in the study**

| HIV/SIV Strain* | Accession number | Reference |
|---|---|---|
| HIV-1 M CRF250 | MW507842 | https://pubmed.ncbi.nlm.nih.gov/33658341/ |
| CPZ.Ptt.LB7 (co) | DQ373064 (OP604642) | https://pubmed.ncbi.nlm.nih.gov/16728595/ |
| CPZ.Ptt.LB715 (co) | KP861923 (OP604672) | https://pubmed.ncbi.nlm.nih.gov/30718403/ |
| CPZ.Ptt.MB897 (co) | JN835461 (OP604644) | https://pubmed.ncbi.nlm.nih.gov/33771926/ |
| CPZ.Ptt.MB66 (co) | DQ373063 (OP604647) | https://pubmed.ncbi.nlm.nih.gov/16728595/ |
| CPZ.Ptt.CAM13 (co) | AY169968 (OP604649) | https://pubmed.ncbi.nlm.nih.gov/33771926/ |
| CPZ.Ptt.GAB2 (co) | AF382828 (OP604673) | https://pubmed.ncbi.nlm.nih.gov/33771926/ |
| CPZ.Ptt.MT145 (co) | JN835462 (OP604651) | https://pubmed.ncbi.nlm.nih.gov/33771926/ |
| CPZ.Ptt.EK505 (co) | JN835460 (OP604674) | https://pubmed.ncbi.nlm.nih.gov/33771926/ |
| HIV-1 N DJ131 | AY532635 | https://pubmed.ncbi.nlm.nih.gov/15320995/ |
| CPZ.Ptt.DP943.2 (co) | EF535993 (OP604653) | https://pubmed.ncbi.nlm.nih.gov/33771926/ |
| GOR.BPID1 (co) | KP004989 (OP604660) | https://pubmed.ncbi.nlm.nih.gov/33771926/ |
| HIV-1 P RBF168 | GU111555 | https://pubmed.ncbi.nlm.nih.gov/19648927/ |
| GOR.CP2139 (co) | FJ424866 (OP604665) | https://pubmed.ncbi.nlm.nih.gov/33771926/ |
| GOR.CP2135 (co) | FJ424863 (OP604675) | https://pubmed.ncbi.nlm.nih.gov/33771926/ |
| GOR.BQID2 (co) | KP004991 (OP604662) | https://pubmed.ncbi.nlm.nih.gov/33771926/ |
| HIV-1 O RBF206 | KY112585 | https://pubmed.ncbi.nlm.nih.gov/33771926/ |
| CPZ.Pts.TAN2 | DQ374657 | https://pubmed.ncbi.nlm.nih.gov/33771926/ |
| CPZ.Pts.TAN3 | EF394358 | https://pubmed.ncbi.nlm.nih.gov/17494082/ |
| CPZ.Pts.TAN1 (co) | EF394356 (EF451055) | https://pubmed.ncbi.nlm.nih.gov/30718403/ |
| CPZ.Pts.TAN10 | ON959378 | This study |
| CPZ.Pts.UG38 | JN091690 | https://pubmed.ncbi.nlm.nih.gov/21775446/ |
| CPZ.Pts.TAN13 (co) | JQ768416 (OP604676) | https://pubmed.ncbi.nlm.nih.gov/30718403/ |
| CPZ.Pts.BF1167 | JQ866001 | https://pubmed.ncbi.nlm.nih.gov/33771926/ |
| CPZ.Pts.ANT_Cot | ON933805 | This study |
| MUS.11GAB (co) | KF304708 (OP604669) | https://pubmed.ncbi.nlm.nih.gov/33771926/ |
| MUS.1085.1_54 | MG450752 | https://pubmed.ncbi.nlm.nih.gov/33771926/ |
| MUS.1085.4_12 | MG450754 | https://pubmed.ncbi.nlm.nih.gov/30718403/ |
| ASC.RT11 (co) | KJ461714 (OP604656) | https://pubmed.ncbi.nlm.nih.gov/33771926/ |
| SMM.E660 (co) | JQ864086 (OP604677) | https://pubmed.ncbi.nlm.nih.gov/22696650/ |
| SMM.92b (co) | KU182919 (OP604678) | https://pubmed.ncbi.nlm.nih.gov/33771926/ |
| SMM.FTq (co) | KU182920 (OP604679) | https://pubmed.ncbi.nlm.nih.gov/33771926/ |
| AGM.TAN1 (co) | U58991 (OP604680) | https://pubmed.ncbi.nlm.nih.gov/33771926/ |
| MND2.M14 (co) | AF328295 (OP604681) | https://pubmed.ncbi.nlm.nih.gov/11435589/ |
| LHO.7 (co) | AF075269 (OP604682) | https://pubmed.ncbi.nlm.nih.gov/33771926/ |
| WRC.98CI (co) | AM713177 (OP604683) | https://pubmed.ncbi.nlm.nih.gov/18922864/ |
| WRC.05GM (co) | AM937062 (OP604684) | https://pubmed.ncbi.nlm.nih.gov/33771926/ |
| SCIV.CAM13K | OP373447 | This study |
| SCIV.CAM13RRK | OP373448 | This study |
| SIVcpzMB897-EnvFS | OP414674 | This study |

*(co), the synthesized env gene is codon optimized with the corresponding Genbank accession number listed in parentheses.

**B. Heavy and light chain variable region sequences of human monoclonal antibodies used in the study**

| Monoclonal antibodies | Heavy chain variable (VDJ) region sequence* | Light chain variable (VJ) region sequence* |
|---|---|---|
| **V2-apex bNAb** | | |
| PG9 | GU272045 | GU272046 |
| PG16 | GU272043 | GU272044 |
| PGT145 | JN201910 | JN201927 |
| PGDM1400 | KP006370<br>QAQLVQSGPEVRKPGTSVKVSCKA<br>PGNTLKTYDLHWVRSVPGQGLQW<br>MGWISHEGDKKVIVERFKAKVTIDW<br>DRSTNTAYLQLSGLTSGDTAVYYCA<br>KGSKHRLRDYALYDDDGALNWAVD<br>VDYLSNLEFWGQGTAVTVSS | KP006383 |
| VRC26.25 | KT371100 | KT371101 |
| CH01 | JQ267523 | JQ267519 |
| BG1 | 5VVF (PDB)<br>AEQLVESGGGLVPPGRSLRLSCSAS<br>GFYFPDYAMAWVRQAPGQGLQWV<br>GFMRGWAYGGSAQFAAFAVGKFAI<br>SRDDGRNVVYLDVKNPTFEDTGVYF<br>CAREQRNKDYRYGQEGFGYSYGM<br>DVWGRGTTVVVSTA | 5VVF (PDB)<br>DIHMTQSPVSLSASVGDRVTITCRA<br>SHFIANYVNWYQQKPGKAPTLLIFE<br>SSTLQRGVPSRFSAYGDGTEFTLSI<br>NTLQPEDFASYICQQSHSPPVTFG<br>AGTRVDQKRTVAA |
| VRC38.01 | KY905214 | KY905228 |
| **Inferred V2-apex precursor** | | |
| PG9_RUA | KC417411 | KC417412 |
| PG16_RUA | KC417413 | KC417414 |
| CAP256-VRC26UCA | KJ134860 | KJ134863 |
| CH01_RUA | KC417409 | KC417410 |
| **Conformation sensitive non-neutralizing antibodies** | | |
| 697-D | KP278511<br>QVQLVQSGAEVKKPGSSVKVSCKA<br>SGGNFNTYTISWVRQAPGQGLEWM<br>GRIIPIFGIVNPAQKFPGRVTINVDKS<br>TNTAYMELSSLRSEDTAVYYCATSG<br>VGLHFGYFDYWGQGTLVTVSS | KP278557 |
| 1393A | KP278509 | KP278555 |
| CH58 | KC417393 | KC417394 |
| CAP228-D3 | MK119170 | MK119171 |
| 3074 | EU794431 | KP278528 |
| 447-52D | KF924247 | KF924246 |
| 17b | 2NXY (PDB)<br>EVQLVESGAEVKKPGSSVKVSCKAS<br>GDTFIRYSFTWVRQAPGQGLEWMG<br>RIITILDVAHYAPHLQGRVTITADKST<br>STVYLELRNLRSDDTAVYFCAGVYE<br>GEADEGEYDNNGFLKHWGQGTLVT<br>VSSAS | 2NXY (PDB)<br>DIVMTQSPATLSVSPGERATLSCRA<br>SESVSSDLAWYQQKPGQAPRLLIY<br>GASTRATGVPARFSGSGSGAEFTL<br>TISSLQSEDFAVYYCQQYNNWPPR<br>YTFGQGTRLEIKR |
| A32 | 4YBL (PDB) | 4YBL (PDB) |

| | | |
|---|---|---|
| | QVQLQESGPGLVKPSQTLSLSCTVS GGSSSSGAHYWSWIRQYPGKGLE WIGYIHYSGNTYYNPSLKSRITISQH TSENQFSLKLNSVTVADTAVYYCAR GTRLRTLRNAFDIWGQGTRVTVSSA S | ALTQPPSASGSPGQSVTISCTGTSS DVGGYNYVSWYQHHPGKAPKLIIS EVNNRPSGVPDRFSGSKSGNTASL TVSGLQAEDEAEYYCSSYTDIHNFV FGGGTKLTVLGQPKAA |

*Heavy and light chain variable region sequences are listed as GenBank accession numbers when available expression plasmids matched these reference sequences or are listed as amino acid sequences when available expression plasmids exhibited differences (highlighted in red), or when only Protein Data Bank (PDB) or no reference sequences were available.

**C. Nucleotide sequences of mutant SIV *env* genes used in the study**

| SIV Strain | Accession number |
|---|---|
| SIVcpzLB7 Q171K (co) | OP604643 |
| SIVcpzMB897 Q171K (co) | OP604645 |
| SIVcpzMB897 Q171K + N130K (co) | OP604646 |
| SIVcpzMB66 Q171K (co) | OP604648 |
| SIVcpzCAM13 Q171K (co) | OP604650 |
| SIVcpzMT145 Q171K (co) | OP604652 |
| SIVcpzDP943.2 Q171K (co) | OP604654 |
| SIVcpzDP943.2 Q171K + N156 glycan (K156N+F158S) (co) | OP604655 |
| SIVgorBPID1 E170K + Q171K (co) | OP604661 |
| SIVgorCP2139 T169K + T170K (co) | OP604667 |
| SIVgorCP2139 T169K + T170K + K166R (co) | OP604666 |
| SIVgorCP2139 T169K + T170K + K166R + N130K (co) | OP604685 |
| SIVgorBQID2 S170K + Q171K (co) | OP604663 |
| SIVgorBQID2 S170K + Q171K + N130T (co) | OP604664 |
| SIVcpzTAN10 Q171K (wt) | OP604671 |
| SIVmus11GAB Q171K (co) | OP604670 |
| SIVmus1085 4-12 Q171K (wt) | OP604668 |
| SIVascRT11 H171K (co) | OP604658 |
| SIVascRT11 H171K + K166R (co) | OP604659 |
| SIVascRT11 H171K + A170K (co) | OP604657 |

*co, the synthesized *env* gene is codon optimized

**D. Longitudinal SGA sequences of SCIV.CAM13K and SCIV.CAM13RRK env sequences are available under GenBank accession numbers XXX-YYY.**

**E. Nucleotide sequences of heavy and light chain variable regions of RM monoclonal antibodies**

| Heavy (H) chain variable (VDJ) region | Accession number | Kappa (K) or lambda (L) light chain variable (VJ) region | Accession number |
|---|---|---|---|
| T925 w24 P1E09H | OP081691 | T925 w24 P1E09K | OP081721 |
| T925 w24 P1D09H | OP081692 | T925 w24 P1D09K | OP081722 |
| T925 w24 P2B11H | OP081690 | T925 w24 P2B11K | OP081720 |
| T925 w24 P1A12H | OP081694 | T925 w24 P1A12K | OP081724 |
| T925 w24 P1C08H | OP081693 | T925 w24 P1C08K | OP081723 |
| T925 w24 P1G03H | OP081698 | T925 w24 P1G03L | OP081728 |

| | | | |
|---|---|---|---|
| T925 w24 P1E11H | OP081695 | T925 w24 P1E11L | OP081725 |
| T925 w24 P1F01H | OP081696 | T925 w24 P1F01L | OP081726 |
| T925 w24 P2C12H | OP081699 | T925 w24 P2C12L | OP081729 |
| T925 w24 P1F12H | OP081697 | T925 w24 P1F12L | OP081727 |
| T927 w32 P2D12H | OP081707 | T927 w32 P2D12L | OP081737 |
| T927 w24 P1D08H | OP081703 | T927 w24 P1D08L | OP081733 |
| T927 w24 P2D06H | OP081706 | T927 w24 P2D06L | OP081736 |
| T927 w24 P1D09H | OP081704 | T927 w24 P1D09L | OP081734 |
| T927 w62 P1C05H | OP081712 | T927 w62 P1C05L | OP081742 |
| T927 w62 P1C01H | OP081711 | T927 w62 P1C01L | OP081741 |
| T927 w62 P1B01H | OP081708 | T927 w62 P1B01L | OP081738 |
| T927 w62 P1B10H | OP081710 | T927 w62 P1B10L | OP081740 |
| T927 w62 P3E08H | OP081718 | T927 w62 P3E08L | OP081748 |
| T927 w62 P1H10H | OP081715 | T927 w62 P1H10L | OP081745 |
| T927 w62 P1F05H | OP081714 | T927 w62 P1F05L | OP081744 |
| T927 w62 P3C02H | OP081717 | T927 w62 P3C02L | OP081747 |
| T927 w62 P1C07H | OP081713 | T927 w62 P1C07L | OP081743 |
| T927 w62 P3A11H | OP081716 | T927 w62 P3A11L | OP081746 |
| T927 w24 P2B12H | OP081705 | T927 w24 P2B12K | OP081735 |
| T927 w24 P1C02H | OP081701 | T927 w24 P1C02K | OP081731 |
| T927 w24 P1D03H | OP081702 | T927 w24 P1D03K | OP081732 |
| T927 w24 P1A11H | OP081700 | T927 w24 P1A11K | OP081730 |
| T927 w62 P1B05H | OP081709 | T927 w62 P1B05K | OP081739 |
| T927 w62 P3G11H | OP081719 | T927 w62 P3G11K | OP081749 |

Shading identifies different expanded lineages.
